# Structures of active melanocortin-4 receptor−Gs-protein complexes with NDP-α-MSH and setmelanotide

**DOI:** 10.1101/2021.04.22.440868

**Authors:** Nicolas A. Heyder, Gunnar Kleinau, David Speck, Andrea Schmidt, Sarah Paisdzior, Michal Szczepek, Brian Bauer, Anja Koch, Monique Gallandi, Dennis Kwiatkowski, Jörg Bürger, Thorsten Mielke, Annette Beck-Sickinger, Peter W. Hildebrand, Christian M. T. Spahn, Daniel Hilger, Magdalena Schacherl, Heike Biebermann, Tarek Hilal, Peter Kühnen, Brian K. Kobilka, Patrick Scheerer

## Abstract

The melanocortin-4 receptor (MC4R), a hypothalamic master regulator of energy homeostasis and appetite, is a G-protein coupled receptor and a prime target for the treatment of obesity. Here, we present cryo-electron microscopy structures of MC4R− Gs-protein complexes with two recently FDA-approved drugs, the peptide agonists NDP-α-MSH and setmelanotide, with 2.9 Å and 2.6 Å resolution. Together with signaling data, the complex structures reveal the agonist-induced origin of transmembrane helix (TM) 6 regulated receptor activation. In both structures, different ligand binding modes of NDP-α-MSH, a high-affinity variant of the endogenous agonist, and setmelanotide, an anti-obesity drug with biased signaling, underline the key role of TM3 for ligand-specific interactions and of calcium ion as a ligand-adaptable cofactor. The agonist-TM3 interplay subsequently impacts the receptor− Gs-protein interfaces, mainly at intracellular loop 2. These structures reveal mechanistic details of MC4R activation or inhibition and provide important insights into receptor selectivity that will facilitate the development of tailored anti-obesity drugs.

## INTRODUCTION

The melanocortin-4 receptor (MC4R) is one of five melanocortin receptor subtypes (MC1-5R) that share a set of similar peptidic ligands and constitute an evolutionarily related group of class A G-protein-coupled receptors (GPCRs). MCRs regulate energy homeostasis, pigmentation, cardiovascular function, and sexual functions (Wikberg and Mutulis, 2008). In particular, the MC4R plays a central role in energy balance and appetite regulation (Krashes et al., 2016). Naturally-occurring human MC4R variants are the most frequent monogenic cause of obesity, with over 160 identified mutants (Heyder et al., 2019).

Activation of MC4R by its natural agonists α-melanocyte-stimulating hormone (α-MSH) or β-MSH leads to appetite-reducing effects. In contrast, binding of the endogenous inverse agonist agouti-related peptide (AgRP) causes orexigenic effects (Biebermann et al., 2012) by reducing high levels of basal signaling activity (Nijenhuis et al., 2001).

Besides stimulation of heterotrimeric G_s_αβγ protein (Gs), MC4R can elicit other signaling pathways associated with the recruitment of Gq, Gi, or arrestin (Tao, 2010).

To date, pharmacological approaches targeting MC4R have failed mostly due to the severe adverse effects of drug candidates caused for instance by a lack of MCR subtype selectivity or G-protein pathway specificity.

Setmelanotide (also termed RM-493, BIM-22493 or IRC-022493) (Chen et al., 2015; Clement et al., 2020; Kuhnen et al., 2016) is the first FDA-approved (2020) medication (brand name *Imcivree*) for the treatment of rare genetic conditions resulting in obesity, including pro-opiomelanocortin deficiency (POMC), proprotein subtilisin/kexin type 1 deficiency (PCSK1), and leptin receptor deficiency (LEPR) (Kuhnen et al., 2020). Setmelanotide is a cyclic high-affinity peptide with a G-protein signaling profile biased towards Gq (Clement et al., 2018). Further, it has a 20-fold subtype selectivity towards MC4R when compared with α-MSH (Kumar et al., 2009). In contrast to other MC4R agonists, setmelanotide does not cause common Gs-signaling related adverse effects, such as tachycardia or hypertension. Although rare, moderate adverse effects, such as skin hyperpigmentation, have been reported.

Another FDA approved (2019), synthetic but non-selective MCR agonist, NDP-α-MSH (also termed afamelanotide; brand name *Scenesse*) is a linear high-affinity analog of α-MSH (Sawyer et al., 1980) for treatment of MC1R-driven melanogenesis, thereby preventing skin damage from sun exposure (phototoxicity) in individuals with erythropoietic protoporphyria. Moderate adverse effects of NDP-α-MSH exposure include headache, nasopharyngitis, or back pain (Langendonk et al., 2015). The numerous adverse effects of known MCR ligands call for the discovery of more selective ligands for specific MCR subtypes, in particular the MC4R, as the global increasing prevalence of human obesity is a growing medical and socioeconomic problem (Ericson et al., 2017).

Accordingly, a comprehensive understanding of MC4R-mediated signaling regulation is of fundamental importance. To explore the structural basis of agonist action and receptor-mediated signaling, we determined two cryo-electron microscopy (cryo-EM) structures of human wild-type MC4R− Gs-protein complexes bound to agonists setmelanotide and NDP-α-MSH, with resolutions of 2.6 Å and 2.9 Å, respectively. Comparison of our active structures with the recently solved MC4R crystal structure bound to an antagonist (Yu et al., 2020) demonstrate the essential role of calcium ion in forming a link between ligands and TM2 and TM3. In addition, the allosteric connection between the peptide agonist binding pocket (LBP) and the G-protein binding cavity (GBC) is mediated by TM3 and facilitated by TM6. Our structural insights combined with signaling data from site-directed mutagenesis reveal mechanistic details of receptor activation and inhibition.

## RESULTS

### MC4R− Gs complex formation with agonists NDP-α-MSH and setmelanotide

To determine the cryo-EM structure of MC4R-signaling complexes with different agonists, we used NDP-α-MSH and setmelanotide due to their clinical significance and high *in vitro* binding potency (K_i_ = 0.7 nM (Yu *et al*., 2020) and 2.1 nM (Kumar *et al*., 2009), respectively) compared with α-MSH (K_i_ = 51 nM (Yu *et al*., 2020)), which was essential for generating stable complex samples. Both ligands differ in their general peptide structure. NDP-α-MSH is a linear 13-amino acid peptide and is only modified at two positions (methionine (M4) to norleucine (Nle) and L-phenylalanine (L-F) to D-phenylalanine (D-F)) compared to α-MSH (Sawyer *et al*., 1980) (Figure 1A). In contrast, setmelanotide is a cyclic 8-amino acid peptide, which resembles α-MSH at only three positions, in the central *H*^*0*^*x*^*1*^*R*^*2*^*W*^*3*^ core motif (Figure 1A) [An unifying MCR ligand peptide numbering scheme based on the conserved *H*^*0*^*x*^*1*^*R*^*2*^*W*^*3*^ motif is introduced in Figure 1A and is used throughout the text.].

**Figure 1:**
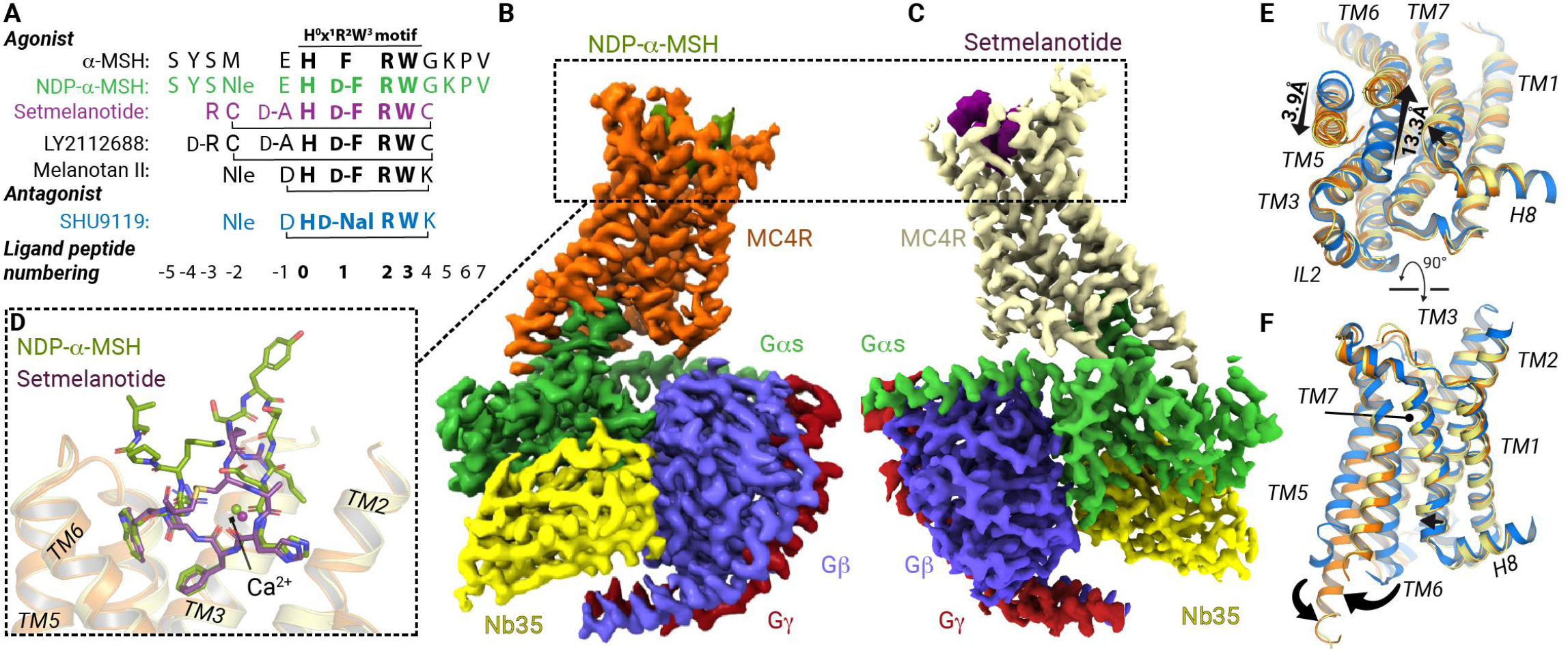
Cryo-EM complex structures of MC4R− Gs bound with NDP-α-MSH and setmelanotide. **(A)** Sequence alignment of MC4R agonists and antagonist SHU9119. For comparison and simplification, a ligand unifying numbering system is introduced based on the α-MSH core *HxRW* motif and will be indicated in superscript for each peptide residuesD. (**B− C)** Cryo-EM densities of MC4R− Gs complexes stabilized by Nb35 and bound agonists, displayed from mirrored perspectives; Gβ, dark blue; Gγ, red; Nb35, yellow. (**D)** Superposition of both agonists. (**E− F)** Comparison between SHU9119-antagonized (blue) MC4R and the active state structures, (**E)** from the cytoplasmic view, (**F)** from inside the membrane plane. Arrows indicate significant relative spatial transmembrane helix differences (TM5, TM6 and TM7).

We confirmed the binding potency of both ligands using a nano-luciferase-based BRET assay (Figure S1) (Stoddart et al., 2015). With our established workflow using *Sf*9 insect cells, we produced the human wild-type MC4R without modifications in sufficient protein amounts for assembling a complex with the Gs-protein (Figure S2). There was a strong tendency for the MC4R to oligomerize after detergent-based extraction from membranes. This oligomer population was reduced by direct formation of the complex with detergent-purified Gs-protein (Rasmussen et al., 2011) and MC4R that remained membrane-embedded (Zhang et al., 2017). After stabilization of the complex through the addition of nanobody-35 (Nb35) (Rasmussen *et al*., 2011) and nucleotide hydrolase apyrase, the resulting nucleotide-free agonist-bound MC4R− Gs− Nb35 complexes were extracted from membranes by solubilization and purified (Figure S2). The stable complexes with either agonists NDP-α-MSH or setmelanotide were used for cryo-EM grid preparation and single-particle analysis, yielding cryo-EM maps with a global resolution of 2.9 Å and 2.6 Å, respectively (Figures 1B and 1C; Figure S3− 6; Table S1; Methods).

### Overall structure of MC4R− Gs complexes bound with agonists NDP-α-MSH and setmelanotide

The cryo-EM maps revealed well-defined densities that facilitate unambiguous modeling of the secondary structure and side chain orientations of the MC4R− Gs complex as well as the cofactor ion calcium (Ca^2+^), several water molecules, and the agonist peptides NDP-α-MSH and setmelanotide bound to the orthosteric LBP (Figures 1B− 1D, Figures S7 and S8).

Only a few components known for their high flexibility, such as the receptor N-(Ntt) and C-termini (Ctt), the intracellular loop (IL) 1, IL3, and the extracellular loop (EL) 1, as well as the alpha-helical domain of Gαs, were not fully built into the final map (modeled residues are listed in suppl. information *Methods*).

Both agonists are bound foremost with the *H*^*0*^*x*^*1*^*R*^*2*^*W*^*3*^ motif between the ELs and within the transmembrane bundle of the receptor (Figure 1D). The ligands engage MC4R through extensive van der Waals, hydrophobic, and polar interactions, with residues in the transmembrane helices (TMs) as well as EL2 (Figures S9− S11; Tables S2− S6). Specific structural MC4R characteristics result in a wide-opened extracellular vestibule of the LBP for large peptidic ligands (Figures 2A− 2C).

**Figure 2:**
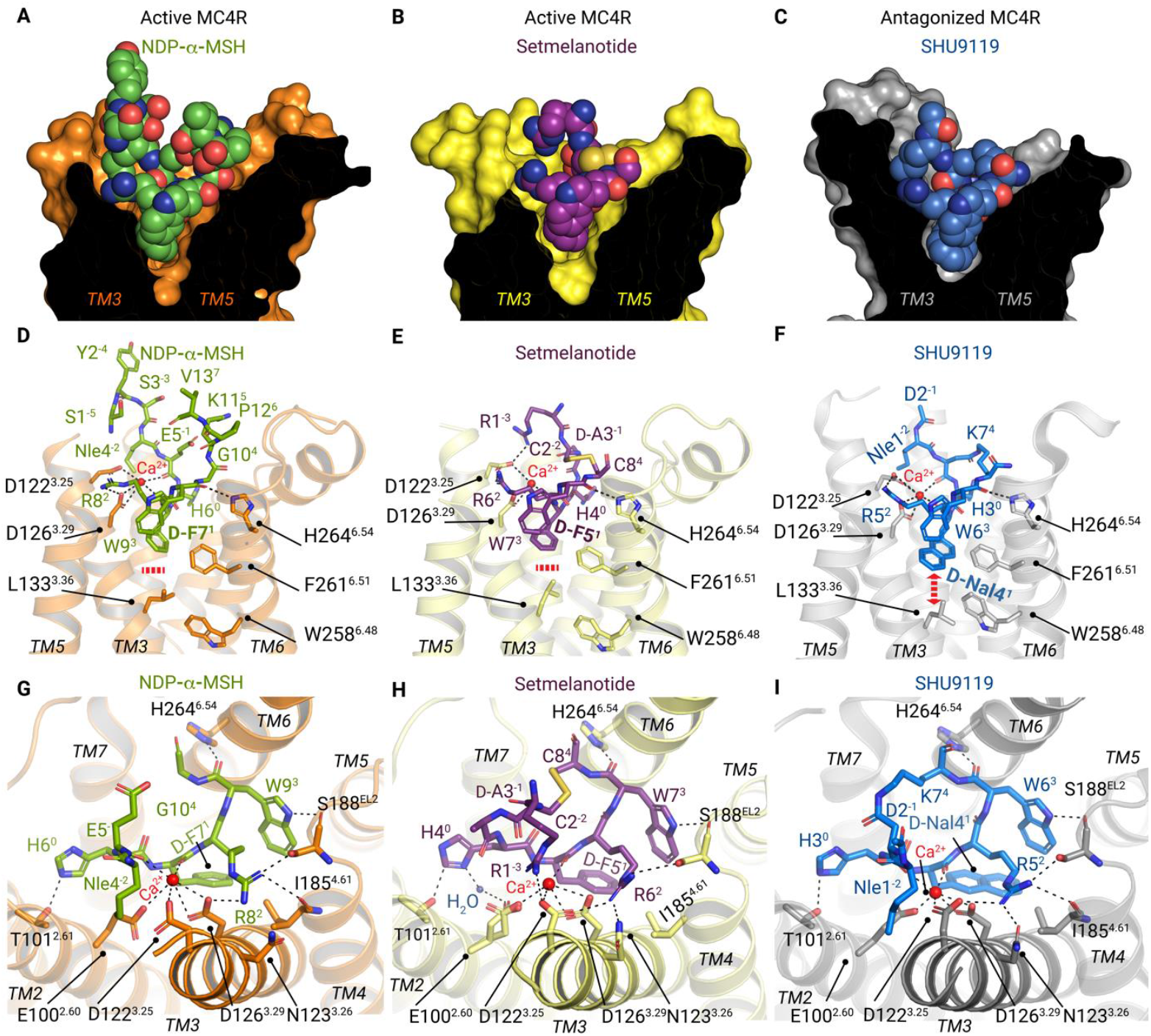
Binding modes of peptidic ligands and calcium at MC4R. **(A− C)** Sphere representations of NDP-α-MSH (carbon atoms; green), setmelanotide (carbon atoms, magenta), and SHU9119 (blue carbon atoms) bound into their respective MC4R binding sites (clipped surface, orange, light-yellow, gray, respectively). (**D− F)** TM3 and TM6 receptor residues involved in peptide and/or Ca^2+^ interactions are highlighted. (**D− E)** The bold red dashed red line indicates no interactions of L133^3.36^ with agonist residue D-F^1^; in (**F)** D-Nal4^1^ of the antagonist SHU9119 interacts with L133^3.36^ and modifies the L133 side chain orientation (red arrow). (**G-I)** Top view of (**D− E)**, highlighting all intermolecular hydrophilic interactions, assessed by a minimum distance of 3.5 Å (black dashed lines) (Figures S9-S11, Tables S2-S5). (**G)** NDP-α-MSH residues 4-11 are shown.

First, the EL2 is only four amino acids long and does not include a cysteine, significantly different from most other class A GPCRs (Woolley and Conner, 2017) (Figure S12). Consequently, the highly conserved disulfide bridge between EL2 and TM3 (Woolley and Conner, 2017) is absent. Instead of C^3.25^ [superscripted numbers according to the unifying Ballesteros & Weinstein numbering for class A GPCRs (Ballesteros and Weinstein, 1995)] in TM3, MC4R has an aspartate (D122^3.25^), which forms part of a ligand and Ca^2+^-binding network (Figures 2D− 2I).

Second, a disulfide bridge between C271^EL3^-C277^EL3^ in EL3 causes a specific helical conformation of the EL3− TM7 transition, which has been so far only structurally described for two lipidic evolutionary related GPCRs (Figures S13A and S13B). A second observed MC4R disulfide bridge between C279^7.30^ (in EL3) and C40^Ntt^ in the N-terminus (Figure S13C) is in agreement with previous functional studies (Chai et al., 2005) and stabilizes the EL3-Ntt interplay, thereby contributing to the formation of the extracellular LBP.

Third, in contrast to most other class A GPCRs, the MC4R has no proline in TM2 (Figures S12B; Figure S13D), nor the highly conserved proline P^5.50^ in TM5 (Figure S12C; Figure S14), which usually causes a kink and bulge, but also a slight rotation of the TMs. This is exemplified by a ∼6Å shift of the extracellular TM2 towards the membrane compared to other known structures, and a straight MC4R TM5 that shows significant differences to GPCRs with a proline (Figure S14), finally modifying the TM5− TM3 interface and together forming the MC4R ligand binding vestibule.

In addition, the MC4R− ligand complexes unravel the previously known contribution of Ca^2+^ as a cofactor (Kopanchuk et al., 2005; Salomon, 1990) and offers novel insights of its impact on the respective ligand-binding modes (Figures 2D− 2I; Figure 3).

**Figure 3:**
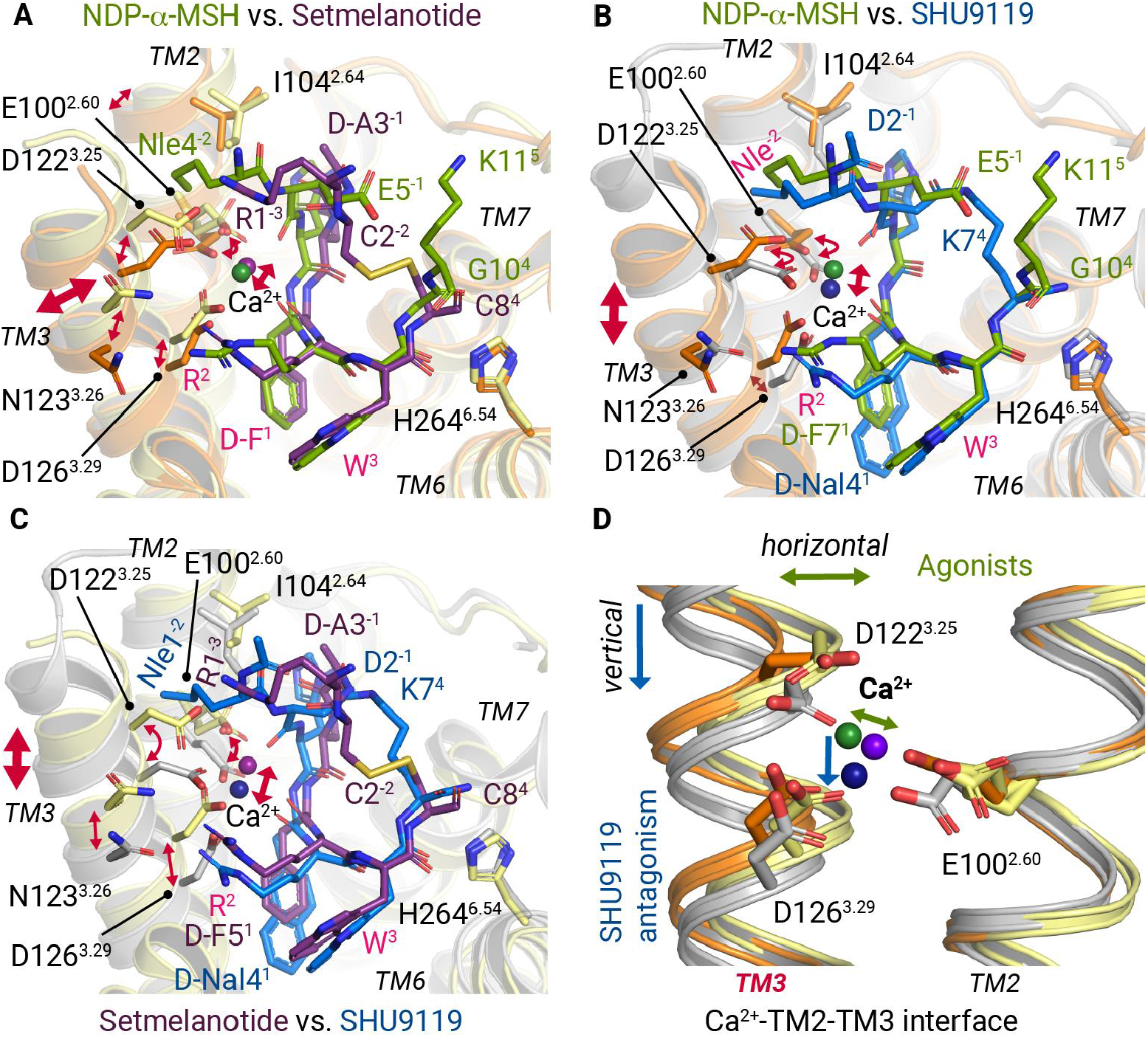
Differential Ca^2+^ and ligand binding at TM3. Superpositions of NDP-α-MSH (residues 4-11), setmelanotide, and SHU9119 bound at MC4R (orange, light-yellow, and gray backbone cartoons). **(A)** Superposition of both agonists. Residues that are shared by the agonists are labeled in pink. **(B)** NDP-α-MSH binding compared with SHU9119, and in **(C)** setmelanotide compared with SHU9119 is represented. Bi-directional red arrows indicating differences in the relative spatial positioning of residues, the TM3, or the Ca^2+^ ion. Detailed intermolecular interactions are summarized in Tables S5 and S6. **(D)** The relative alterations of TM3 orientation are also found in Ca^2+^ and its interacting residues of the *EDD* motif.

Both the wide extracellular vestibule and the cofactor calcium define a unique LBP, which is adapted to integrate signals from ligands with differences in size, as well as inducing specific cellular responses via different G-protein signaling pathways.

Agonist binding in the receptor LBP induces local reorientations in the highly conserved amino acid motifs *CWxP*^*6*.*50*^ (C257^6.47^-W258^6.48^-P260^6.50^, in MC4R-TM6), *N(D)P*^*7*.*50*^*xxY* (D298^7.49^-P299^7.50^-Y302^7.53^ in MC4R, in MC4R-TM7), and *DR*^*3*.*50*^*Y* (D146^3.49^-R147^3.50^-Y148^3.51^, in MC4R-TM3), which are associated with global helical movements in the receptor structure towards an active state, enabled for G-protein binding.

These helical movements are mainly an inward twisting of TM5 and an ∼13Å outward motion of TM6, a hallmark of receptor activation to open the binding cavity for G-proteins (Figures 1E and 1F).

### Similarities in agonist binding at MC4R

Comparison of the two MC4R structures bound to the agonists NDP-α-MSH and setmelanotide and the previously published MC4R structure in complex with the antagonist SHU9119 (Yu *et al*., 2020) shows, that all ligands are buried deeply in the extracellular receptor region between the TMs, but with differently shaped LBPs (Figures 2A− 2C). These unique shapes are caused by varying ligand residues (Figure 1A) and the fact that SHU9119 and setmelanotide are cyclic and more compact in contrast to the linear peptide NDP-α-MSH (Figure 1D and Figure 2).

All three ligands share a common central amino acid motif *H*^*0*^*x*^*1*^*R*^*2*^*W*^*3*^. This four-finger-like motif is essential for the formation of most relevant interactions within the LBP. The x^1^ position is always located at the bottom-part of the LBP (Figures 2D− 2F). NDP-α-MSH and setmelanotide have the stereoisomer D-phenylalanine (F^1^) instead of an L-F at position x^1^. This substitution is important for the increased potency of NDP-α-MSH compared with α-MSH (Sawyer *et al*., 1980).

The *H*^*0*^*x*^*1*^*R*^*2*^*W*^*3*^ motif forms similar interactions for both agonists, including hydrogen bonds between H^0^ and T101^2.61^ in TM2, and W^3^ with S188^EL2^ in EL2 and H264^6.54^ in TM6 (Figures 2G− 2H). The ligands also share several similar hydrophobic contacts, such as H^0^ to F284^7.35^-L288^7.39^-F51^1.39^, or D-F^1^ to I129^3.32^, and W^3^ to Y268^6.58^-I194^5.40^-L197^5.43^ (Figures S9 and S10; complete list in Table S6).

In both agonist-bound structures a D-F^1^ at position x^1^ stabilizes Ca^2+^ through a main chain interaction (Figure 1A; Figures 2G− 2H), while the side chain points into the core of the receptor and is surrounded by hydrophobic amino acids, namely C130^3.32^, L133^3.35^, I185^4.61^, L197^5.34^, F261^6.51^ and L288^7.39^. Functional characterization of substitutions at these positions and others in the extended LBP are provided in Figure 4, Figures S15-S16, Tables S7 and S8. The hydrophobic residues I291^7.42^ and L133^3.35^ are in close vicinity of W258^6.48^ of the highly conserved *CWxP*^*6*.*50*^ motif in TM6 that is usually in contact to ligands in several class A GPCRs (e.g. muscarinic acetylcholine receptor− G_11_ complex (Maeda et al., 2019), 5-HT2A serotonin receptor− G_q_ complex (Kim et al., 2020)). In contrast, for MC4R no direct interactions between the ligands and W258^6.48^ exist (Figures 2D− 2E). Overall, both agonists have a set of similar receptor contacts specifically to TM3 and TM6, although the interaction pattern with the cofactor calcium is in fact different.

**Figure 4:**
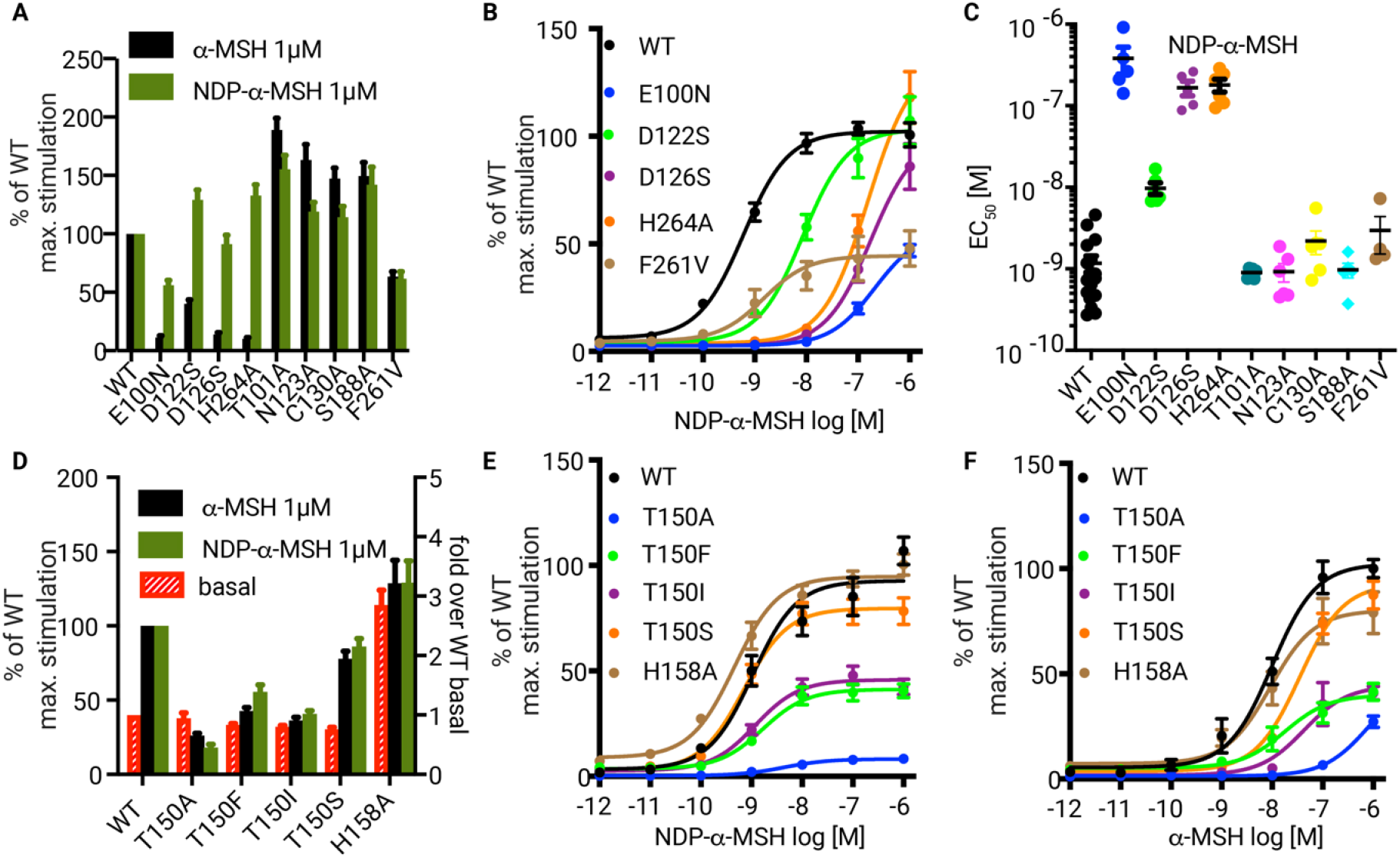
Functional data of amino acid substitutions of MC4R modified in the ligand- and G-protein binding regions. **(A-C)** cAMP signaling data for mutants in the ligand binding region and in the **(D-F)** G-protein interface are plotted. **(A and D**) Maximal Gs-protein signaling determined as cAMP accumulation of MC4R WT and mutants after addition of 1 µM α-MSH or NDP-α-MSH indicated as fold of MC4R WT max. signaling [%] (left y-axis). Basal cAMP accumulation of MC4R WT and mutants are depicted as fold over WT basal (right y-axis). (**B, E-F**) Concentration-response curves of Gs-protein signaling determined as cAMP accumulation of WT and indicated mutants after stimulation by NDP-α-MSH (**B, E**) and α-MSH (**F**) shown as fold change of WT maximum signaling [%]. (**C**) EC_50_ values [M] of NDP-α-MSH induced signaling calculated from concentration response curves in (B).

### Calcium is an essential and flexible link between the ligands and MC4R

In both agonist bound MC4R structures presented here, Ca^2+^ is attached centrally between and TM2 and TM3 and stabilizes peptide folding. In NDP-α-MSH, Ca^2+^ is five-fold coordinated by both main chain interactions with E5^-1^ and D-F7^1^ and receptor side chain interactions with E100^2.60^, D122^3.25^ and D126^3.29^ (Figure 2G). The latter three residues constitute the *EDD* motif in TM2 and TM3.

To verify the relevance of the ligand-interface between TM2 and TM3, we generated serine or asparagine mutations (retained hydrophilic side chains) of E100^2.60^, D122^3.25^, and D126^3.29^, and tested their ability to accumulate cAMP (Gs-induced signaling) by NDP-α-MSH or α-MSH stimulation (Figure 4; Tables S7 and S8). E100N and D126S substitutions resulted in a significant reduction of the NDP-α-MSH potency by two orders of magnitude compared to D122S with a modest EC_50_ reduction (9 nM instead of 1.2 nM for wild-type MC4R). Previous mutagenesis studies have already shown that substitution of D126A almost completely abolished and in the case of D122A and E100A, significantly reduced NDP-α-MSH-mediated signaling (Pogozheva et al., 2005; Yang et al., 2000). Of note, all three mutants nearly abolished endogenous α-MSH-signaling (Figure 4A; Tables S7 and S8), indicating a stronger impact of the *EDD* motif and associated Ca^2+^ binding on α-MSH action. This information is supported by the previous finding that α-MSH, unlike NDP-α-MSH, does not induce signaling at the D122A MC4R variant (Nickolls et al., 2003).

### Specificities of setmelanotide interactions with calcium and MC4R

Compared to NDP-α-MSH, setmelanotide forms unique interactions with Ca^2+^ and TM3 residues in MC4R. The setmelanotide specific R1^-3^ plays a key role beside R6^2^ and binds to D122^3.25^ (Figure 2H). Together, both residues R1^-3^ and R6^2^ constitute a tight arginine clamp in hydrogen bond distance to D122^3.25^ of the *EDD* motif, enabled by cyclization of the peptide. This arginine clamp has three particular consequences. Firstly, in contrast to the NDP-α-MSH complex, D122^3.25^ is fully oriented towards Ca^2+^, verified by the cryo-EM density map. Secondly, the side chain of R6^2^ shows a slightly different orientation toward TM3 by rotating away from TM4 as in the NDP-α-MSH− MC4R structure (Figures 2G− 2H). This is accompanied by a horizontal TM3 shift in the setmelanotide-MC4R complex and thus leads to a ligand-dependent Ca^2+^ positioning (Figures 3A and 3D). Thirdly, the setmelanotide interaction between R1^-3^ and the *EDD* motif reduces the number of interactions of the cofactor Ca^2+^ involved in stabilizing the peptide-TM2-TM3 interface by an only four-fold coordination of the ion. The reduced number of interactions results in a double-conformation of E100^2.60^, in which one conformation participates in a hydrogen bond network with a water molecule and H4^0^ (Figure 2H).

Of note, the MC4R agonist LY2112688 supports the relevance of R1^-3^ in setmelanotide. The only difference to setmelanotide is a D-R1^-3^ instead of L-R1^-3^ (Figure 1A) and setmelanotide is ∼50-fold more potent than LY2112688 (and ∼80 fold more potent than α-MSH) for inducing Gq-mediated signaling (Clement *et al*., 2018). This strongly supports that the specific pharmacological profile of setmelanotide originate from the observed unique interactions of R1^-3^ at the upper part of the LBP in TM3, which is the most significant difference compared to NDP-α-MSH.

### Essential features at TM6 requisite for the active MC4R state

In class A GPCRs a strong outward movement of TM6 (∼ 13Å in MC4R) toward the membrane is a hallmark of active state conformations, because it opens the intracellular cavity for G-protein binding (Figure 1E). This displacement is enabled and structurally apparent by an increase in the kink formation at the highly conserved *CWxP*^*6*.*50*^ motif (_*257*_*CWAP*_*260*_ in MC4R), which is essential for the previously proposed “toggle switch” activation mechanism in different GPCRs (Visiers et al., 2002). Comparison of the MC4R with known (in-)active class A GPCR structures revealed similar changes around the W258^6.48^ of the *CWxP*^*6*.*50*^ motif between the inactive or antagonized versus the active structures as observed e.g. in rhodopsin (Choe et al., 2011) and the 5-HT2A serotonin receptor (Kim *et al*., 2020) upon activation (Figure S17). Hence, we propose a toggle-like TM6 movement also during MC4R activation, which occurs around the *CWxP*^*6*.*50*^motif.

Two residues in MC4R TM6 are involved in ligand binding upstream of the *CWxP*^*6*.*50*^ motif, namely F261^6.51^ and H264^6.54^. H264^6.54^ forms a primary hydrogen bond interaction with the ligand’s backbone oxygen of W^3^ near the extracellular vestibule entrance (Figures 2G− 2I). Hence, the H264A mutant exhibits a significant decrease in potency (EC_50_) (Figures 4B-4C) and completely diminished signaling for the endogenous agonist α-MSH (Figure 4A).

In contrast, F261^6.51^ has only a weak interaction to the the x^1^ position of the ligand, but the F261V substitution showed a significant reduced cAMP accumulation (Figure 4A>-4B; Tables S7 and S8). This implies that the H264^6.54^− F261^6.51^ region acts as an initial agonist-dependent trigger, with H264^6.54^ directly contacting agonists, while F261^6.51^ may play an essential role in relaying the signal towards the helix-tilting TM6− *CWxP*^6.50^ region (Figure 5A, Figure 5C-5D). The MC4R− *CWxP*^6.50^ substitution W258A leads to a significant reduction but not a total loss in Gs signaling (Figure S16, and (Pogozheva *et al*., 2005)). This is accompanied by a reduced cell surface expression (Figures S16A), suggesting a loss of structural integrity of this variant. The W258F mutant, on the other hand, is expressed at the cell surface like wild-type, but with a higher basal receptor activity and a slightly reduced Gs signaling by ligand stimulation compared to the wild-type MC4R (Figure S16).

**Figure 5:**
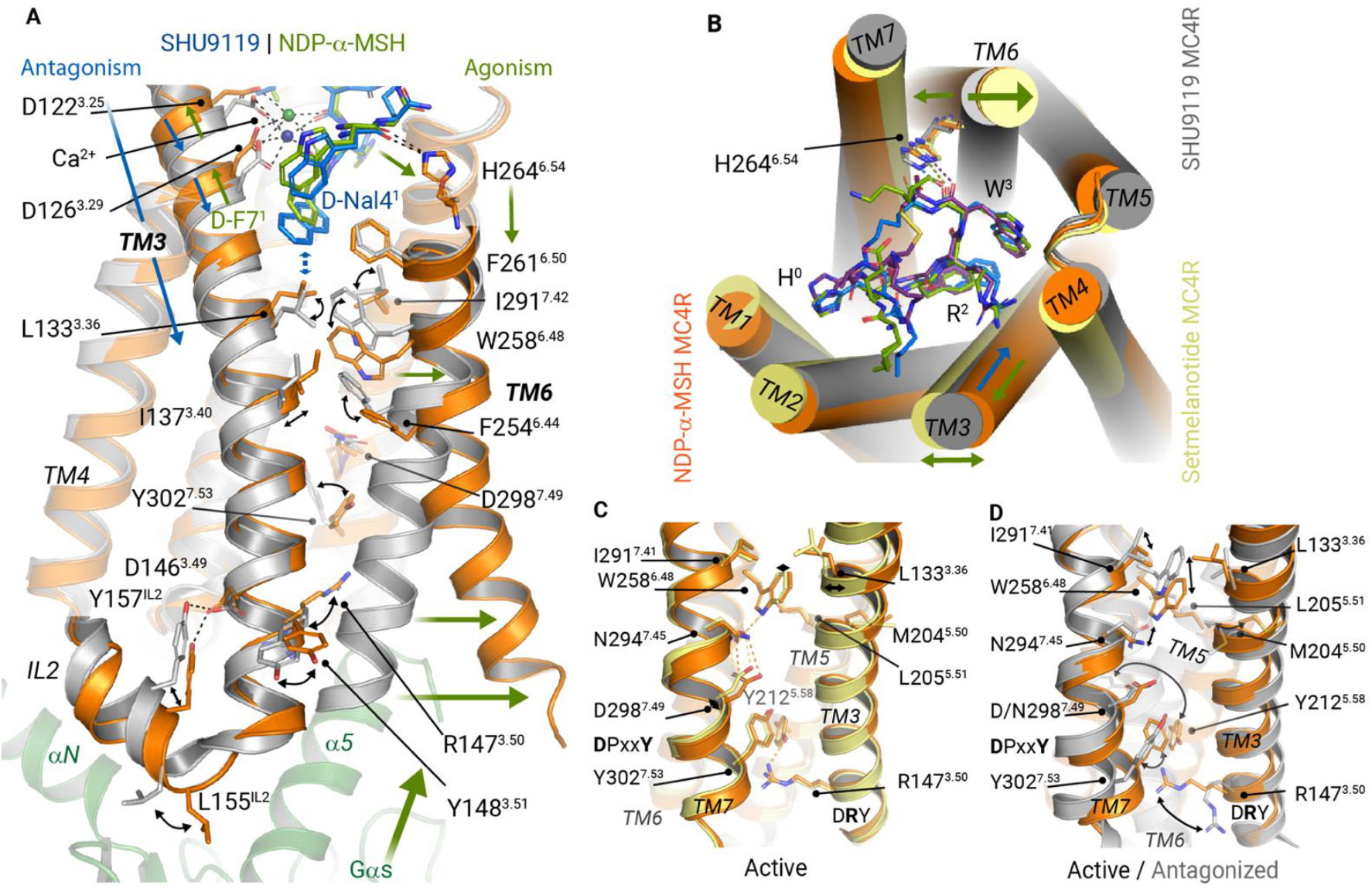
MC4R receptor activation is facilitated at TM6 and mediated at TM3. **(A)** Superposition of the agonist NDP-α-MSH (green/orange) or the antagonist SHU9119 (blue/gray) bound to MC4R. The supposed activation-pathway from the LBP to the Gs-protein interface is highlighted by green arrows and is triggered by ligand interactions at TM6, inducing the TM6 opening around W258^6.48^. Antagonistic action is facilitated by the interplay of D-Nal4^1^ with L133^3.36^ (indicated by blue arrows) and a subsequently blocked “toggle switch” at W258^6.48^. **(B)** Top view on ligand pockets highlights (arrows) the relative positioning of transmembrane helices TM1-6 (antagonized versus active structures). **(C− E)** Comparison of **(E)** active and (**D)** antagonized MC4R structures. (**E)** Superposition of active NDP-α-MSH− MC4R and SHU9119− MC4R structures. Relative movements, at the *CWxP*^*6*.*50*^, *P*^*5*.*50*^*(M)IF, N(D)P*^*7*.*50*^*xxY*, and *DR*^*3*.*50*^*Y* motifs, that accompany the receptor activation are highlighted by black arrows. Key residues involved in receptor activation are shown as sticks.

The importance of an aromatic residue at this position is further underlined by the fact that ∼70% of non-olfactory class A GPCRs contain a W^6.48^ and ∼16% a F^6.48^. Altogether, the W258^6.48^ is shifted between the antagonized and the active state structures (Figure S17) and is of significance for signaling regulation. However, this tryptophan is not the only key-switch for MC4R signal transduction in the transmembrane core, further amino acids contribute to receptor activation.

### Key sites for signal propagation at the transmembrane core

The MC4R− *CWxP*^*6*.*50*^ motif is spatially surrounded by an extended hydrophobic environment constituted by I137^3.40^, F201^5.47^, F254^6.44^, F261^6.51^, and I291^7.42^, of which only F254^6.44^ and I291^7.42^ display significant displacements comparing the antagonized with the agonized structures (Figure 5A). It is important to note that in most other class A GPCRs, only small amino acids such as alanine or glycine are present at position 7.42 (Figure S12E), and only a few examples such as the ghrelin or thyrotropin receptors have larger aromatic amino acids there. In MC4R, the mutant I291A is unable to induce cAMP signaling, while substitution of I291 by the larger hydrophobic phenylalanine side chain partially retained signaling (Figure S16). This observation together with the specific interaction of I291^7.42^ to W258^6.48^ observed in the agonist bound structures implies a hydrophobic interplay between positions 7.42 and 6.58 in the MC4R that is mandatory for regulation of MC4R activation.

In addition, F254^6.44^ (TM6) together with I137^3.40^ (TM3), and M204^5.50^ (TM5) constituting a *M*^*5*.*50*^*IF* motif in MC4R, reminiscent of the class A GPCR-typical *P*^*5*.*50*^*IF* motif (P^5.50^-I^3.40^-F^6.44^) involved in the maintenance of an inactive receptor state (Kobilka, 2011). Interestingly, all substitutions of residues in the MC4R-*M*^*5*.*50*^*IF* motif (I137A, I137F, F254A, F254M and M204A) do not affect agonist-induced signaling (Figure S18, Tables S7− S8). However, M204^5.50^ causes a straight, helical conformation of TM5, whereas most other class A GPCRs contain a P^5.50^ at this position (∼ 80% conserved) that induces a kink and bulge in the transmembrane helix (Yohannan et al., 2004) (Figure S14). The MC4R M204A substitution and the L205F mutant of the neighboring amino acid exhibiting significantly increased basal signaling compared to wild-type MC4R (Figure S18), which indicates that this TM5 segment is crucial for regulating basal signaling activity.

Despite the hydrophobic interaction with I291^7.42^ in TM7, W258^6.48^ is also connected in the active state via a hydrogen bond to N294^7.45^, which in turn is coupled to D298^7.49^ of the *N(D)P*^*7*.*50*^*xxY* motif in TM7 (Figure 5C). This interaction shifts the *N(D)P*^*7*.*50*^*xxY* motif slightly toward the GBC (Figure 5C), accompanied by a rotation of Y302^7.53^ in the direction of Y212^5.58^ in TM5. This orientates Y212^5.58^ into a position that allows stabilization of the active conformation of R147^3.50^ in the *DR*^*3*.*50*^*Y* motif, which forms part of the G-protein interface.

### Antagonizing the MC4R by impeding the active state formation

The antagonistic peptide SHU9119, recently co-crystallized with the antagonized MC4R (Yu *et al*., 2020), is a shortened, modified, and circularized derivative of NDP-α-MSH (Hruby et al., 1995) (Figure 1A). The binding mode of SHU9119 is generally very similar to that of NDP-α-MSH. Nearly identical hydrophobic contacts exist in the central *H*^*0*^*x*^*1*^*R*^*2*^*W*^*3*^ motif as in NDP-α-MSH (Figure S11, Table S4). W6^3^ is also clamped by H264^6.54^ (TM6) and S188^EL2^, and R5^2^ connects TM3-TM4 and EL2 through hydrogen bonds with S188^EL2^ and I185^4.61^ (Figure 2I; Figure S11, Table S4).

All three ligands, therefore, show a similar interaction pattern in the upper part of the LBP, including the participation of Ca^2+^ in the interaction with the *EDD* motif. The question arises of how SHU9119 antagonizes the MC4R?

This can only be explained by the action of the unnatural amino acid D-naphthylalanine (D-Nal4^1^) in SHU9119 at position x^1^ in the *H*^*0*^*x*^*1*^*R*^*2*^*W*^*3*^ motif that corresponds to D-F^1^ in NDP-α-MSH and setmelanotide (Figure 1A, Figures 2D− 2F). In contrast to the active state structures presented here, the bulky aromatic side chain of D-Nal4^1^ of SHU9119 pushes the side chain L133^3.36^ down, accompanied by a vertical downward shift in the upper half of TM3 of around 1 Å (Figure 5A). Thereby, the side chain of L133^3.36^ is moved in front of W258^6.48^, which eliminates the capacity for the TM6 outward movement (Figure 5A). This structural observation is functionally supported by the fact that a single mutation at L133^3.36^ to a more flexible and not bulky methionine side chain reverses SHU9119 to an agonist (Yang et al., 2002).

Of note, SHU9119 is an agonist for the MC1R and MC5R that have a natural M128^3.36^ or V126^3.36^ at this position (Figure S12B and Figure S19), respectively. Complementarily, melanotan II (Dorr et al., 1996) is almost identical to SHU9119 except that it contains a D-F^1^ instead of the D-Nal4^1^ and agonizes MC4R (Figure 1A). Our functional characterization of MC4R L133^3.36^ substitutions to either alanine or phenylalanine reveal no impact on NDP-α-MSH signaling, supporting the observation that the smaller D-F^1^ does not affect the position of L133^3.36^ (Tables S6 and S7; Figure S16).

In summary, SHU9119 shows a similar binding pattern in the upper part of the MC4R LBP compared to the agonists, but acts as an antagonist due to the D-Nal4^1^-L133^3.36^ interaction. This comparison supports the key role of the above discussed hydrophobic interplay of W258^6.48^ and I291^7.42^, which are captured in an inactive or antagonized state by the shifted L133^3.36^ (Figure 5A).

### The intracellular TM3 as a transducer of agonist-induced G-protein binding

In the antagonized MC4R structure, R147^3.50^ of the *DR*^*3*.*50*^*Y* motif in TM3 forms potential hydrogen bonds with the T150^3.53^ and D146^3.49^ side chains and with the backbone of N240^6.30^ in the IL3− TM6 transition (Figure 6A), which is in contrast to other inactive class A GPCR structures (e.g., β2AR, Figure 6C). In addition, the side chain of N240^6.30^ interacts through a hydrogen bond with the backbone oxygen at position T150^3.53^, which suggests an intracellular dual lock formed by hydrogen bonds between TM3 and TM6 that stabilizes the inactive state (Figure 6A). Furthermore, IL2 forms a short 3_10_-helix in the inactivated MC4R, which is associated by the interaction of D146^3.49^ with Y157^IL2-3.60^ in IL2.

**Figure 6:**
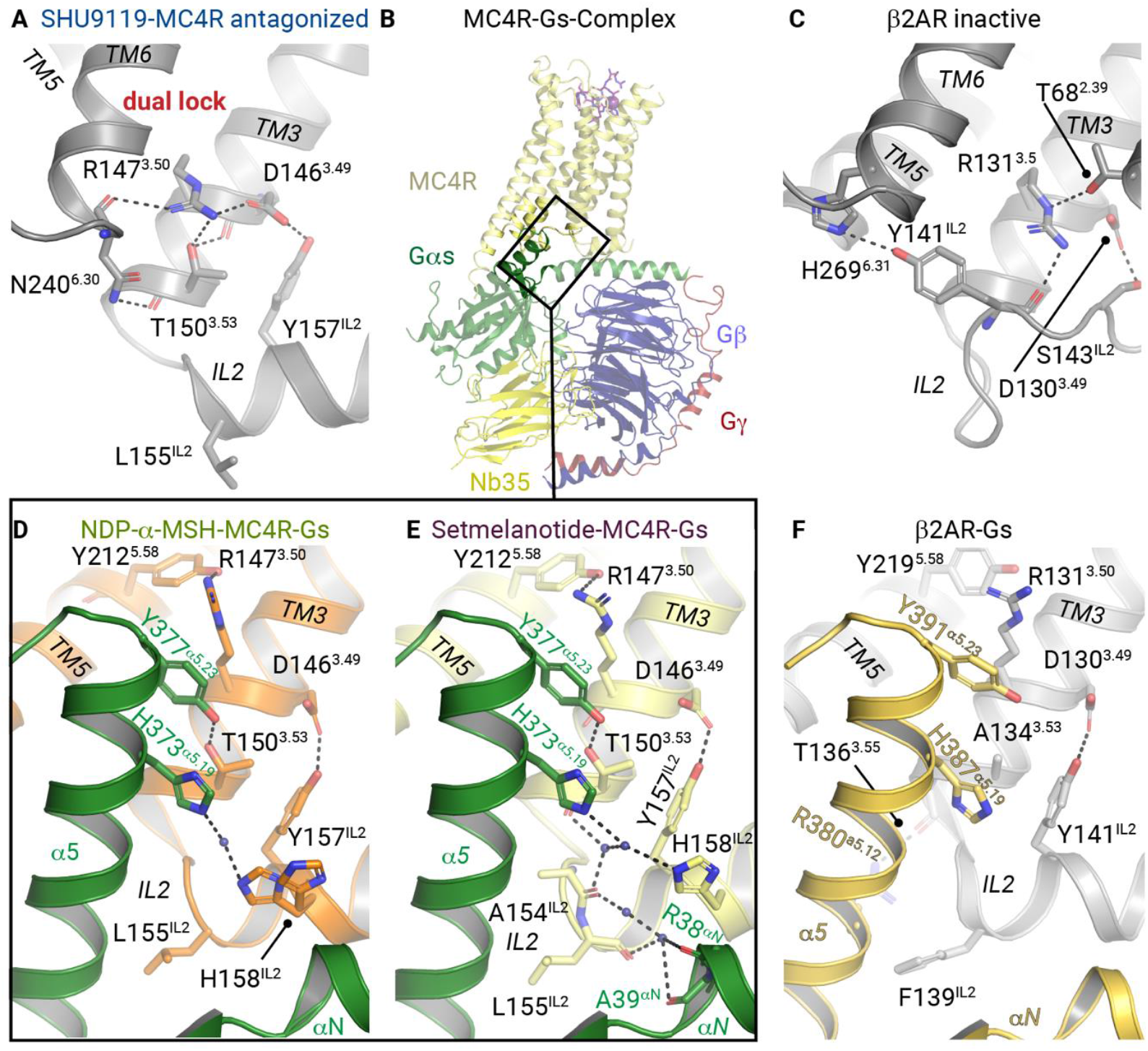
Intracellular interactions in the binding crevice between MC4R and Gs-protein. Upper panel: Intramolecular interactions at the IL2− TM3− TM6 site stabilizing (**A)** the antagonized MC4R (PDB ID: 6w25), and (**C)** the inactive β2AR (PDB ID: 2rh1). (**B)** Overall view on MC4R− G_s_ with enlarged areas in panel shown in (**D− E)**. Active receptor− Gs complex structures of MC4R bound to **(D)** NDP-α-MSH, **(E)** setmelanotide and **(F)** β2AR (PDB ID: 3sn6). Black dashed lines indicate hydrophilic interactions.

Comparing the active with the antagonized MC4R structure, receptor activation is strongly accompanied by *DR*^*3*.*50*^*Y* motif side chain rearrangements, considered as typical for class A GPCR activation (Figure 6D− 6F) (Park et al., 2008). MC4R− R147^3.50^ is stabilized by a hydrogen bond to Y212^5.58^ in TM5, constituting the G-protein cavity base. Extensive interactions in the interface between the activated MC4R and Gs-protein occur via the α5-helix of the Gαs domain and its C-terminal loop, termed C-cap, which has been already described for other GPCR− G-protein complexes (Rasmussen *et al*., 2011; Scheerer et al., 2008).

Following disruption of the MC4R TM3− TM6 dual lock by ligand interaction and the TM6 outward tilt (Figure 6A), both residues, R147^3.50^ and T150^3.53^ acquire a key role in constituting an active state conformation. R147^3.50^ forms a cation-π stacking interaction with the side chain of Y377^α5.23^ (CGN in superscript (Flock et al., 2015)) in the α5− C-cap.

The side chain of Y377^α5.23^ is further stabilized by a hydrogen bond to T150^3.53^ (Figures 6D− 6E), which was involved in stabilization of the inactive state. Such stabilization by an additional hydrogen bond interaction has not been identified in other GPCR− Gs complex structures known so far and highlights T150^3.53^ as a key site in the switching process between inactive and active MC4R conformations. A similar interaction has been observed only between S126^3.53^ of the muscarinic receptor M1 and Y356 ^α5.23^ from the G_11_-protein (Maeda *et al*., 2019). Of note, most class A GPCRs contain an alanine at the corresponding position of MC4R− T150^3.53^ (Figure S20). Our functional data from mutagenesis studies confirm the important role of this intermolecular polar interaction (Figure 4D-4F; Figure S21, Tables S7 and S8). T150^3.53^ substitutions by hydrophobic and acidic residues drastically reduces agonist-induced maximal cAMP accumulation. In contrast, the T150S mutant displays a comparable level of signaling for stimulation by NDP-α-MSH (Figure 4D-4E; Tables S7 and S8).

### Common and unique G-protein interactions with the IL2

One of the most important interactions between the IL2 of MC4R and Gs is formed by residues L155^IL2-3.58^ and F362^α5.08^ (Figures 7A and 7B). In the MC4R− Gs complex, the side chain of L155^IL2-3.58^ is located in the interface between the Gαs αN− β1 junction, the β2− β3 turn, and the α5-helix (Figures 7A and 7B). This IL2-β2/β3-α5 lock of the L155^IL2-3.58^ is reminiscent of the known F139^IL2-3.58^-F376^α5.08^ interplay in the β2AR− Gs complex (Figure 7C) or to Gq-coupling interactions at M1 and M3 receptors (Moro et al., 1993).

**Figure 7:**
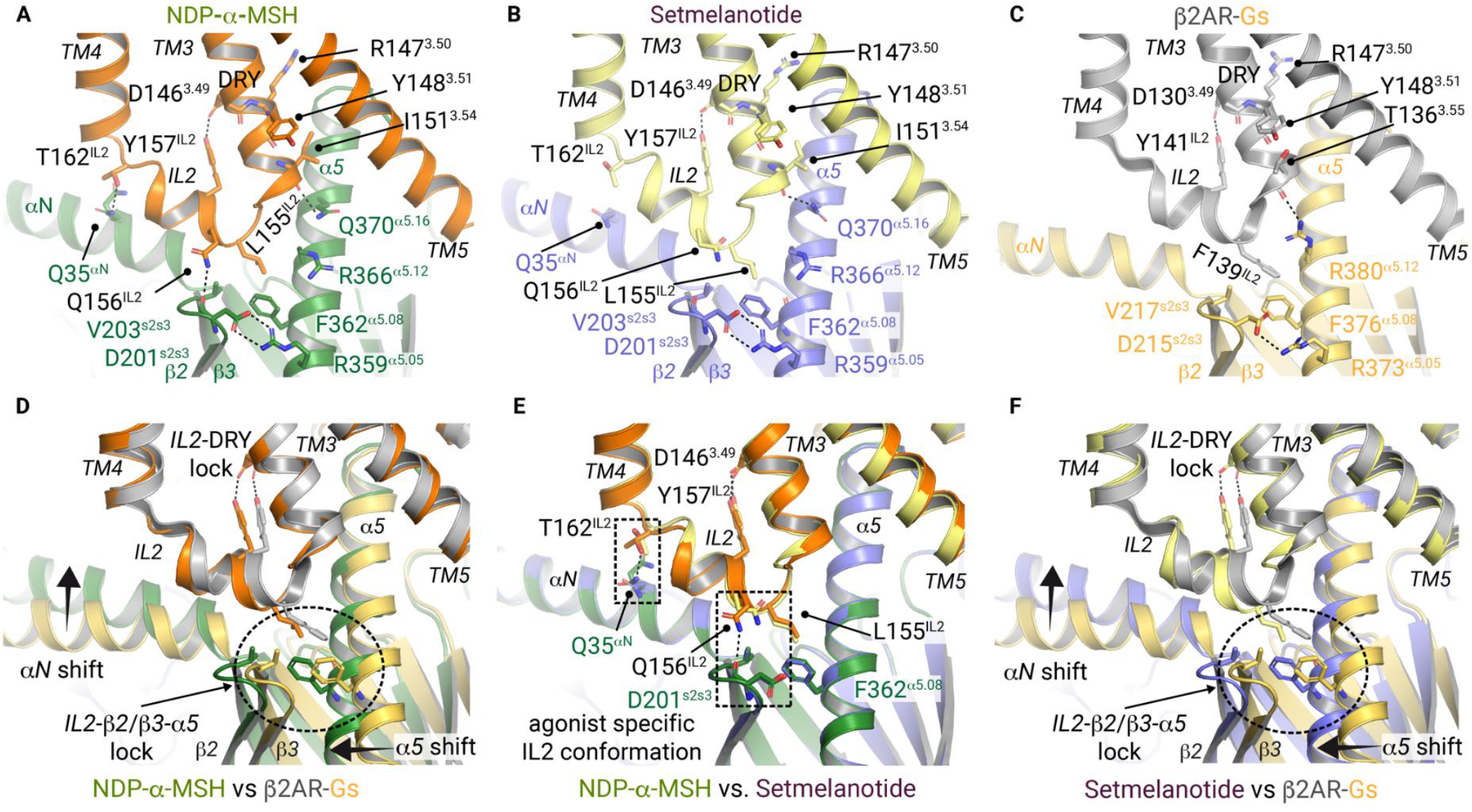
Gs-protein adjustment at IL2. **A− C** Display of the TM3− TM5 G_s_-protein binding interface of the (**A)** NDP-α-MSH, (**B)** setmelanotide bound MC4R− Gs complexes and (**C)** the β2AR**−** Gs complex (PDB ID: 3sn6). Interactions of IL2 and TM3 are displayed as black dashes. (**D− F)** The superposition of NDP-α-MSH− MC4R− Gs and setmelanotide− MC4R− Gs with β2AR− Gs complexes display the IL2-β2/β3-α5 lock of L155 ^IL2^ (MC4R) with V203 and F362 (Gαs), as well as F139 ^IL2^ (β2AR) with V217 and F376 (Gαs) adjusting the position of the α5 helix. A relative α5 shift can be noticed, that results in a slight rotation of the entire Gs-coupled to MC4R compared to β2AR, most prominently visible by an αN helix shift. **(E)** Superposition of both agonist bound MC4R− Gs structures highlight changes of the hydrogen bonds between Gαs and IL2 residues (dashed boxes).

Of note, a large number of class A GPCRs display a leucine or isoleucine at the IL2-3.58 position that often exists in combination with a tyrosine in IL2 (IL2-3.60) (Figure S20). In agreement with the essential role of IL2 and position 3.58 for Gs-coupling, previously investigated mutants L155A, R147A, T150A, and Y157A eliminate any basal signaling activity of MC4R (Yang and Tao, 2020), presumably by disturbing the interface contacts between the receptor and its effector protein.

Finally, the β2AR− Gs complex shows a tighter α5-helix engagement toward TM5 (Figures 7D− 7E), likely caused by differences in the specific IL2-3.58 to Gs-α5.08 interactions. Consequently, several further interactions in the TM3-IL2− α5-helix network differ from the β2AR− Gs complex (Figures 7A− 7C), leading to a displaced αN-helix in the active MC4R− Gs complexes (Figures 7D and 7F).

Interestingly, subtle differences at the IL2− G-protein interface can be observed between the two agonist− MC4R− Gs complex structures. First, in the NDP-α-MSH− MC4R complex, T162^IL2-3.65^ presumably forms a hydrogen bond to Q35^αN^ in the αN-helix, and Q156^IL2-3.59^ to the backbone of D201^s2s3^ in the β2− β3 turn of Gαs (Figures 7A, 7B, and 7E; Figures S22 and S23). None of these contacts are present in the setmelanotide complex, most likely due to a rotation of the side chains of T162^IL2-3.65^ and Q156^IL2-3.59^ (Figure 7B; Figures S22 and S23).

Another important difference between the two agonist− MC4R complexes is found in the interactions formed by residue H158^IL2-3.61^ in IL2. In both complexes, H158^IL2-3.61^ is part of a water-mediated interaction network to H373^α5.19^ in the Gs-α5 helix (Figures 6D− 6E). However, in the NDP-α-MSH− MC4R− Gs complex the H158^IL2-3.61^ side chain is more flexible and samples two different rotamer conformations, which is not the case in the setmelanotide− MC4R− Gs complex (Figures 6D− 6E; Figure S24). Yet, there is an additional water-mediated stabilization to A154^IL2-3.57^ in IL2 (Figure 6E). Interestingly, the H158A mutation leads to constitutive receptor activation and causes a slight increase in agonist-induced Gs-signaling (Figure 4D-4E, Tables S7 and S8 and (Yang and Tao, 2020)) in agreement with the pathogenic gain-of-function mutant H158R (Hinney et al., 2006). Compared to the antagonized MC4R structure without structurally visible interaction of H158, it is not yet obvious why mutants at H158 lead to constitutive receptor activation. However, in the context of the known constitutively activating mutants D146A, F149A, and F152A (Yang and Tao, 2020) the importance of the entire IL2− TM3 transition as a key-switch for regulation of MC4R-signaling and G-protein engagement is highlighted.

## DISCUSSION

Here we report cryo-EM structures of active state MC4R− Gs complexes bound to the FDA– approved peptide agonists NDP-α-MSH and setmelanotide (Figure 1). Both structures provide details of agonist binding and receptor activation, with further details gained when compared to the recently determined antagonized SHU9119-MC4R structure (Yu *et al*., 2020) (Figure 2 and Figure 3). Summarizing the structural findings in combination with signaling data, extracellular binding of the peptide agonists occurs in complex with the cofactor calcium and at key receptor residues in TM2, TM3 and TM6. Here we assign the agonist-dependent activation trigger at TM6 to the ligand-binding region of H264^6.54^-F261^6.51^ directly adjacent to the helix-kink at the *CWxP*^6.50^ motif, which is involved in the known “toggle-switch” activation mechanism with the resulting TM6 movement typcial for class A GPCRs (Figure 5).

Notably, extracellular interactions formed by SHU9119 are not significantly different to those formed by both agonists, but SHU9119 blocks the TM6 movement essential for MC4R signaling in the transmembrane region. SHU9119s antagonism depends on the interaction with L133^3.36^ and is therefore MC4R selective, since it acts as an agonist at MC1R (L^3.36^M) (Yang *et al*., 2002) and MC5R (L^3.36^V) with neither receptor having a leucine at the reciprocal position (Figure S12B). This conclusion is further mirror-like supported by agonistic effects of the ligand melatonan II on MC4R, which just differs from the antagonistic SHU9119 in the ligand position 1 with a smaller D-F (no contact to L133) versus the larger D-Nal in SHU9119 (Fig. 1A).

Moreover, the binding modes of agonistic ligands compared to each other and to the antagonist SHU9119 show different orientations of the Ca^2+^ binding site, which is formed by ligand and receptor residues. These differences are accompanied by local shifts in the TM3 adjustment relative to other helices (Figure 3).

Compared to the antagonized SHU9119-MC4R structure we identify several active state conformational changes that play a concerted role in signal transduction. A large movement of TM6 starting at the *CWxP*^6.50^ motif with altered side chain interactions of residues I291^7.42^ and W258^6.48^, small conformational alterations in TM7 related to local shifts at the *N(D)P*^*7*.*50*^*xxY* motif, as well as reorientations of Y^5.58^ in TM5 and the conserved *DR*^*3*.*50*^*Y* motif in TM3 are significant for the active structures. Subsequently, the conserved *DR*^*3*.*50*^*Y* motif in TM3 and the adjacent IL2 both constitute important parts of the interface to the Gs protein.

In TM3 T150^3.53^ is essential for Gs binding (Figure 4). This residue switches from being a key player in the inactive state, where it stabilizes the TM3-TM6 dual lock (Figure 6), to a binding partner of the α5 helix in Gs. This finding is remarkable considering the non-conserved nature of this class A GPCR position (Figure S12B). The Gs protein is bound further at a second essential site, the IL2. The detailed interaction pattern of IL2 to Gs is somewhat different in both agonist-MC4R structures, indicating that both ligands induce specific binding patterns to G-proteins such as Gs, but probably also to Gq (Figure 7).

A possible explanation for the different binding modes of IL2 to Gs is the specificity in agonist binding to MC4R, which depend on detailed interactions to TM3 and the transition to EL2. The setmelanotide residue R1^-3^ mediates specific contacts to TM3, which may be related to the biased signaling property of this agonist compared to MC4R agonist NDP-α-MSH without this arginine (Figure 2 and Figure 3, Table S7). In addition, comparison of previously published Gq signaling data (Clement *et al*., 2018) and the primary sequence of setmelanotide and the LY2112688 compound (Figure 1A) also suggests that the biased pharmaceutical profile of setmelanotide can be narrowed down to the direct arginine interaction to TM3 of MC4R. These differences in the ligand binding pocket suggest a first possible explanation for the enhanced Gq-signaling of setmelanotide that is crucial for its successful application in appetite regulation (Clement *et al*., 2018).

However, to fully characterize G-protein selectivity at the MC4R and to further differentiate Gq- and Gs-coupling, an agonist− MC4R− Gq complex is required. Other open questions pertaining to the unique MC4R system await structural elucidation, such as the determination of the apo-state conformation, dimeric constellations relevant to MC4R as an endocrine active GPCR (Kleinau et al., 2016) and MC4R bound with endogenous ligands such as α-, β-MSH, AgRP or the cofactor protein MRAP (Rouault et al., 2017). Altogether, the structural findings presented here will facilitate the development of new MCR subtypes and G protein-selective anti-obesity drugs.

## ACKNOWLEDGEMENT

We thank Petra Henklein for agonist peptide synthesis and Sabine Jyrch and Cigdem Cetindag for expert technical assistance. We thank Benedikt Kirmayer and Boris Schade for assistance with cryo-EM sample preparation and data acquisition. We acknowledge access to electron microscopic equipment at the core facility BioSupraMol of Freie Universität Berlin, supported through grants from the Deutsche Forschungsgemeinschaft and the state of Berlin for large equipment according to Art. 91b GG (INST 335/588-1 FUGG, INST 335/589-1 FUGG, INST 335/590-1 FUGG). We are grateful to Rhythm Pharmaceuticals, Inc. for providing several mg quantities of the setmelanotide ligand sample for first *in vitro* experiments.

This work was mainly supported by the Deutsche Forschungsgemeinschaft (DFG, German Research Foundation) through CRC 1423, project number 421152132, subproject A01 (to P.S.) and additional funding by subprojects A05 (to C.M.T.S. and P.S.), Z03 (to A.B.S. and P.S.), B02 (to P.K. and H.B.) and C01/Z04 (to P.W.H.), through DFG Grant KU 2673/6-1 (to P.K.); through CRC 1365, project number 394046635, subproject A03 (to P.S.); through Germany’s Excellence Strategies - EXC2008/1 (UniSysCat) - 390540038 (to P.S.). P.W.H. and B.K.K are supported through the Stiftung Charité and the Einstein Center Digital Future. B.K.K. is a Chan Zuckerberg Biohub investigator and an Einstein BIH visiting fellow. P.S. acknowledge the Einstein Center of Catalysis (EC2).

## AUTHOR CONTRIBUTIONS

N.A.H. established and performed the MC4R cell culture production, purification, Gs-protein production and agonist− MC4R− Gs− NB35 complexation with experimental input from D.S., M.S., B.B., A.K., M.G., D.K. and advice from D.H., B.K.K. and P.S.; N.A.H., A.K. and M.S. cloned all MC4R constructs; Antibody/Nb35 production performed from M.S., A.K. and B.B. with advise from B.K.K.; A.B.S. provided parts of the agonist peptides; J.B. and T.M. performed the initial Negative-stain screening of the sample grids, T.H. prepared cryo-EM grids, collected images, reconstructed the final map and supervised the cryo-EM procedure with input of N.A.H., M.Sch. and P.S.; T.H. and N.A.H. performed the cryo-EM data analysis; C.M.T.S and M.Sch. provided advice for cryo-EM interpretations; A.S. built and refined the cryo-EM models with contributions from N.A.H. and P.S.; N.A.H. established and performed nanoBRET assay with input of M.S. and A.K.; S.P. and H.B. performed the cAMP signaling data; S.P., N.A.H., G.K., P.K., P.S. and H.B. analyzed the signaling data; G.K. performed the sequence analyses; P.W.H. provided advice for structure interpretation; N.A.H., G.K. and P.S. analyzed the structures; N.A.H., G.K., A.S., T.H., S.P., H.B. and P.S. prepared the figures; G.K., N.A.H., and P.S. wrote the manuscript with contributions of all co-authors. P.S. initiated, funded and supervised the project. All authors edited and revised the manuscript.

## Additional Files

### Supplemental information

Methods

Figures S1 to S24 Tables S1 to S8

References supplemental information

