## Supplemental information - Methods - Supplemental Figures S1-S24 - Supplemental Tables S1-S8- for "Structures of active melanocortin-4 receptor−Gs-protein complexes with NDP-α-MSH and setmelanotide"

##### Construct preparations for cryo-electron microscopy (cryo-EM)

###### *Protein expression of human melanocortin-4 receptor (MC4R)*

Wild-type human MC4R (UniprotKB-P32245) was modified to include an N-terminal hemagglutinin signal sequence, followed by a FLAG-tag epitope (DYKDDDK). The C-terminal eGFP, followed by polyhistidine (His-) and rho-1D4 tags, is removable by HRV-3C protease cleavage (construct name: MC4R–eGFP) and was inserted into a pOET3 vector.

For the production of MC4R–eGFP, recombinant baculovirus was generated by co-transfecting *Sf*9 cells (from *Spodoptera frugiperda*) with pOET3_MC4R–eGFP and linearized BAC10:1629$_{KO}$ (Schwefel et al., 2014; Zhao et al., 2003) using Trans-IT Insect (Mirus Bio). *Sf*9 cells were cultured in SF900 II serum-free medium (Invitrogen) at 28 °C for virus generation. A 1 L preparation of *Sf*9 cells at 2 x $10^6$ cells $ml^{-1}$ were infected with 10 ml of P2 virus MC4R–eGFP virus. Cultures were grown at 27 °C, harvested by centrifugation 48 h post infection and stored at -20 °C.

###### *Protein expression and purification of $G\alpha_s\beta_1\gamma_2$ and Nb35*

Bovine $G\alpha_s$-short subunit (UniprotKB-P04896-2) in pFastbac vector and rat $G\beta_1$ (UniprotKB-P54311) and bovine $G\gamma_2$ (UniprotKB-P63212) subunits in pFastbacDual vector were previously used and described (Rasmussen et al., 2011). Heterotrimeric $G\alpha_s\beta_1\gamma_2$ protein (or named Gs) was expressed in *Trichuplusia ni* (*Tni*) insect cells, maintained in ESF 921$^{TM}$ serum free insect cell culture media (Expression Systems) at 28 °C. The virus was prepared using Bac-to-Bac$^{TM}$ baculovirus expression system (Thermo Fisher Scientific). The cells were infected with both $G\alpha_s$ and $G\beta_1\gamma_2$ virus, based on small scale titrations and harvested after 48 h post infection and stored at -20 °C. $G_s$ was purified as described previously (Rasmussen et al., 2011). The single-domain antibody Nanobody-35 (Nb35) was previously described (Rasmussen et al., 2011). Nb35 was expressed in *E. coli* strain WK6, extracted and purified by immobilized metal (Ni-NTA) affinity chromatography according to previously described methods (Rasmussen et al., 2011).

###### *Complex formation and purification*

MC4R–Gs–Nb35 complexes with both agonists NDP-α-MSH and setmelanotide (from now on abbreviated by agonist) complexes were formed in *Sf9* membranes. *Sf9* cell pellets containing MC4R-eGFP were resuspended in 20 mM HEPES pH 7.5, 50 mM NaCl, 2 mM $MgCl_2$, 1 mM $CaCl_2$, 25 µM tris (2-carboxyethyl) phosphine (TCEP), 25 U/ml apyrase (New England Biolabs) 2.5 mg/ml leupeptin (Enzo Life Sciences, Inc.), 0.16 mg/ml benzamidine (Sigma-Aldrich) and 1 µM of the respective agonist (in-house peptide synthesis). Gs was pre-incubated with Nb35 was added and incubated overnight at 4 °C. The membrane sample containing agonist-MC4R–Gs–Nb35 complex was collected by centrifugation at 46.000xg and carefully resuspended in 20 mM HEPES pH 7.5, 150 mM NaCl, 2 mM $MgCl_2$, 1 mM $CaCl_2$, 25 µM TCEP, 2.5 mg/ml leupeptin, 0.16 mg/ml bezamidine, 1 µM agonist and 1% n-dodecyl β-D-maltoside (DDM), 0.1 % cholesteryl hemisuccinate (CHS) (Anatrace, Inc.). After 2 h the solubilized protein was separated from insoluble remains by centrifugation at 46.000 xg. The supernatant was diluted twofold with 20 mM

HEPES pH 7.5, 150 mM NaCl, 2 mM $MgCl_2$, 5 mM $CaCl_2$, 25 µM TCEP, 2.5 mg/ml leupeptin, 0.16 mg/ml benzamidine and 1 µM agonist. ANTI-FLAG M1 resin (Sigma-Aldrich) was added and incubated for 2 h at 4 °C rotating. M1 resin was collected by centrifugation (500 x g, 5 min) and loaded into a wide-glass column and washed for 5 column volumes with wash buffer (20 mM HEPES pH 7.5, 150 mM NaCl, 1mM $CaCl_2$, 25 µM TCEP, 1 µM agonist) with 0.1 % DDM and 0.01 % CHS. Followed by a incubation in wash buffer with 0.8 % lauryl maltose neopentyl glycol (LMNG), 0.08 % CHS (Anatrace, Inc.) and 0.02 % DDM. Subsequently, LMNG/CHS concentration was lowered in a stepwise manner to 0.01% LMNG, 0.001 % CHS over a period of 1 h. Elution of the complex was initiated by addition of 20 mM HEPES pH 7.5, 150 mM NaCl, 25 µM TCEP, 1 µM agonist, 0.01 % LMNG, 0.001 % CHS, 5 mM EDTA and 0.2 mM DYKDDDDK peptide (GenScript Biotech). C-terminal eGFP was removed by the addition of HRV-3C protease protease (in-house purified), incubated at 4 °C overnight. After concentration the agonist–MC4R–Gs–Nb35complex was loaded onto a Superdex 200 Increase 5/150 GL (Sigma-Aldrich). Receptor containing fractions were concentrated to 5 mg/ml and directly vitrified. 1 L of MC4R–eGFP expressing cells yielded 0.25 mg complex.

#### Cryo-electron microscopy

##### *Cryo-EM sample preparation and image acquisition*

Vitrification of NDP-α-MSH and setmelanotide–MC4R–Gs–Nb35 complex was conducted immediately after sample preparation at a concentration of 1.2 mg/ml and 5 mg/ml, respectively. 3.8 µl of the sample was applied to glow-discharged holey gold grids (UltrAuFoil R1.2/1.3 300 mesh, Quantifoil Micro Tools GmbH), blotted for 4 s and plunge-frozen in liquid ethane using an FEI Vitrobot Mark IV (Thermo Fisher Scientific) set to 10 °C and 100% humidity.

Images were acquired using a FEI Titan Krios G3i microscope (Thermo Fisher Scientific) operated at 300 kV equipped with a FEI Falcon 3EC detector (Thermo Fisher Scientific) running in counting mode at a nominal magnification of 96,000x giving a calibrated pixel size of 0.832 Å/px. Movies were recorded for 40.78 s accumulating a total electron dose of 40 $e^-/Å^2$ fractionated into 33 frames. EPU 2.8 was utilized for automated data acquisition with AFIS enabled using a nominal defocus between -0.8 and -2 µm.

A total of 5618 micrographs were collected for NDP-α-MSH–MC4R–Gs–Nb35 and 7583 micrographs for setmelanotide–MC4R–Gs–Nb35. These were used for further image processing. Further details are given in the Table 1.

##### *Cryo-EM image processing*

The entire data analysis was conducted within the cryoSPARC v2.15 framework (Figures S3-S6). Image analysis of the NDP-α-MSH–MC4R–Gs–Nb35 dataset (Figures S3 and S4) started with movie alignment and dose-weighting using “Patch motion correction” followed by “Patch CTF estimation”. Initial particle picking was done with “Blob picker“ using a particle diameter of 180 Å. Particle images were extracted with a box size of 280 px, Fourier-cropped to 70 px (3.328 Å/px). After reference-free 2D classification, selected class averages were used for template-based particle picking with a 170 Å particle mask. A total of 2,746,119 particle images were subjected to two cycles of 2D classification to clean the dataset. Ab initio reconstruction of particle images belonging to shiny classes was applied to generate a reference model for 3D classification after which 260,451 particle images were selected for further processing. Homogeneous refinement after re-extraction of the particles, Fourier-cropped to 140 px (1.664 Å/px) generated a 3D reconstruction of 3.45 Å global resolution. Another round of heterogeneous refinement was applied to finally select 221,682 particle images for unbinned extraction (280 px, 0.832 Å/px). Iterations of homogeneous refinement and Global CTF refinement were applied to correct for higher order aberrations yielding a final reconstruction of 2.86 Å resolution after non-uniform (NU) refinement (Punjani et al., 2020). Using NU-refinement, masking of the all-helical domain was not necessary to yield a high-resolution map (Figure 1; Figure S3 and S4). Processing of the setmelanotide–MC4R–Gs–Nb35 data (Suppl. Fig S5-S6) was done as described for the NDP-α-MSH–MC4R–Gs–Nb35 dataset using the previously generated templates for picking of 4,330,500 particle images. After a single round of 2D classification, 4,267,612 particle images were subjected to two iterative rounds of 3D classification with the NDP-α-MSH-MC4R-$G_s$αβγ-Nb35 reconstruction as reference filtered to 30 Å. Micrographs with local motions above 10 px or estimated resolutions worse than 4 Å were discarded, leaving a total of 797,185 particle images for another round of 3D classification. Homogeneous refinement of 431,973 particle images after re-extraction with a box size of 280 px (0.832 Å/px) yielded a resolution of 2.82 Å that could be improved to 2.77 Å by CTF refinement. After a final 3D classification 370,621 particles were selected for NU refinement resulting in a 2.58 Å reconstruction. (Figure 1, Figure S6)).

***Model building and refinement***

The models of the NDP-α-MSH-MC4R–Gs–Nb35 complex as well as setmelanotide–MC4R–Gs–Nb35 complex were derived from the inactive MC4R structure (PDB ID: 6w25 (Yu et al., 2020)) together with the $G_s$αβγ–Nb35 complex of the β2-adrenergic receptor–Gs complex (PDB ID: 3sn6 (Rasmussen et al., 2011)) as initial models. Both MC4R complexes were built and adjusted manually using the program COOT (Emsley et al., 2010). Model building for the MC4R ligands NDP-α-MSH and setmelanotide was started *de novo* using COOT (Emsley et al., 2010). Local-refined as well as overall cryo-EM maps were used to add water molecules. After every round of manual refinement and for the final round Real-space refinement (Afonine et al., 2018) was performed with the program PHENIX (Adams et al., 2010) using geometric restraints, a global minimalization protocol and B-factor refinement. Both models were additionally refined with isotropic B-factors in reciprocal space using REFMAC5 (Vagin et al., 2004) of the CCP4 (Collaborative Computational Project, number 4) software suite (Winn et al., 2011). The refinement was carried out in the resolution range of 233-2.88Å and 233-2.6Å for the NDP-α-MSH–MC4R and setmelanotide–MC4R complexes, respectively (Suppl. Tab.1).

The final model of the NDP-α-MSH–MC4R complex includes the following amino acids (based on the final overall cryo-EM map); MC4R: 40-108; 118-230, 239-316; NDP-α-MSH: 1-13; $G_s$α-protein: 13-47; 194-236; 249-280; 293-306; 322-380; $G\beta_1$: 3-340; $G\gamma_2$: 9-63; Nb35: 1-128.

The final model of the setmelanotide–MC4R complex includes the following amino acids (based on the final overall cryo-EM map); MC4R: 40-107; 117-230, 240-316; setmelanotide: 1-8; $G_s\alpha$-protein: 14-47; 193-236; 248-280; 293-310; 318-351; 355-380; $G\beta_1$: 4-340; $G\gamma_2$: 9-63; NB35: 1-128. Structure validation was performed with the programs PHENIX (Adams et al., 2010), MolProbity (Chen et al., 2010), SFCHECK (Vaguine et al., 1999) and OneDep of the Protein Data Bank (Winn et al., 2011). Potential hydrogen bonds and van der Waals contacts were analysed using the programs HBPLUS (McDonald and Thornton, 1994) and LIGPLOT 1.45+ (Laskowski and Swindells, 2011). All structure superpositions of backbone α-carbon traces were performed using the CCP4 program LSQKAB (Collaborative Computational Project, 1994). All molecular graphics representations in this work were created using the PyMol Molecular Graphics System Version 1.3 (Schrödinger, LLC, New York, NY) and UCSF Chimera (Pettersen et al., 2004).

Coordinates and structure factors have been deposited in the Protein Data Bank (PDB, (Berman et al., 2000)) with identification codes (XXX and XXX).

#### MC4R ligand binding assays

##### *Saturation and competition binding assay using NanoLuc™ Luciferase assay (nanoBRET)*

Wild-type MC4R was modified to include an N-terminal hemagglutinin signal sequence, followed by Luciferase (NanoLuc™ Luciferase; Promega) (Stoddart et al., 2015) and cloned into pMT4 vector. Human embryonic kidney 293 (HEK293T) cells grown in DMEM/F-12 (Thermo Fisher Scientific) medium (supplemented with L-Glutamin, HEPES, phenol red, sodium pyruvate pH 6.9-7.3) (Thermo Fisher Scientific, Gibco) were transiently transfected using FuGENE® HD transfection reagent (Promega). 20.000 cells per well were seeded in white corning assay 96 well plates. After 24 h medium was removed and replaced with 75 µl ligand serial dilutions in Opti-MEM reduced serum media (Thermo Fisher Scientific) without phenol red and incubated for 2 h. For saturation experiments, TAMRA-NDP-α-MSH (TAMRA-NDP) labeled with the fluorophore 5-carboxytetramethylrhodamine (TAMRA) was titrated from 1 µM to 1 pM. For competition experiments each well contains 10 nM TAMRA-NDP and the competing ligand was titrated from 1µM to 10 pM for setmelanotide. Non-specific binding was measured by the addition of 20 µM NDP-α-MSH, in order to saturate the ligand binding pocket with non-fluorescent ligand. After 2h, 25 µl Furimazine (Promega) was added and incubated for 15 min. Luminescence and resulting BRET (bioluminescence resonance energy transfer) was measured in Spectramax ID3 (Molecular Devices) and CLARIOstar Plus (BMG LABTECH) plate reader with 460 nm (short-pass filter) and 610 nm (long-pass filter). The BRET ratio was calculated by the quotient of long-pass divided by short-pass (Stoddart et al., 2015). GraphPad PRISM 8 (GraphPad Software Inc.) was used for analysis by sigmoidal dose-response (variable slope) for dose-response measurements and one site - Fit $K_i$ for competition experiments.

#### Functional characterization by NanoGlo®HiBiT and AlphaScreen™ assays

##### *Cell lines, cloning and reagents*

HEK293 cell line was purchased from ATCC. Cells were authenticated by single nucleotide

polymorphism (SNP) analysis and regularly tested for mycoplasma contamination. Cultivation took place in L-glutamine containing minimal essential medium MEM (Merck Biochrom) supplemented with 5 % fetal bovine serum FBS (Thermo Fisher Scientific, Gibco) and 1% non-essential amino acids NEA (Merck Biochrom) at 37 °C and humidified air containing 5 % $CO_2$. For cAMP accumulation assays, 1.5 x $10^4$ cells per well were seeded in poly-L-lysine coated (Merck Biochrom) translucent 96 well plates (Falcon) and incubated for 24 h. In an identical fashion, determination of total and cell surface expression (NanoGlo®HiBiT assay, Promega) were performed in white opaque, poly-L-lysine coated 96 well plates (Corning #3917).
MC4R cDNA was amplified from genomic DNA and cloned into eukaryotic expression vector pcDps. The receptor was N-terminally tagged with the hemagglutinin (5'-YPYDVPDYA-3') epitope (HA) for cAMP measurements and luciferase-based assays. For NanoGlo®HiBiT assays, MC4R was cloned into pBiT3.1-N (Promega) using EcoRI/BamHI restriction sites, resulting in HiBiT protein tag N-terminally spaced by eleven amino acids. All single point mutations were incorporated into the expression vectors using site-directed mutagenesis. Cloned constructs were sequenced and verified with BigDye-terminator sequencing (PerkinElmer Inc.) using an automatic sequencer (ABI 3710 XL; Applied Biosystems). α-MSH, NDP-α-MSH and 3-Isobutyl-1-methylxanthine (IBMX) were purchased from Sigma-Aldrich.

##### *Transfection*

For determination of cAMP accumulation HEK293 cells were transfected 24 h after seeding. Cells were transfected with 45 ng plasmid DNA and 0.45 µl Metafectene (Biontex) per well in MEM without supplements. For the NanoGlo®HiBiT assay, transfection was performed as described previously elsewhere (Paisdzior et al., 2020). In short, low amounts of HiBiT-tagged receptor mutants were transfected (0.45 ng/well) and carrier DNA (pGEM-3Zf(+), Promega) was added to 45 ng DNA/well in total in advanced MEM (Thermo Fisher Scientific, Gibco) to ensure comparable transfection conditions to the other performed assays. Transfection for both HiBiT-assay (cell surface and total expression) was carried out simultaneously to ensure comparability.

##### *Determination of total and cell surface expression using NanoGlo®HiBiT assay*

The amount of receptors expressed on the cell membrane as well as total cell expression as determined using the NanoGlo®HiBiT detection system (Promega). Assay was performed according to manufacturer's protocol (rapid measurements) and has been described elsewhere (Paisdzior et al., 2020). In short, 48 h after transfection, media was changed in Opti-MEM reduced serum media (Thermo Fisher Scientific, Gibco) without phenol red (50 µl/well) to remove background noise. Determination of cell surface expression was performed by injection of 50 µl of HiBiT extracellular substrate in the appropriate buffer supplemented with LgBiT. Total expression as carried out similarly, with 50 µl/well of HiBiT Lytic substrate combined with LgBiT in the appropriate buffer containing detergents to lyse the cells (Promega). After orbital shaking for 3 min at 300 cycles per minute, plates were incubated for 10 min at room temperature. Luminescence was measured using a plate reader (Mithras LB 940, Berthold Technologies). As background control, cells transfected with empty vector pcDNA3 were used and values were subtracted from the sample emissions.

***Determination of Gs activation by measurement of cAMP accumulation using AlphaScreen™ assay***

Ligand-induced activation of MC4R was determined by using the AlphaScreen™ assay (Perkin Elmer Life Science) according to manufacturer's protocol and described elsewhere (Biebermann et al., 2006). In brief, 48h post transfection, cells were challenged with either α-MSH or NDP-α-MSH (1 µM to 0.1 nM) in stimulation buffer (138 nM NaCl, 6 mM KCl, 1 mM $MgCl_2*6H_2O$, 5.5 mM glucose, 20 mM HEPES, 1 mM $CaCl_2 * 2H_2O$, 0.1 % BSA, pH 7.4) containing 1 mM IBMX for 40 min for at 37 °C and 5 % $CO_2$. Incubation was stopped by freezing cells at -80 °C for 10 min prior to cAMP measurements. The determination of cAMP accumulation was performed following to the manufactures' instructions (Perkin Elmer Life Science) and measured with a plate reader (Mithras LB 940, Berthold Technologies).

***Statistical analysis***

Statistical analysis was performed using GraphPad Prism 6. Appropriate tests were carried out and indicated for each individual data set. Statistical significance was set at $*p \leq 0.05$, $**p \leq 0.01$, $***p \leq 0.001$ and $****p \leq 0.0001$.Concentration-response curves of each experiment were analyzed by fitting a non-linear regression model for sigmoidal response in GraphPad PRISM 6 (GraphPad Software Inc.) to determine $EC_{50}$ values. Statistics for all functional data are given in Tables 7 and 8.

#### SUPPLEMENTAL FIGURES

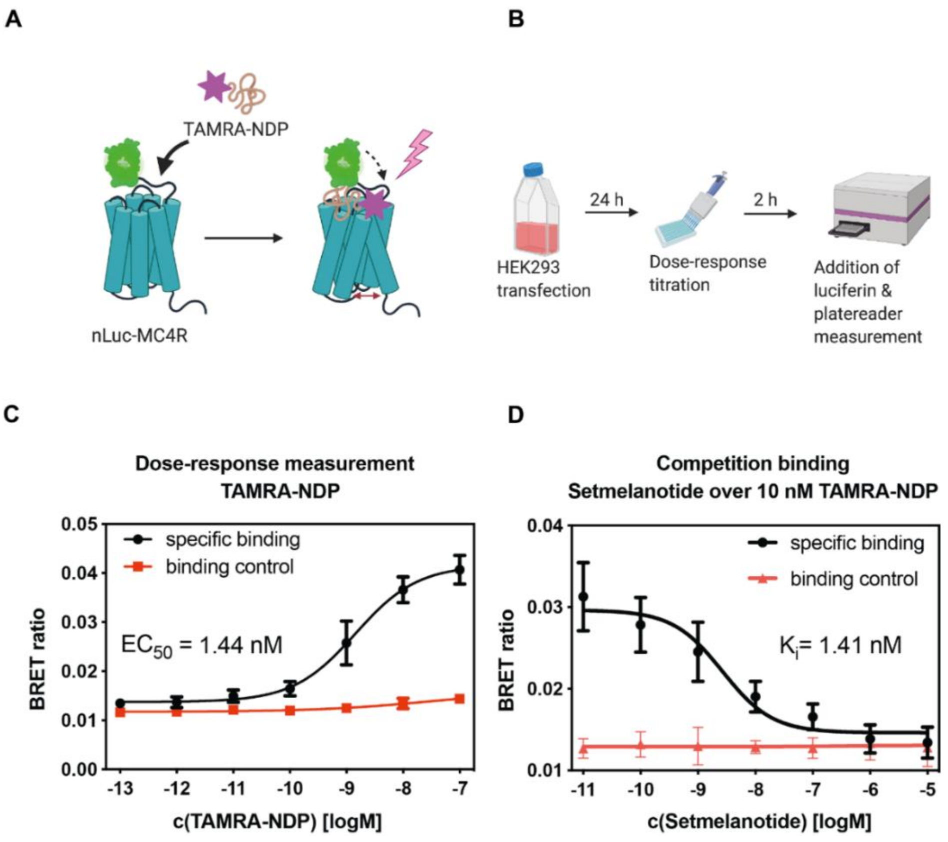


**Figure S1:** ***Experimental set-up of NanoLuc™ Luciferase assay and ligand binding data.***
(**A**) Schematic representation of NanoLuc™ Luciferase (nLuc) based MC4R ligand binding assay. nLuc protein was fused at the receptor N-terminus. Bioluminescence resonance energy transfer (BRET) is observed in dependence of the relative distance of the fluorescent (with the fluorophore 5-carboxytetramethylrhodamine (TAMRA)) labelled NDP-α-MSH (TAMRA-NDP) and nLuc-MC4R.
(**B**) Ligand binding assay workflow, HEK293T cells were infected 24 h prior, followed by media exchange against ligand titrations. Ligand binding equilibration is ensured by 2 h incubation time with subsequent addition of the nLuc substrate luciferin and measurement of the short-pass filter (460 nm) and long-pass filter (610 nm) using a fluorescent plate reader. The BRET ratio is the quotient of long-pass by short-pass.
(**C**) Titration of TAMRA-NDP from 10 µM to 0.1 pM is plotted as dose-response measurement with an $EC_{50}$ of 1.44 nM. (**D**) Setmelanotide binding was determined by competing the agonist setmelanotide against 10 nM TAMRA-NDP with a resulting $K_i$ of 1.41 nM. The addition of 20 µM non-fluorescent labeled NDP-α-MSH enhances the BRET effect induced by non-specific binding (binding control). (**A**, **B**) was created by *Biorender*.

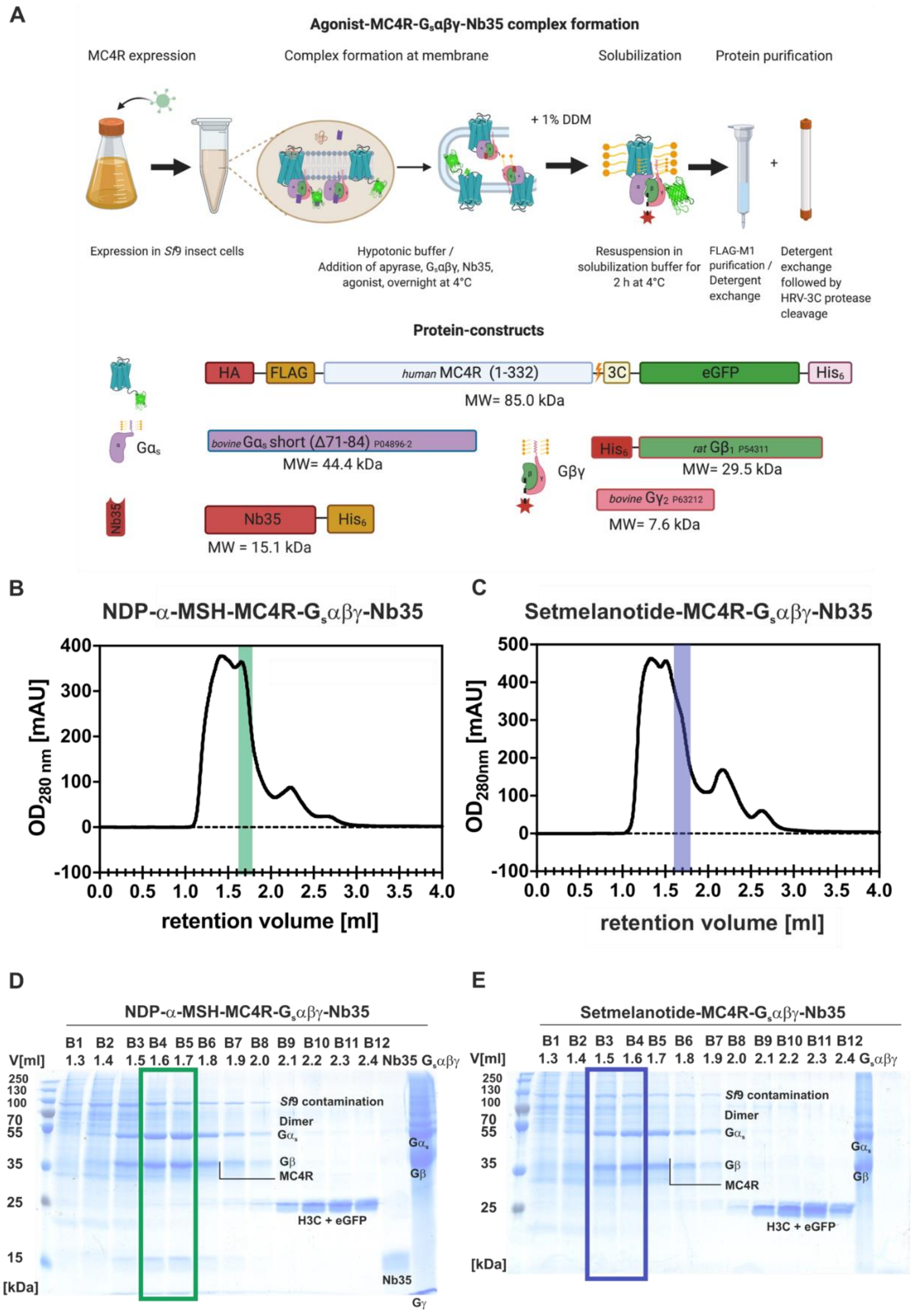


**Figure S2:** ***Workflow and biochemistry data*.**

(**A**) Workflow for the assembly of the MC4R-$G_s\alpha\beta\gamma$ complexes stabilized by NDP-α-MSH or setmelanotide. MC4R−eGFP and $G_s\alpha\beta\gamma$ (Gs) were expressed in *Sf*9 and *Tn*i cells, respectively. Gs was purified and added in excess to MC4R−eGFP expressing *Sf*9 cells resuspended in a hypotonic buffer, containing apyrase, Nanobody 35 (Nb35) and 1µM of agonists (NDP-α-MSH or setmelanotide). After overnight incubation membranes were resuspended in a buffer containing 1 % DDM and 0.1 % CHS. Followed by a FLAG M1 antibody purification. During the washing steps, DDM was exchanged against 0.01 % LMNG and 0.001 % CHS. After elution HRV-3C protease was added and the $His_6$ tag at Gβ and eGFP at MC4R was cleaved overnight. The agonist−MC4R−Gs−Nb35 complexes were separated from HRV-3C protease and remaining eGFP by size-exclusion chromatography using a Superdex 200 Increase 5/150 GL column for the complexes with

(**B**) NDP-α-MSH and

(**C**) setmelanotide. Subsequent SDS gel chromatography of

(**D**) NDP-α-MSH−MC4R−Gs−Nb35 and

(**E**) setmelanotide−MC4R−Gs−Nb35 were applied to confirm the stoichiometric ratio of MC4R and Gs. Gs-protein and Nb35 were used as controls in the last two lanes. For both complexes fractions B4 and B5 were concentrated to 5 mg/ml and directly vitrified. Figure (**A**) was created by *Biorender*.

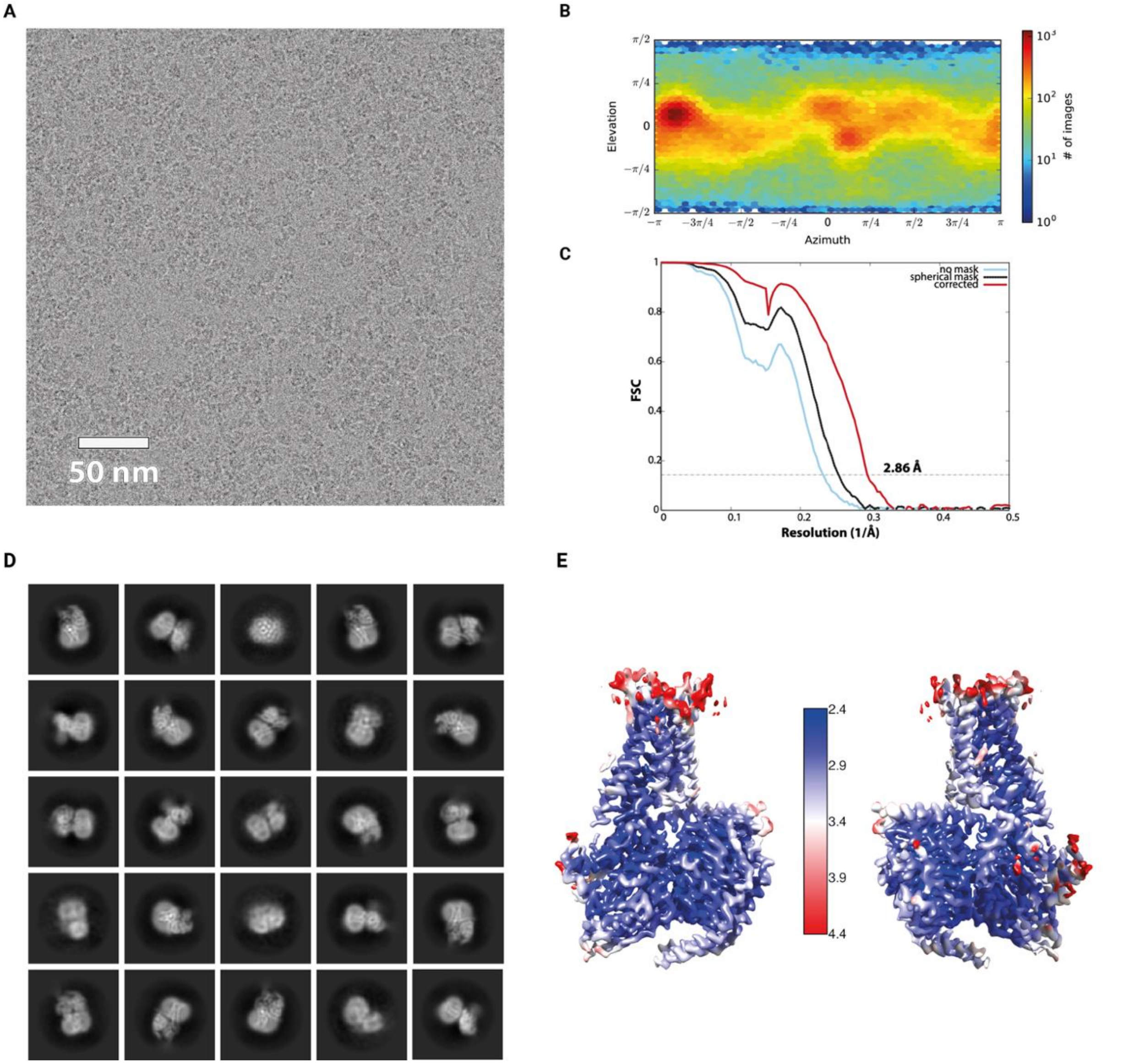


**Figure S3:** ***Cryo-EM data analysis of the active NDP-α-MSH−MC4R−Gs−Nb35 complex dataset***.

(**A**) Representative cryo-electron micrograph. The scale bar corresponds to 50 nm in the image.

(**B**) Distribution of projection directions as estimated during homogeneous refinement with program cryoSPARC.

(**C**) Global resolution estimation by Fourier shell correlation calculations (FSC = 0.143 cutoff) after "gold standard" refinement. The light blue curve was calculated without masking, the black curve by applying a spherical mask and the red one after phase randomization using a soft-mask.

(**D**) Representative 2D class averages confirm random distribution of projection directions.

(**E**) Representation of local resolution estimation determined with cryoSPARC. The final cryo-EM density is colored according to the local resolution ranging from dark blue (2.4 Å) to red (4.4 Å).

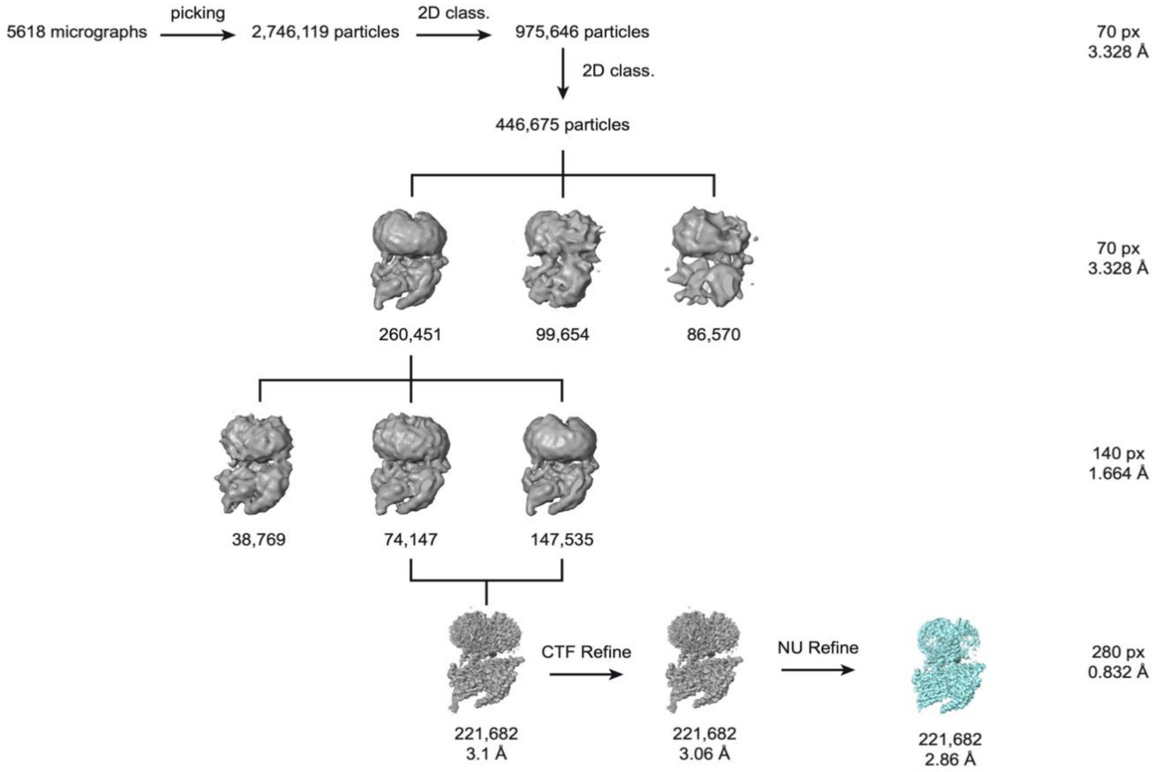


**Figure S4:** ***Cryo-EM refinement sorting scheme of the active NDP-α-MSH−MC4R−Gs−Nb35 complex dataset.*** After template-based particle picking, 2,746,119 particle images were extracted and Fourier-cropped to a box size of 70 px (pixel size 3.328 Å). After two subsequent runs of reference-free 2D classifications, 446,675 particles were selected and subjected to heterogeneous 3D refinement yielding 260,451 particle images. Re-extraction of particle images Fourier-cropped to 140 px box size (1.664 Å pixel size) was followed by another round of heterogeneous refinement. The reconstruction of two classes was virtually identical by visual inspection, therefore particle images were combined and re-extracted unbinned with a box size of 280 px (0.832 Å/px). Homogeneous refinement generated a reconstruction with a resolution of 3.1 Å, which was improved to 2.86 Å by CTF refinement followed by non-uniform (NU) refinement.

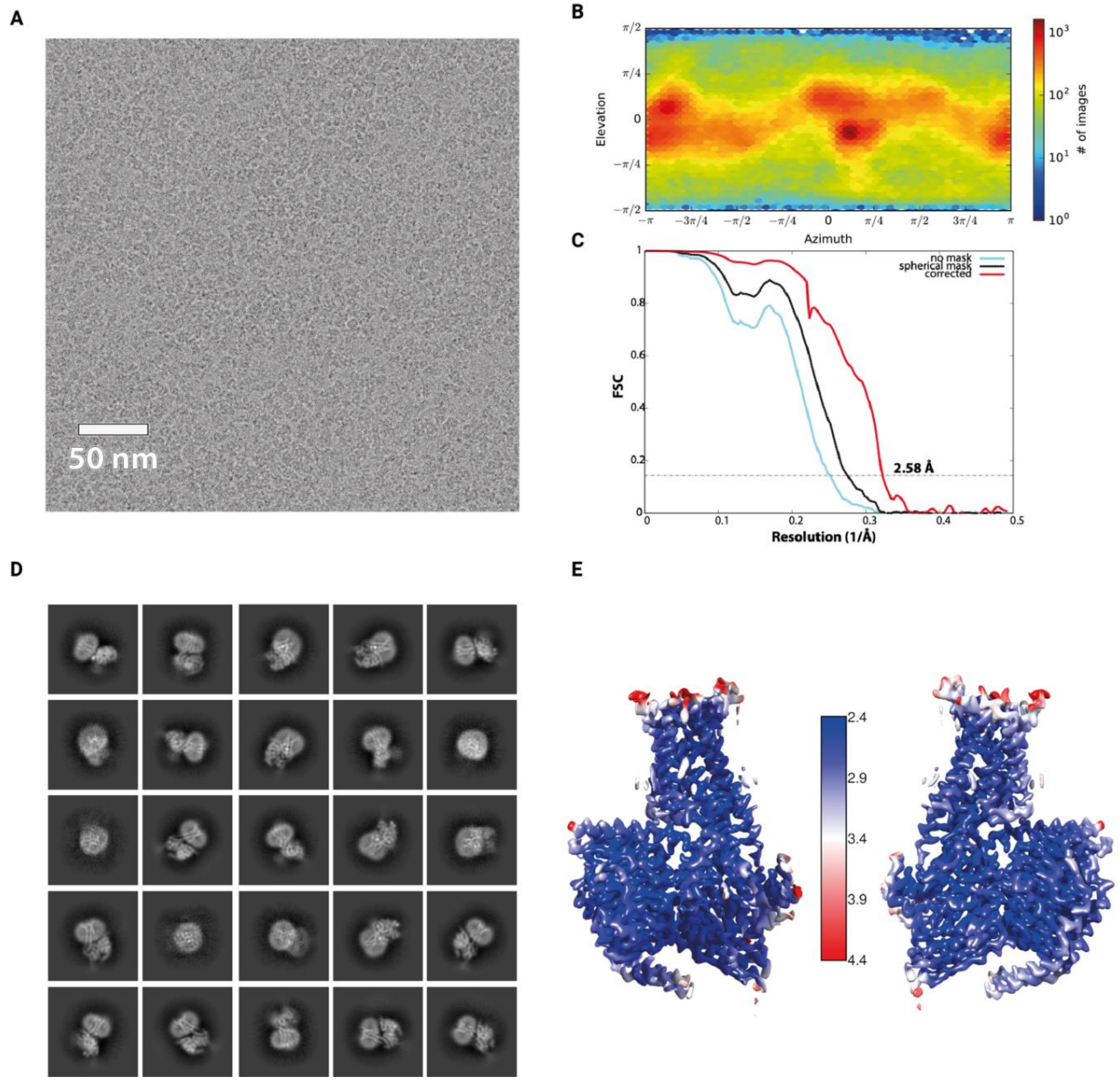


**Figure S5*****: Cryo-EM data analysis of the active setmelanotide−MC4R−Gs−Nb35 complex dataset.***

(**A**) Representative cryo-electron micrograph. The scale bar represents 50 nm in the image.

(**B**) Distribution of projection directions as estimated during homogeneous refinement with cryoSPARC.

(**C**) Global resolution estimation by Fourier shell correlation calculations (FSC = 0.143 cutoff) after "gold standard" refinement. The light blue curve was calculated without masking, the black curve by applying a spherical mask and the red one after phase randomization using a soft-mask.

(**D**) Representative 2D class averages confirm random distribution of projection directions.

(**E**) Representation of local resolution estimation determined with cryoSPARC. The final cryo-EM density is colored according to the local resolution ranging from dark blue (2.4 Å) to red (4.4 Å).

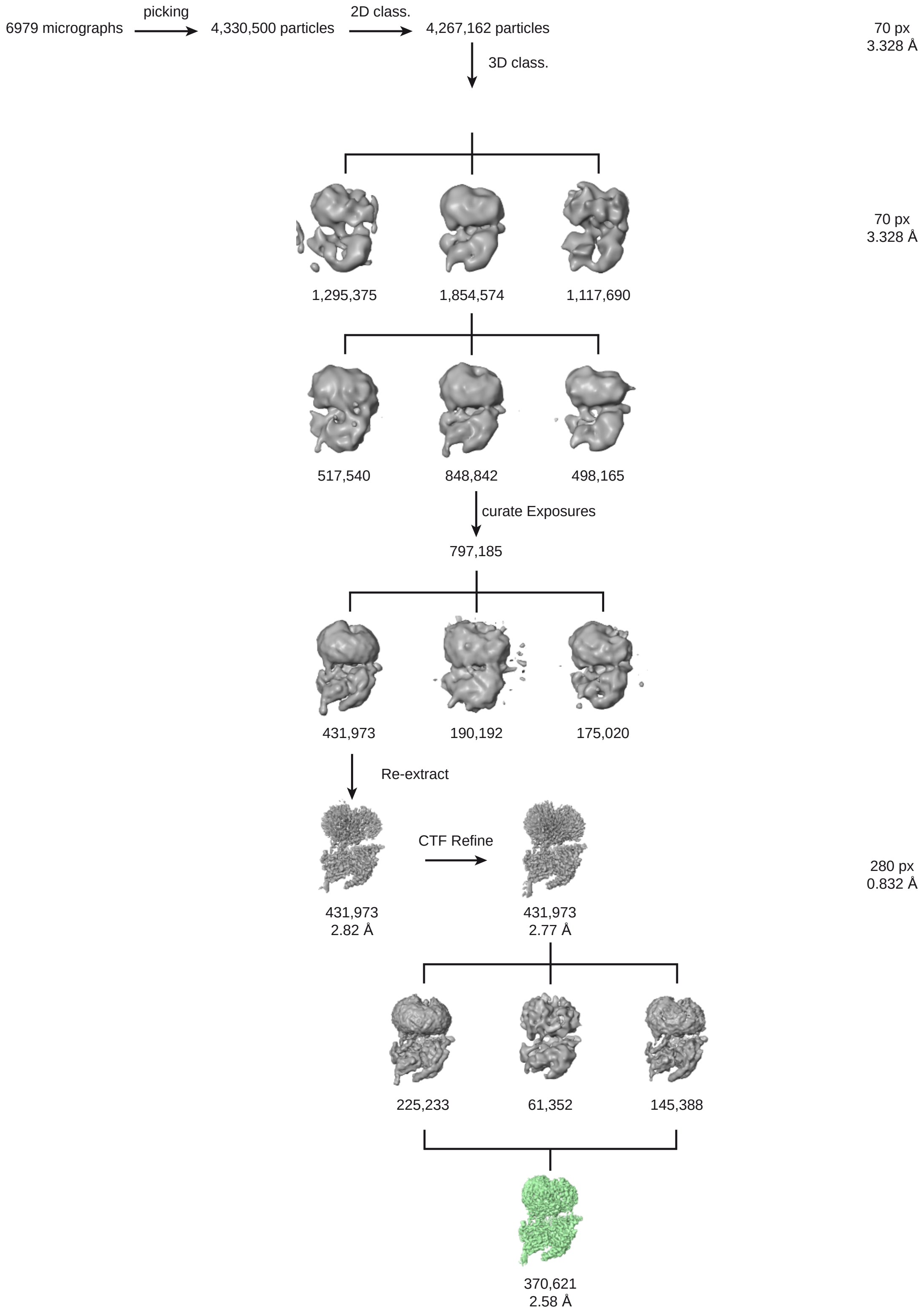


**Figure S6:** ***Cryo-EM refinement sorting scheme of the active setmelanotide−MC4R−Gs−Nb35 complex dataset.*** Initially, 4,330,500 particles were picked and extracted by Fourier-cropping with a box size of 70 px (3.328 Å/px). 2D classification was followed by two iterative 3D classifications using the NDP-α-MSH−MC4R−Gs−Nb35 filtered to 30 Å as template yielding 848,842 particle images. Micrographs were curated omitting resolutions above 4 Å and high local motion, leaving 797,185 particles for another iteration of heterogeneous refinement. 431,973 particles were re-extracted at full resolution (0.832 Å/px) and subjected to homogeneous refinement. The resulting 2.82 Å reconstruction was improved by CTF refinement to 2.77 Å. Another heterogeneous refinement was conducted to select 370,621 particle images for a final NU refinement, yielding a 2.58 Å reconstruction of the setmelanotide−MC4R−Gs−Nb35 complex.

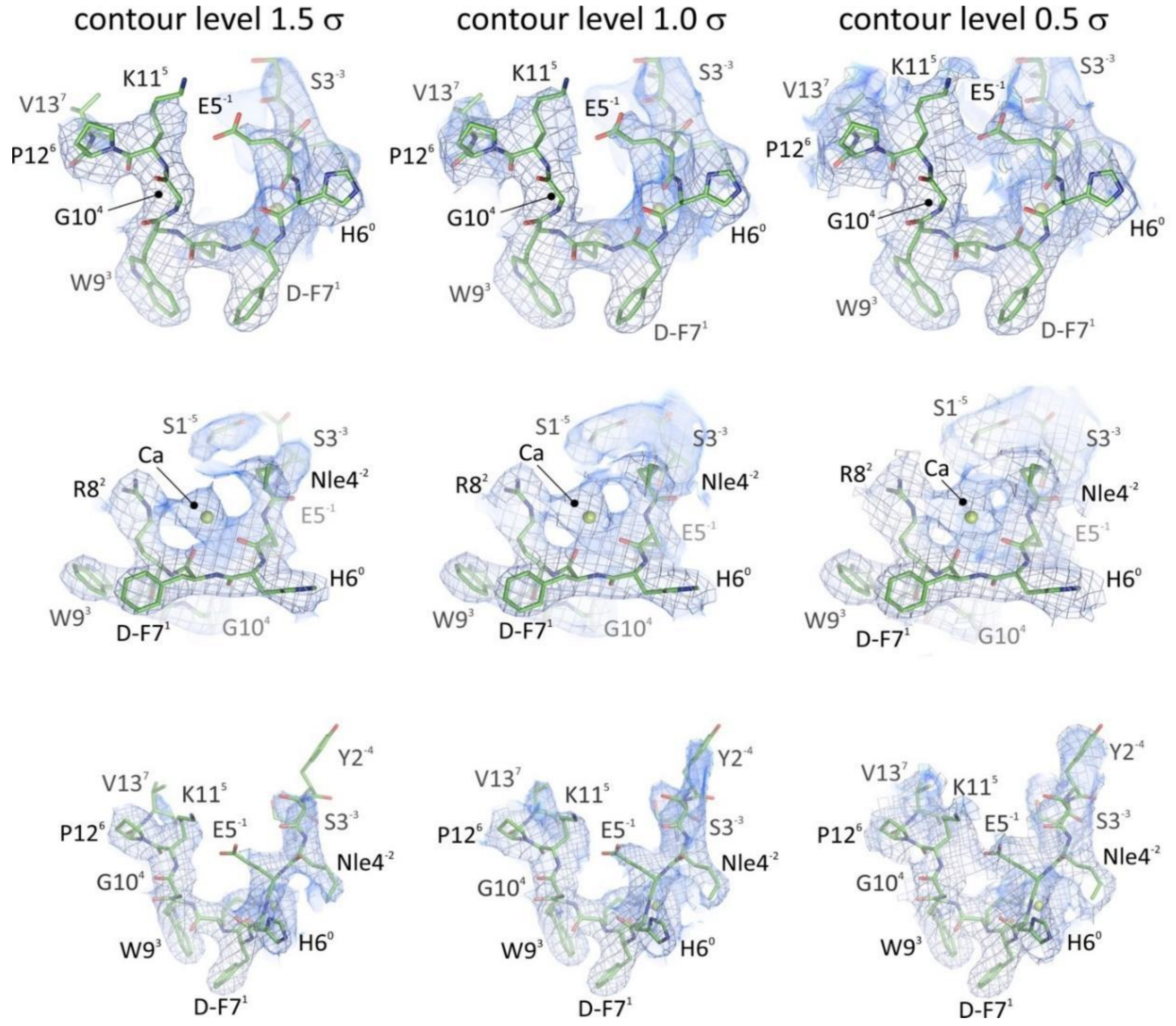


**Figure S7:** ***Quality of the cryo-EM density map of the NDP-α-MSH ligand of NDP-α-MSH−MC4R−Gs−Nb35 complex.*** Three different views (top to bottom) on the NDP-α-MSH ligand (green color) together with the coordinating calcium ion (lemon green color). All figures show cryo-EM densities of the ligand (light blue colored mesh/volume) contoured at three different contour levels (1.5 (left), 1.0 (middle) and 0.5 σ (right)). NDP-α-MSH is depicted as sticks and the calcium ion as sphere representation.

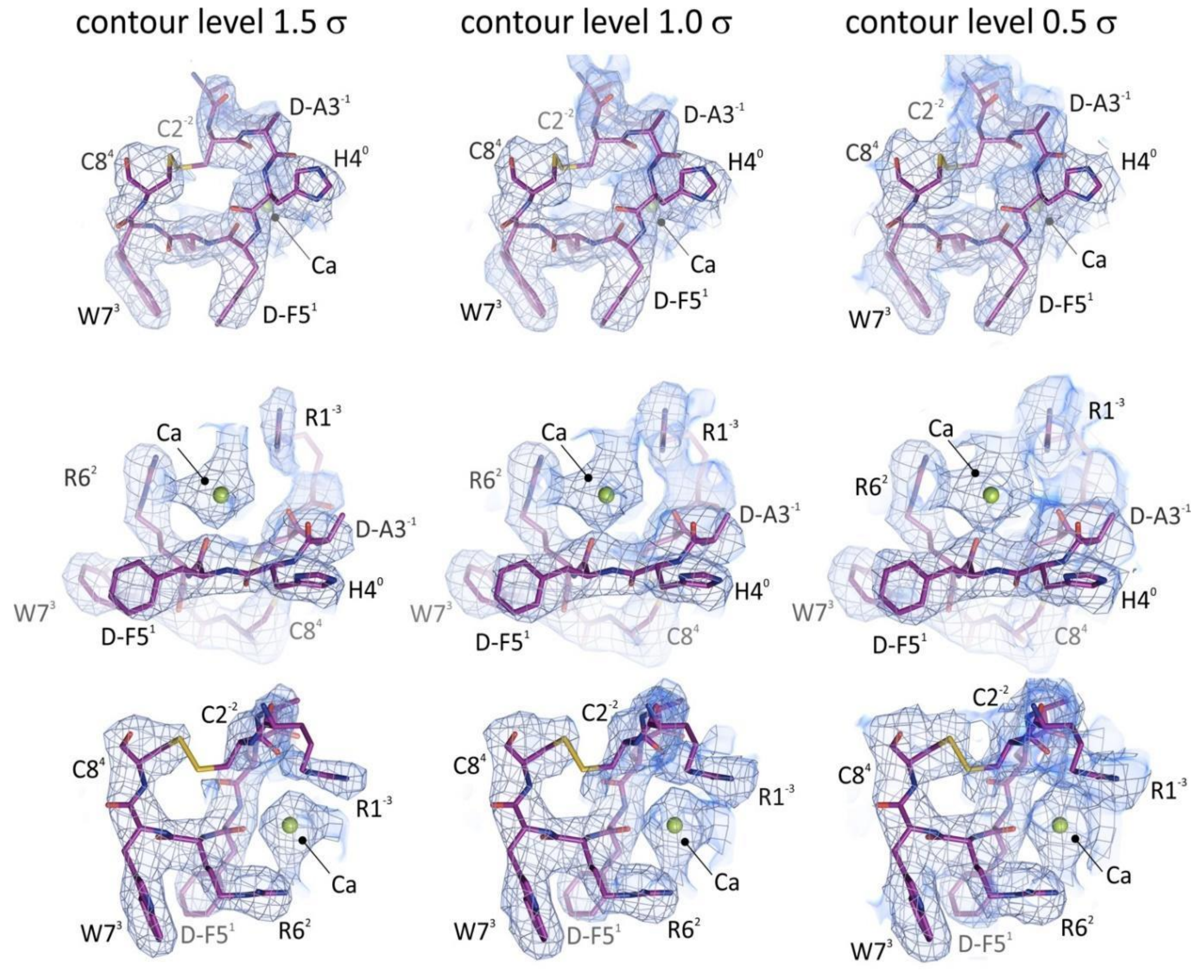


**Figure S8:** ***Quality of the cryo-EM density map of setmelanotide of the setmelanotide−MC4R−Gs−Nb35 complex.*** Three different views (top to bottom) on the setmelanotide ligand (purple color) and the binding calcium ion (lemon green color). All figures show the cryo-EM densities of the ligand (light blue colored mesh/volume) contoured at three different contour levels (1.5 (left), 1.0 (middle) and 0.5 σ (right)). Setmelanotide is depicted as sticks and the calcium ion as sphere representation.

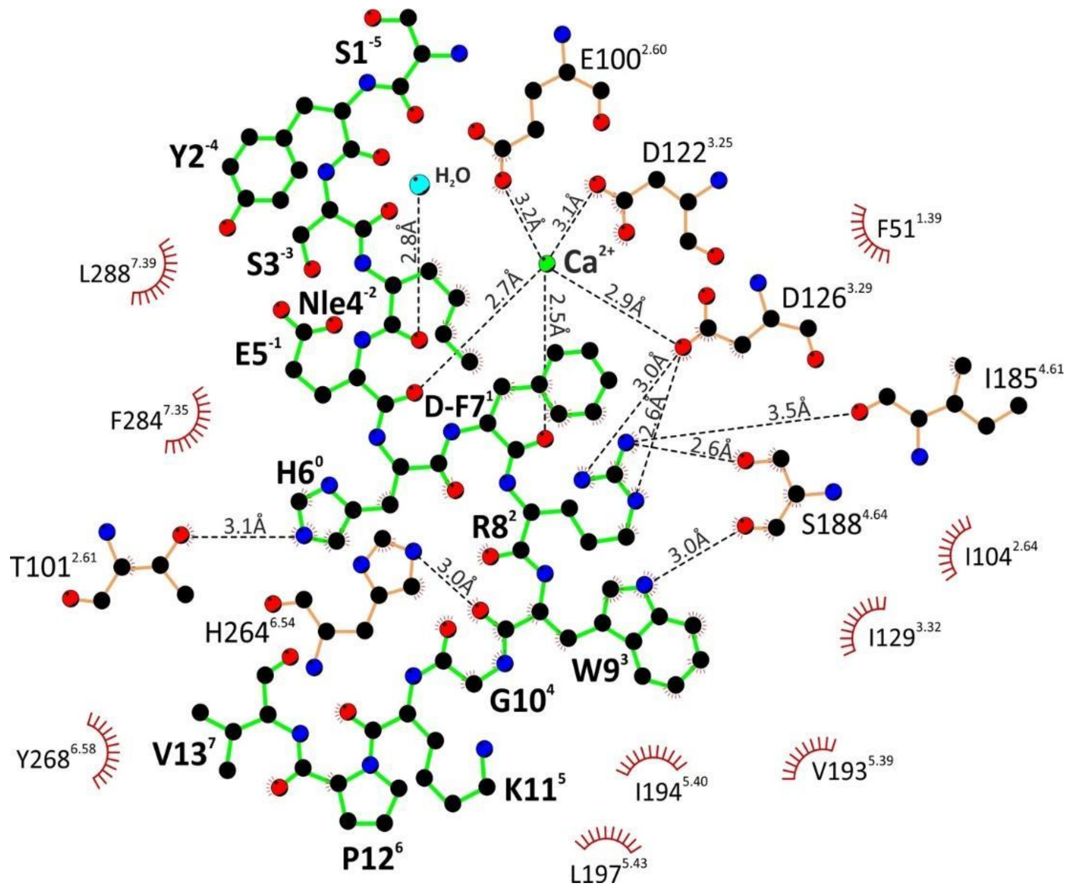


**Figure S9:** ***Interactions between the agonist NDP-α-MSH and the MC4R.*** Potential hydrogen bonds and hydrophobic interactions were analyzed using HBPLUS (McDonald and Thornton, 1994) with a maximum donor acceptor distance of 3.5 Å as implemented in the program LigPlot+ v.2.1 (Laskowski and Swindells, 2011), which was used to draw this schematic view. Potential hydrogen bonds are indicated by black dashed lines and distance labels. Residues with closest distances of less than 3.9 Å are considered to be in van der Waals contact. These residues are shown as dashed, red segments of a circle. Water molecules (cyan color) and the calcium ion (green color) are depicted as spheres, respectively.

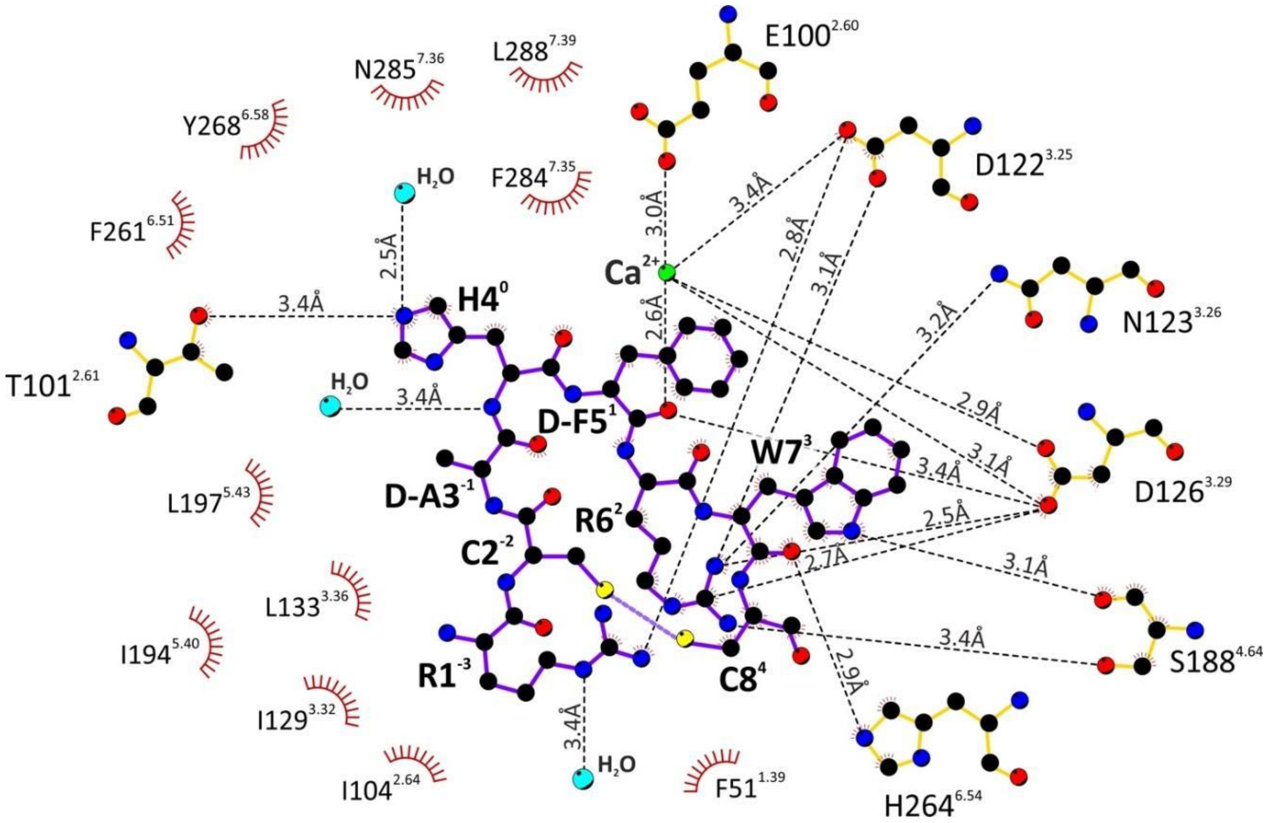


**Figure S10:** ***Interactions between the agonist setmelanotide and MC4R.*** Potential hydrogen bonds and hydrophobic interactions were analyzed using HBPLUS (McDonald and Thornton, 1994) with a maximum donor acceptor distance of 3.5 Å as implemented in the program LigPlot+ v.2.1 (Laskowski and Swindells, 2011), which was used to draw this schematic view. Potential hydrogen bonds are indicated by black dashed lines and distance labels. Residues with closest distances of less than 3.9 Å are regarded to be in van der Waals contact. These residues are shown as dashed, red segments of a circle. Water molecules (cyan color) and the calcium ion (green color) are depicted as spheres, respectively.

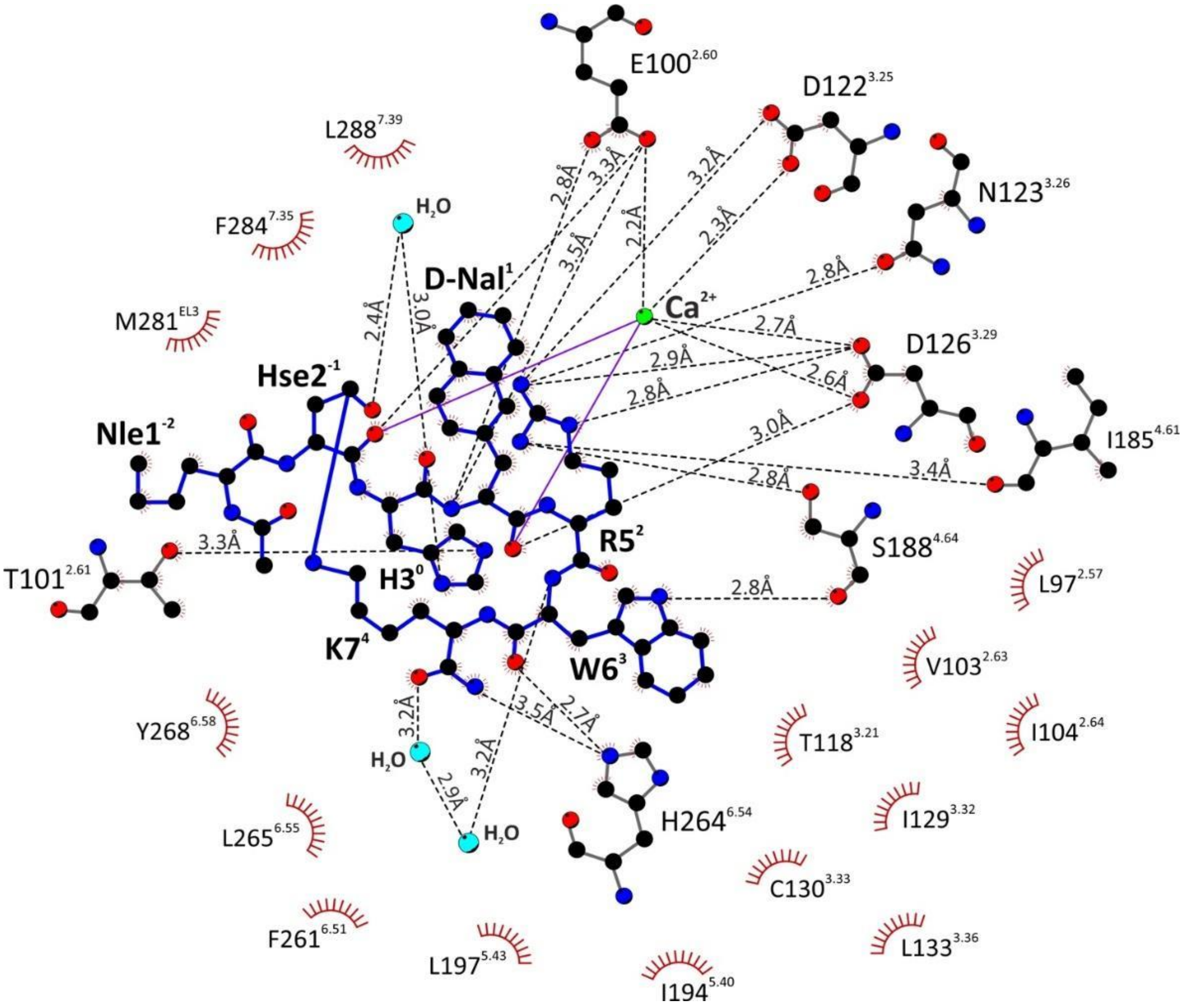


**Figure S11:** ***Interactions between antagonist SHU9119 and MC4R.*** Potential hydrogen bonds and hydrophobic interactions were analyzed using HBPLUS (McDonald and Thornton, 1994) with a maximum donor acceptor distance of 3.5 Å as implemented in the program LigPlot+ v.2.1 (Laskowski and Swindells, 2011), which was used to draw this schematic view. Potential hydrogen bonds are indicated by black dashed lines and distance labels. Residues with closest distances of less than 3.9 Å are deemed to be in van der Waals contact. These residues are shown as dashed, red segments of a circle. Water molecules (cyan color) and the calcium ion (green color) are depicted as spheres, respectively.

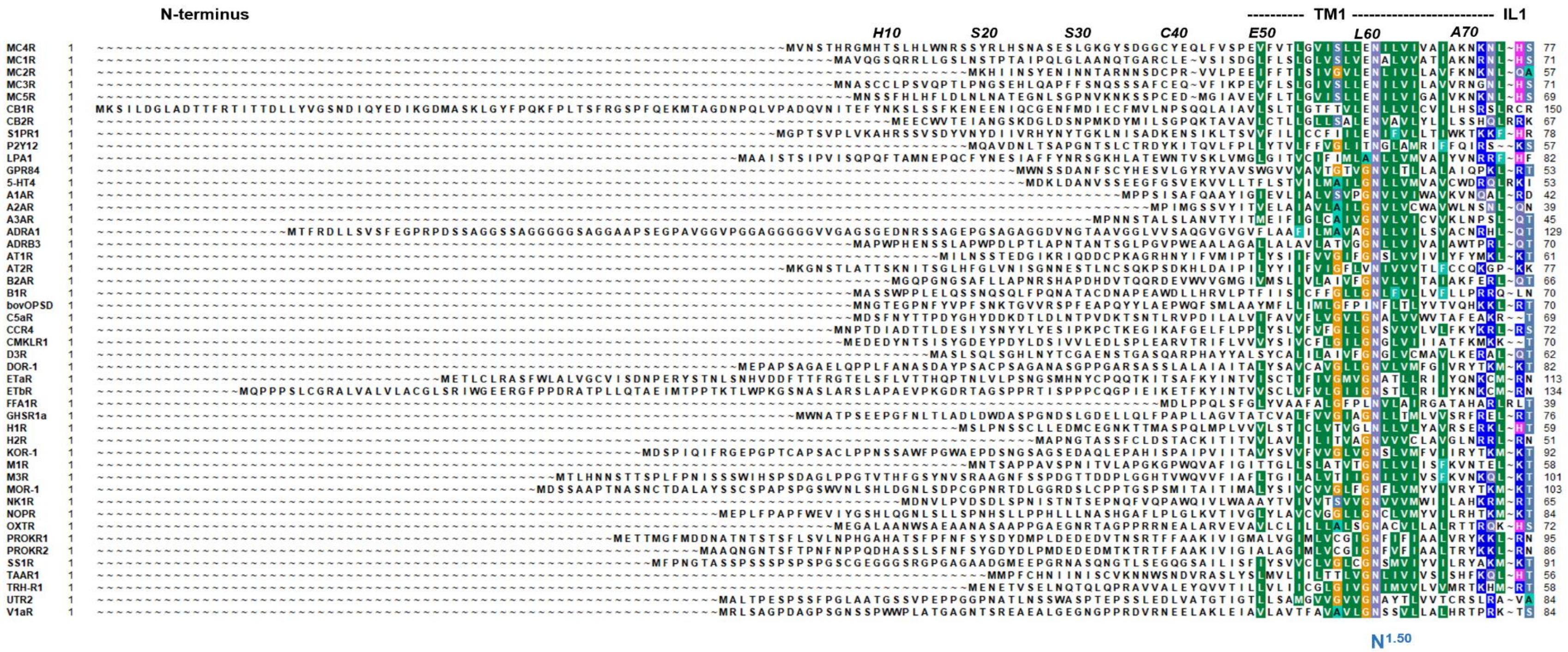

**Figure S12A-E:** ***Sequence alignment of 47 human class A GPCR***. Highly conserved positions according to the unifying *Ballesteros and Weinstein* numbering scheme (Ballesteros and Weinstein, 1995) are indicated by respective numbers below the alignment. The structural dimensions of each MC4R transmembrane helix (TM) or inter-connecting loops (IL = intracellular loop; EL = extracellular loop) are indicated above the sequences, as well as numbers for the MC4R sequence. The alignment was visualized using the software BioEdit (Hall, 1999). Specific background colors indicating conservation (Blossum62 matrix) and reflecting chemical properties of the amino acid side chains: black, proline; blue, positively charged; cyan/green, aromatic and hydrophobic; green, hydrophobic; red, negatively charged; gray, hydrophilic; dark red, cysteines; and magenta, histidine.

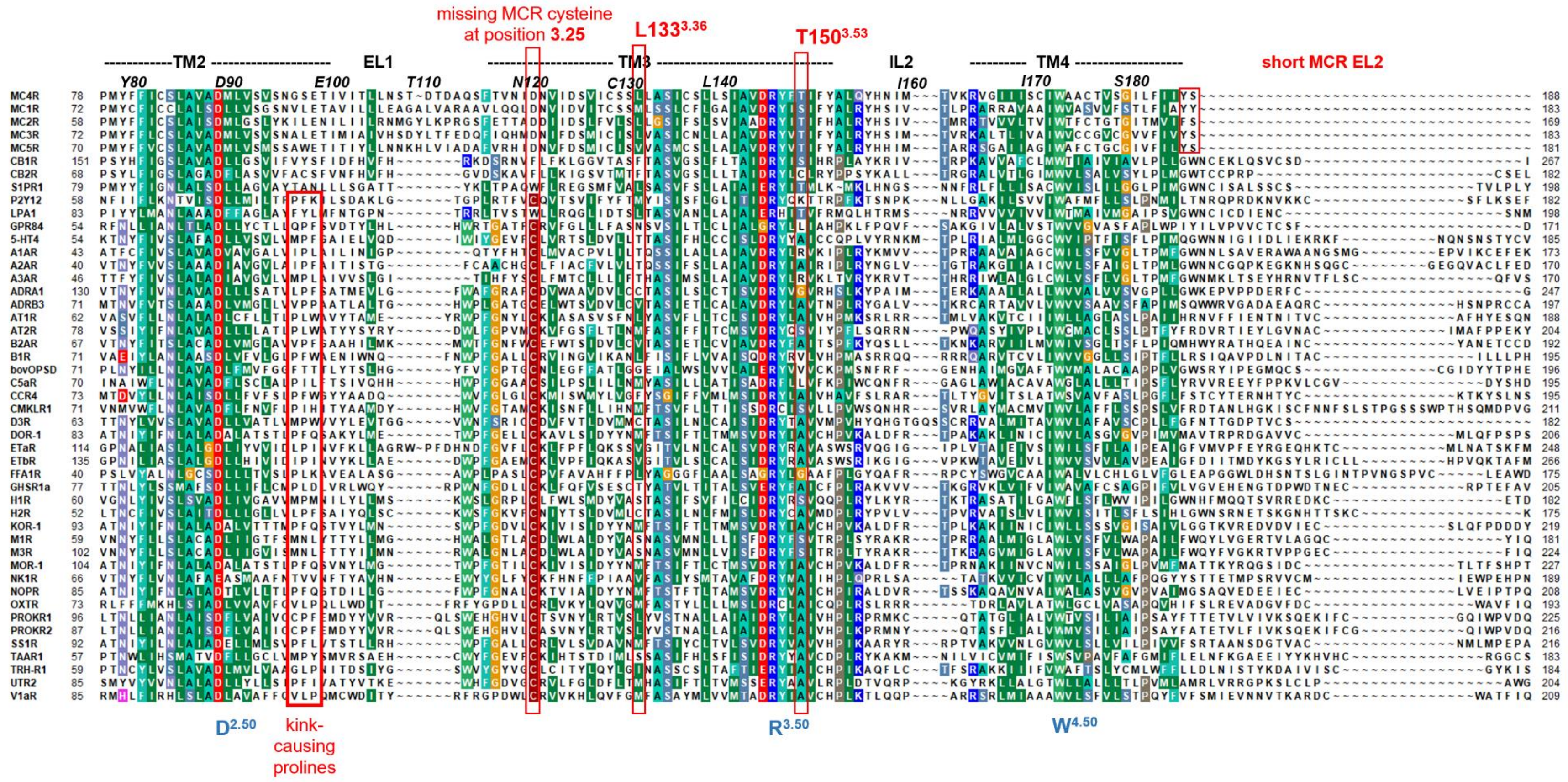
missing MCR cysteine at position 3.25
L133 3.36
T150 3.53
TM2
EL1
TM3
IL2
TM4
short MCR EL2
D2.50
kink-causing prolines
R3.50
W4.50

**Figure S12B**

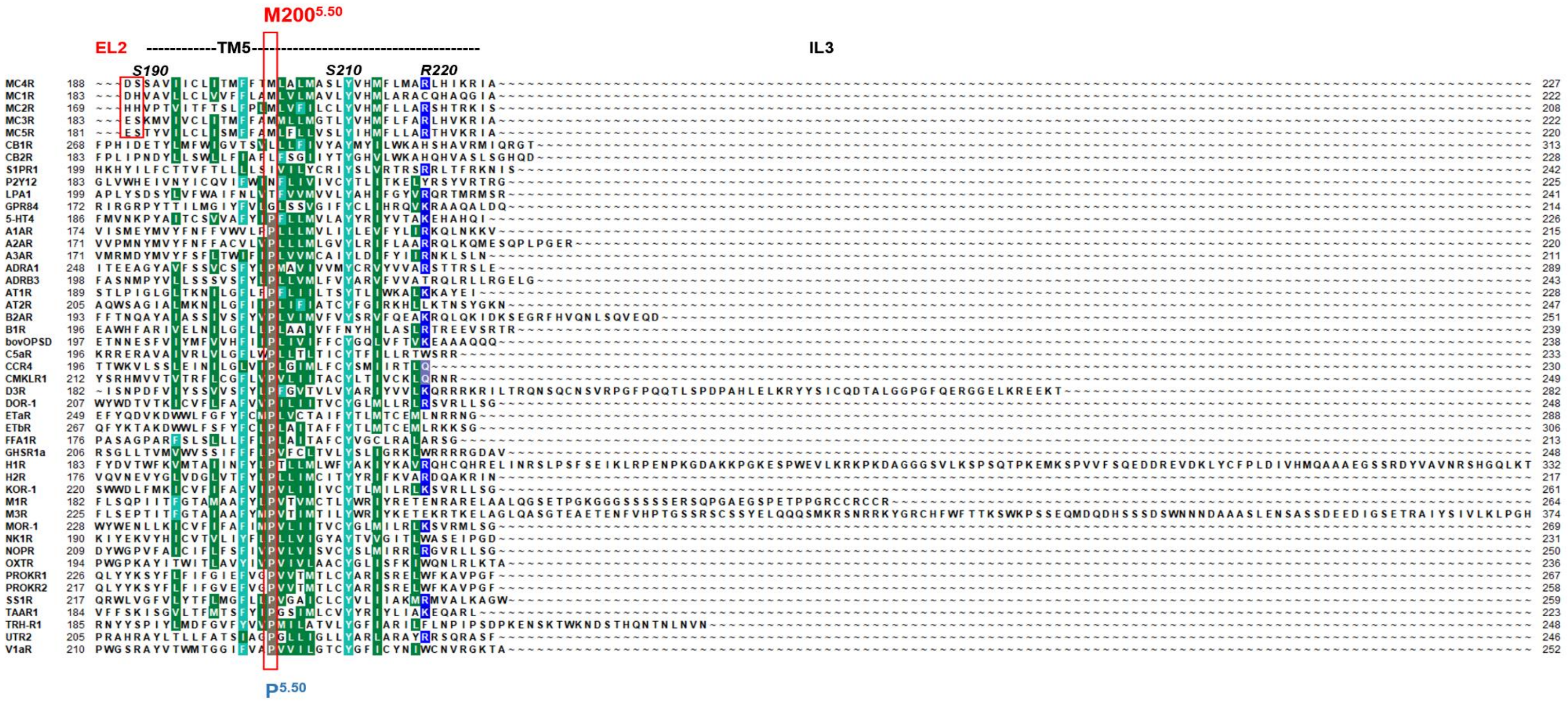


**Figure S12C**

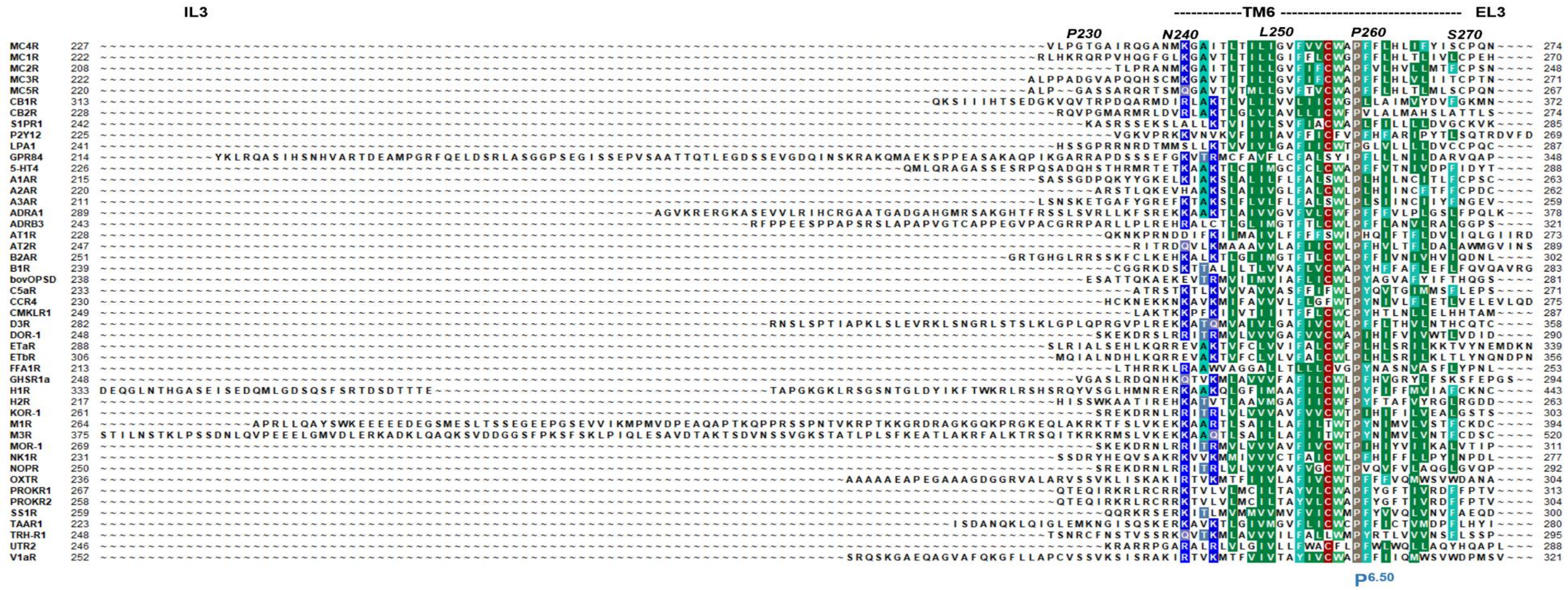


Figure S12D

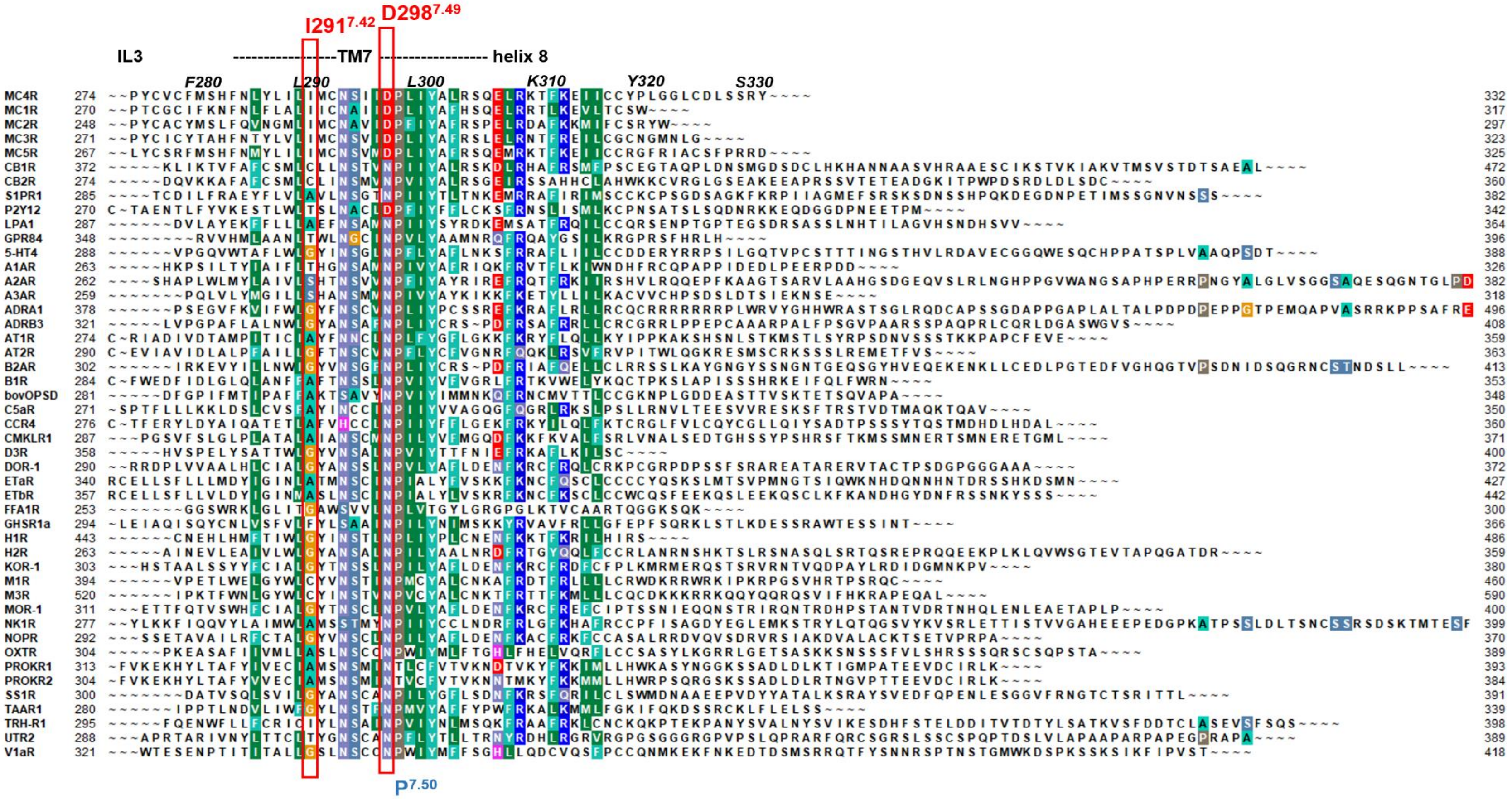
I291 7.42
D298 7.49
IL3
TM7
helix 8
F280
L290
L300
K310
Y320
S330
P7.50

Figure S12E

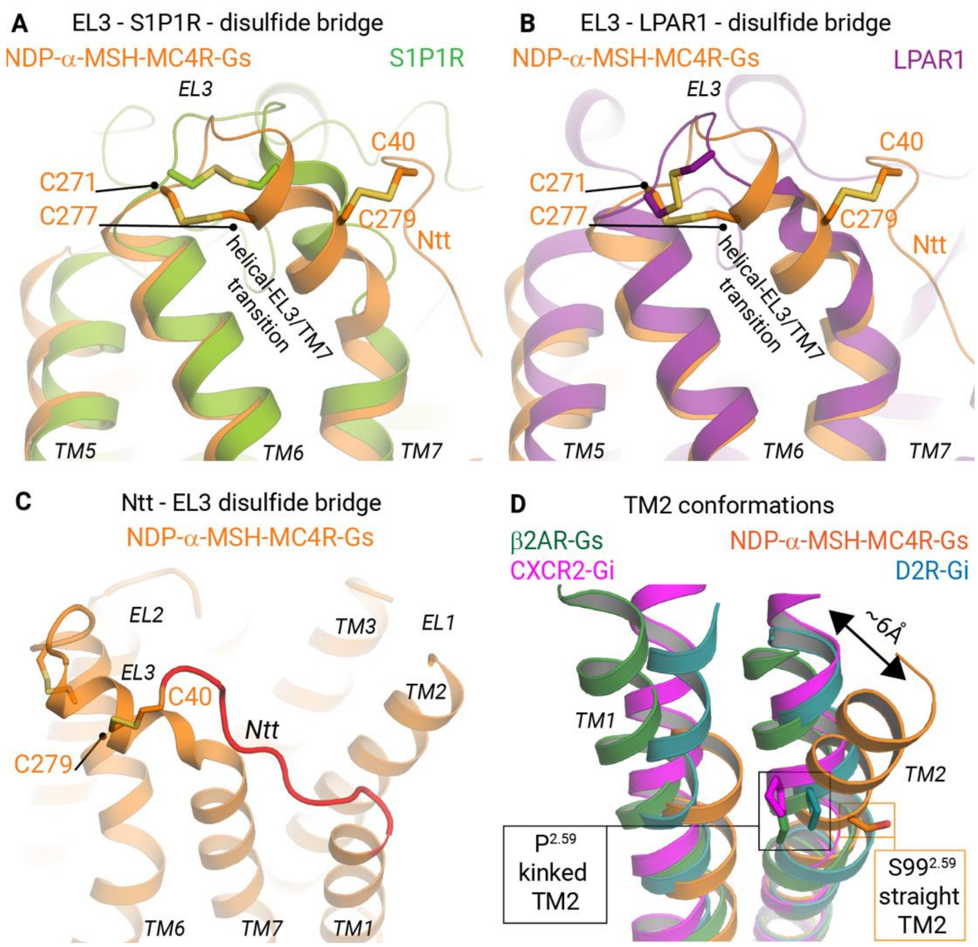


**Figure S13:** ***Specificities of the MC4R in the extracellular region.*** The active state MC4R structure (orange color) shares with

(**A**) the sphingosine 1-phosphate receptor 1 (S1P1R, light green color, PDB ID: 3v2y (Hanson et al., 2012)) and

(**B**) the lysophosphatidic acid receptor 1 (LPAR1, deep purple color, PDB ID: 4z34 (Chrencik et al., 2015) a disulfide bridge in the EL3, which is involved in forming a specific conformation including a helical transition to TM7.

(**C**) The MC4R has a second disulfide bridge between EL3 and the receptor N-terminus (Ntt) that constraints the extracellular helix close to TM7 and by this participates in forming the large peptide ligand-binding region.

(**D**) Contrary to most other class A GPCRs, MCRs have no proline in TM2 (fig. S12B) which usually causes a kink bulge and slight rotation of TM2 toward the extracellular region. The superposition of the MC4R structure in the active state with various already determined GPCR structures (β2AR, forest green color, PDB ID: 3sn6 (Rasmussen et al., 2011); D2R, blue color, PDB ID: 6vms (Yin et al., 2020); CXCR2, magenta color, PDB ID: 6lfo (Liu et al., 2020)) illustrates the difference in spatial TM2 allocation (5.9 Å distance measured between position 2.65 in MC4R and β2AR).

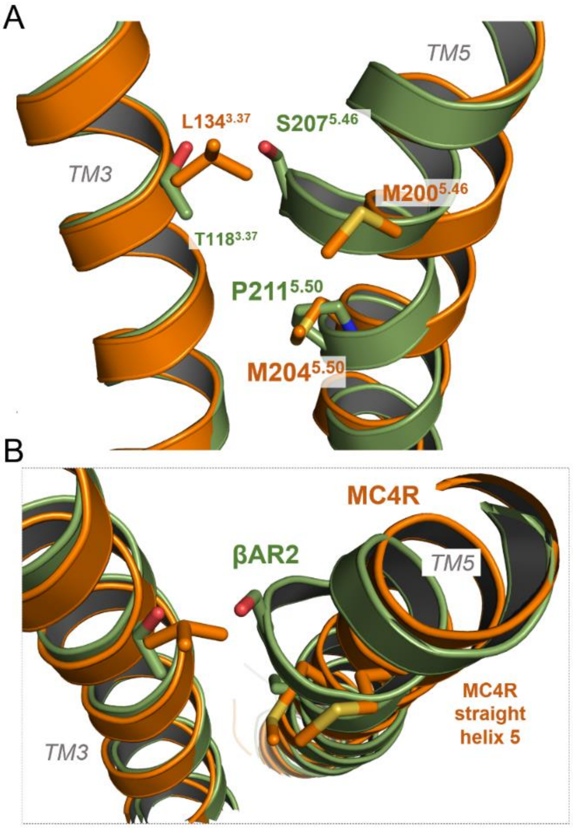


**Figure S14:** ***Straight MC4R helix 5 conformation without a proline induced kink.***
(**A**) Comparison between MC4R (orange) and the β2AR (green, PDB ID: 3sn6) structures reveal differences in the TM5 helix conformation. Caused by a highly conserved proline in class A GPCRs (fig. S12C) TM5 is usually kinked as observed also in the β2AR. In consequence, amino acids at the proline induced kink-bulge pointing to the ligand binding region and forming a specific interface with TM3 **(A)**. S207$^{5.46}$ in the bulge of the β2AR−TM5 is known to interact with ligands and is important for agonistic versus antagonistic effects. This structural helix-specificity is different in MC4R because of a methionine at position 5.50 instead of a proline. The helix is regular without any bulge or kink. This contributes to forming a more distant interface to TM3
**(B)** but also corresponding amino acids are located at spatially distinct positions compared to other class A GPCRs, e.g. position 5.46 **(A)**.

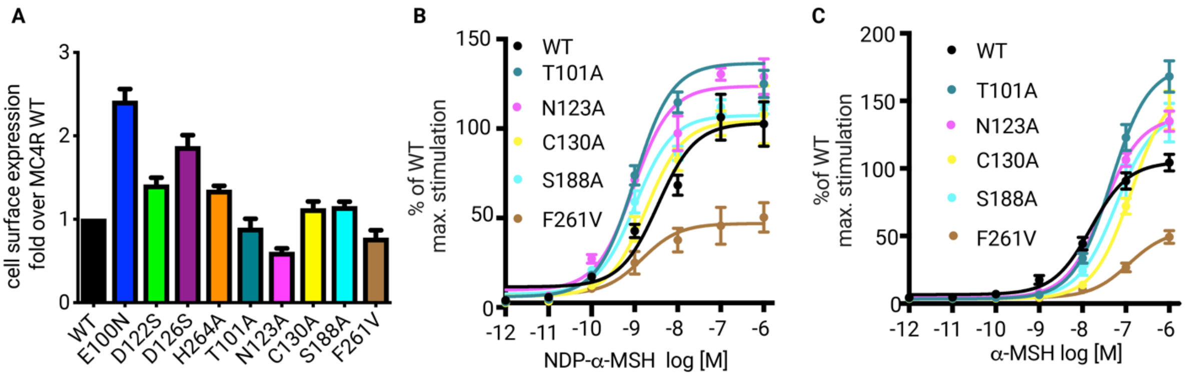


**Figure S15:** ***Functional data of amino acid substitutions in the ligand binding region of MC4R.***

(**A**) Cell surface expression of wild-type (WT) MC4R and mutants shown as fold change over WT.

(**B**) Concentration-response curves of Gs-protein signaling determined as cAMP accumulation of WT and indicated mutants after stimulation by NDP-α-MSH shown as fold change of WT max. signaling [%].

(**C**) Concentration-response curves of Gs-protein signaling determined as cAMP accumulation of WT and indicated mutants after challenge of α-MSH shown as fold of WT max. signaling [%].

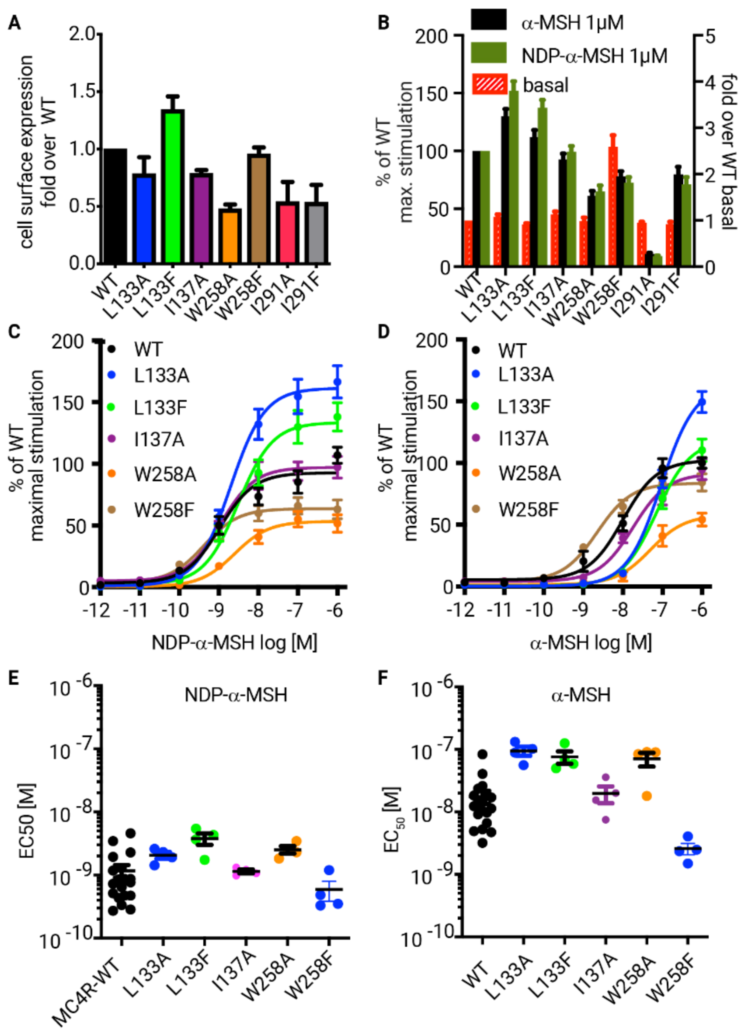


**Figure S16:** ***Functional data of amino acid substitutions in the transmembrane region of MC4R.***

(**A**) Cell surface expression of wild-type MC4R (WT) and mutants shown as fold over WT.

(**B**) Maximal Gs-protein signaling determined as cAMP accumulation of MC4R WT and mutants after addition of 1 µM α-MSH or NDP-α-MSH indicated as fold of MC4R WT max. signaling [%] (left y-axis). Basal cAMP accumulation of MC4R WT and mutants are depicted as fold over WT basal (right y-axis).

(**C**) Concentration-response curves of Gs-protein signaling determined as cAMP accumulation of MC4R WT and indicated mutants after challenge of NDP-α-MSH shown as fold of MC4R WT max. signaling [%].

(**D**) Concentration-response curves of Gs-protein signaling determined as cAMP accumulation of WT and indicated mutants after addition of α-MSH shown as fold of MC4R WT max. signaling [%].

(**E**) $EC_{50}$ values [M] of NDP-α-MSH and

(**F**) $EC_{50}$ values [M] α-MSH-induced signaling calculated from concentration response curves in (**C**) and (**D**), respectively.

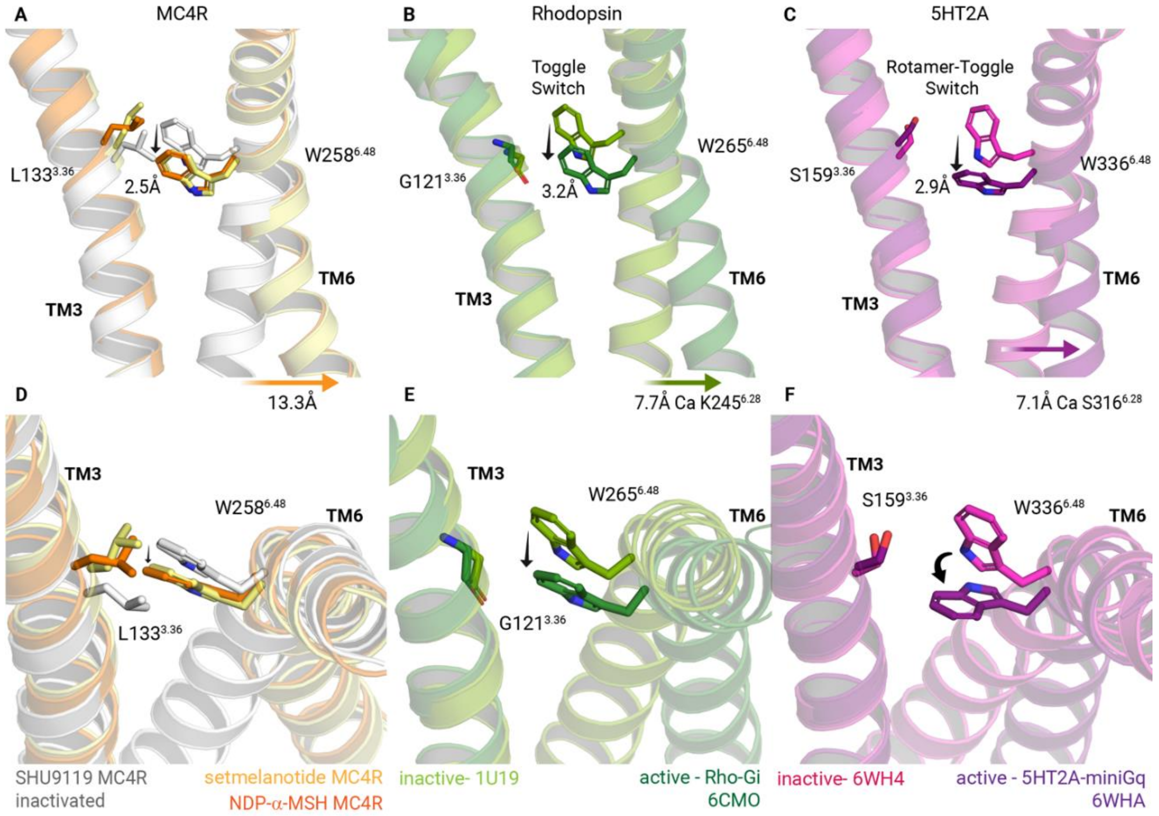


**Figure S17:** ***Localization of $W^{6.48}$ in different active states of various class A GPCRs.*** Visualization of distance differences between inactive and active state conformations of $W^{6.48}$ in

(**A**) MC4R,

(**B**) bovine rhodopsin (PDB IDs: inactive- 1u19 (Okada et al., 2004), active- 6cmo (Kang et al., 2018)), and

(**C**) 5-$HT_{2A}$-receptor (5HT2A) (PDB IDs: inactive- 6wh4, active- 6wha (Kim et al., 2020)). The distances are associated with receptor activation (toggle switch model), thereby the tryptophan undergoes a vertical-lateral shift (measured at NE1 of Trp). In the 5HT2A structure the tryptophan side chain undergoes additional strong rotation. The associated TM6 outward movements are measured at Cα positions of intracellularly located (**A**) $M241^{6.31}$, (**B**) $K245^{6.28}$ and (**C**) $S316^{6.28}$, and reflects the largest spatial difference of TM6 between respective active states.

(**D-F**) Visualization of the horizontal $W^{6.48}$ shift from the top-view. The amino acid position 3.36 at TM3 in opposite to position 6.48 is different between all three receptors. However, in MC4R the $L133^{3.36}$ with a large hydrophobic side chain has a direct impact on $W^{6.48}$. In the SHU9119 antagonized MC4R structure (Yu et al., 2020) this leucine inhibits an activation related shift of TM6 at position $W^{6.48}$.

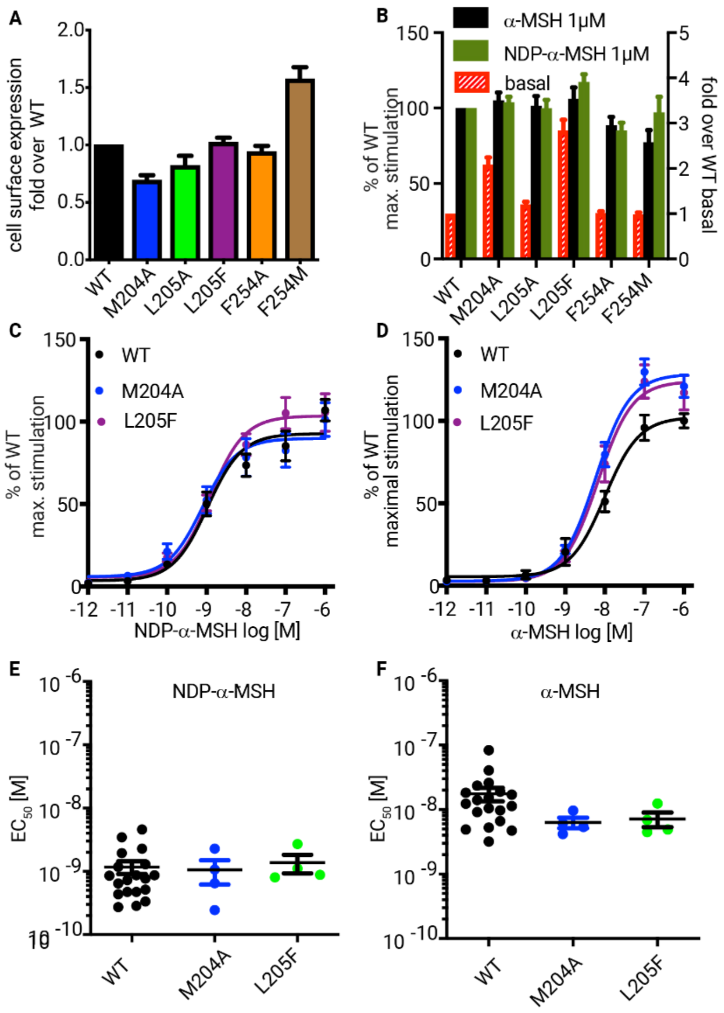


**Figure S18:** ***Functional data of amino acid substitutions in the (extended) "MIF-motif" of MC4R.***

(**A**) Cell surface expression of wild-type MC4R (WT) and mutants shown as fold over WT.

(**B**) Maximal Gs-protein signaling determined as cAMP accumulation of MC4R WT and mutants after addition with 1 µM α-MSH or NDP-α-MSH indicated as fold of WT max. signaling (left y-axis) [%]. Basal cAMP accumulation of MC4R WT and mutants are depicted as fold over WT basal (right y-axis).

(**C**) Concentration-response curves of Gs-protein signaling determined as cAMP accumulation of MC4R WT and indicated mutants after addition of NDP-α-MSH shown as fold of WT max. signaling [%].

(**D**) Concentration-response curves of Gs signaling determined as cAMP accumulation of MC4R WT and indicated mutants after addition of α-MSH shown as fold of WT max. signaling [%].

(**E**) $EC_{50}$ values [M] of NDP-α-MSH and

(**F**) $EC_{50}$ values [M] α-MSH induced signaling calculated from concentration response curves in (**C**) and (**D**), respectively.

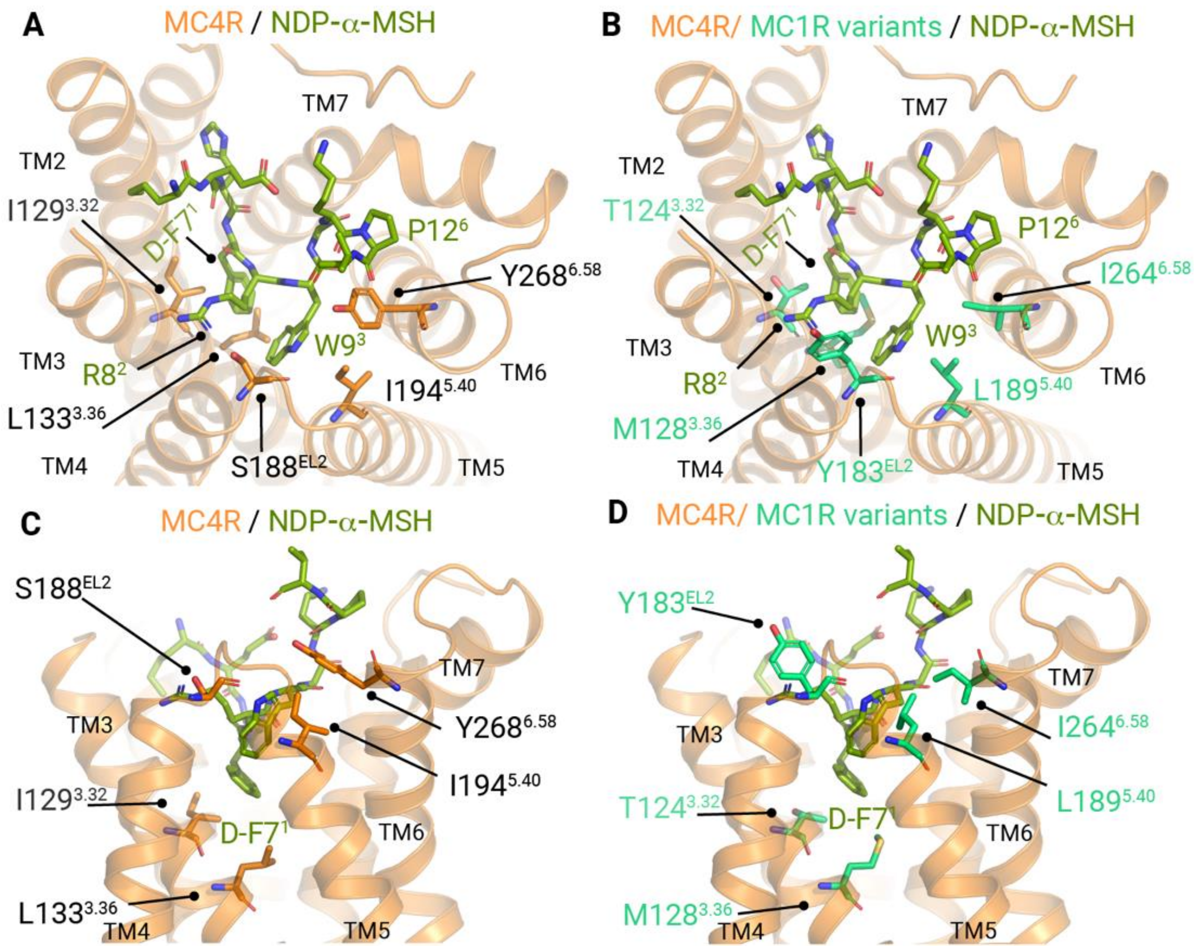


**Figure S19:** ***MC1R amino acid differences in the extended ligand binding pocket based on the NDP-α-MSH−MC4R−Gs−Nb35 complex structure.*** Variations of MC1R compared to MC4R forming the NDP-α-MSH binding site were *in silico* substituted at the NDP-α-MSH−MC4R−Gs−Nb35 complex structure. Ribbon representations, top-view (**A**/**B**) and side view (**C**/**D**), of the ligand binding pocket of NDP-α-MSH (green color) bound to MC4R (orange color) with stick representation of amino acid residues $I129^{3.32}$, $L133^{3.36}$, $S188^{EL2}$, $I194^{5.40}$ and $Y268^{6.58}$, which are different between MC4R and MC1R.

In (**B**/**D**) aforementioned residues were substituted with the corresponding residues at MC1R (green/cyan color), namely, $T124^{3.32}$, $M128^{3.36}$, $Y183^{EL2}$, $L189^{5.40}$ and $I264^{6.58}$. Of note, $S188^{EL2}$ in MC4R EL2 corresponds to $Y183^{EL2}$ in MC1R. The backbone carbonyl group of MC4R $S188^{EL2}$ interacts with the $W9^3$ and the side chain hydroxyl group of $S188^{EL2}$ with $R8^2$ of the agonist NDP-α-MSH.

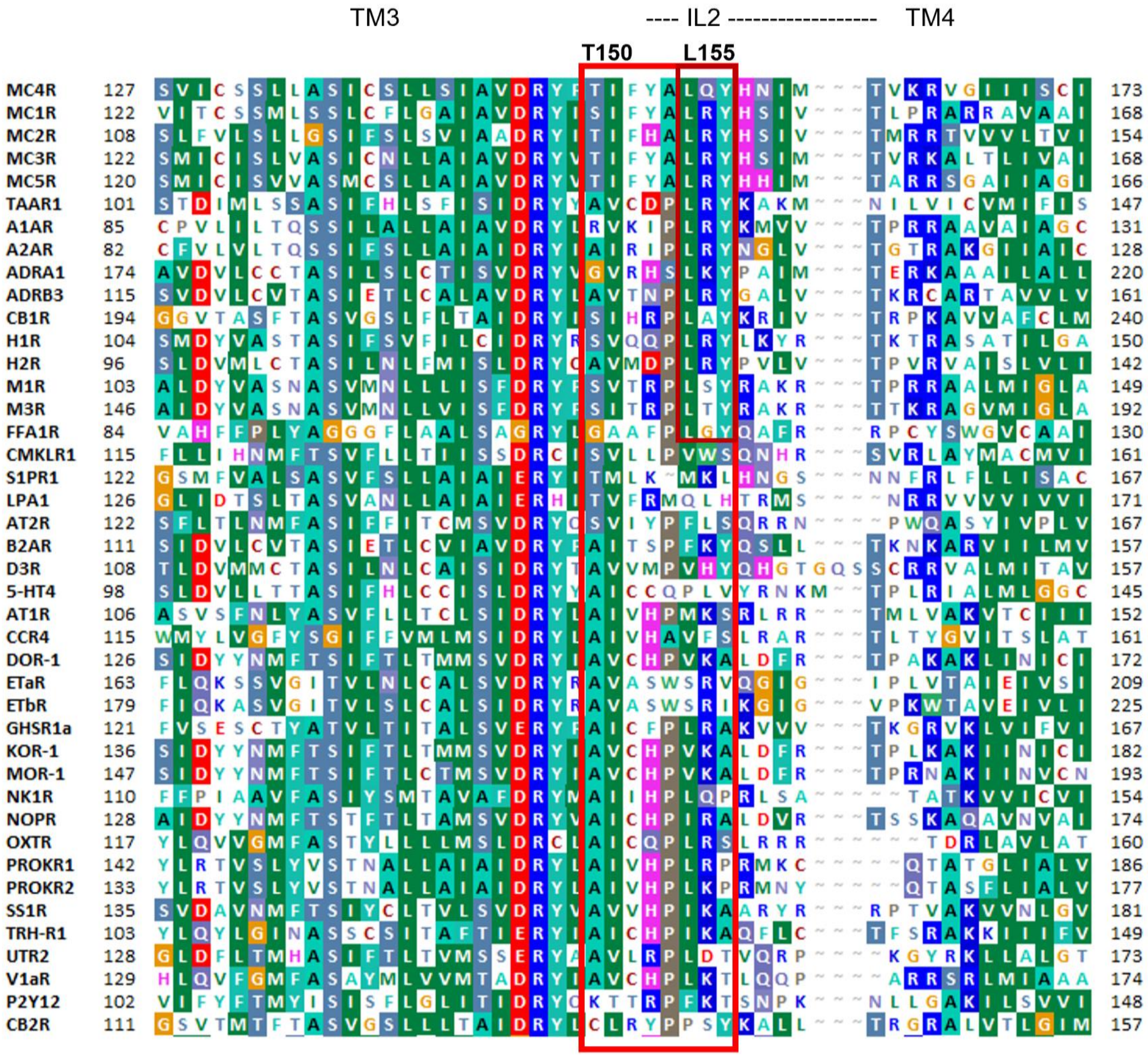


**Figure S20:** ***Intracellular loop two (IL2) amino acids in various class A GPCRs.*** The structural dimension of the MC4R IL2 is indicated above respective sequence, as well as the in the main text discussed specific residues T150$^{3.53}$ (TM3) and L155$^{IL2}$. The alignment was visualized using the software BioEdit (*4*). Background colors indicating conservation (Blossum62 matrix) among different receptors and reflecting chemical properties of the amino acid side chains: black - proline; blue - positively charged; cyan/green - aromatic and hydrophobic; green - hydrophobic; red - negatively charged; gray - hydrophilic; dark red - cysteines; and magenta - histidine.

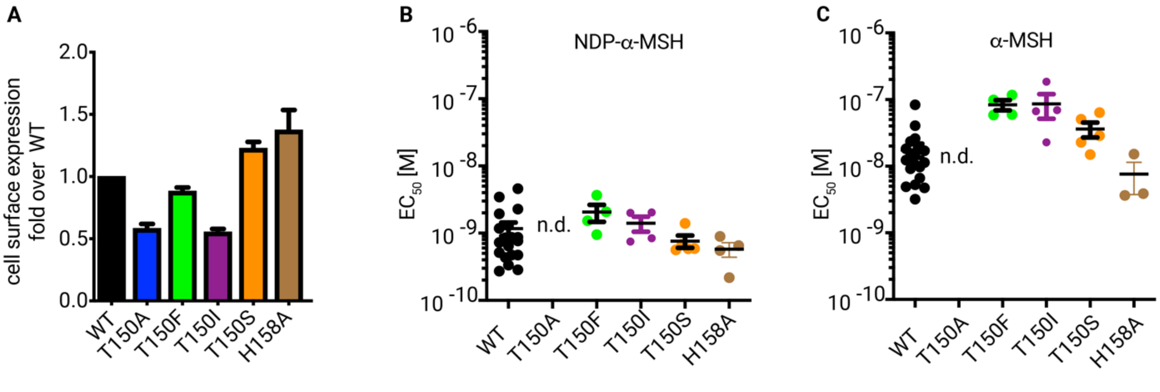


**Figure S21:** ***Signaling of receptor variants modified in the Gs-protein binding region.***

(**A**) Cell surface expression of wild-type MC4R (WT) and mutants shown as fold over WT.

(**B**) $EC_{50}$ values [M] of NDP-α-MSH and

(**C**) $EC_{50}$ values [M] α-MSH induced signaling calculated from concentration response curves.

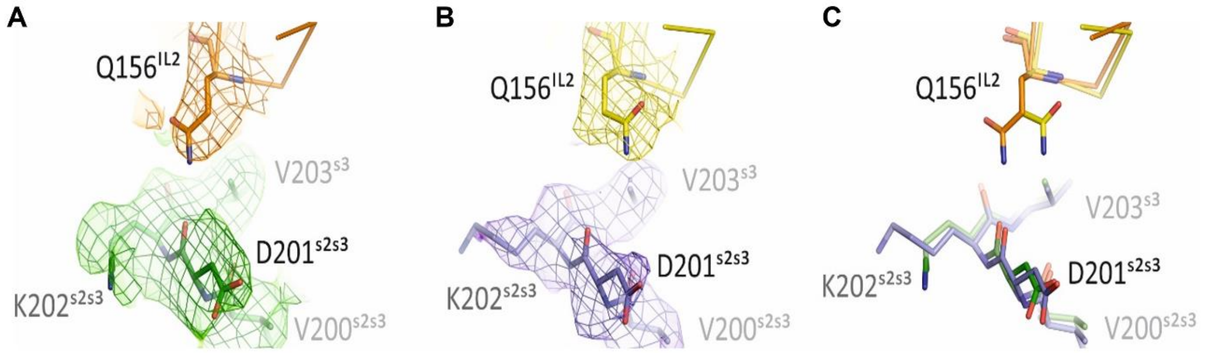


**Figure S22:** ***Cryo-EM map differences of the NDP-α-MSH−MC4R−Gs−Nb35 and setmelanotide-MC4R−Gs−Nb35 complexes at Q156$^{IL2}$ in the IL2−Gs interface.***

**(A)** Close-up view on the cryo-EM maps for Q156$^{IL2}$ in the NDP-α-MSH−MC4R−Gs−NB35 and

**(B)** setmelanotide−MC4R−Gs−Nb35 complex and

**(C)** the superposition of both complexes.

By comparison, both agonist-bound MC4R complexes display two different rotamers for Q156$^{IL2}$, which results in the case of the NDP-α-MSH−MC4R−Gs−Nb35 complex in a hydrogen bond distance between Q156$^{IL2}$ and D201$^{s2s3}$. This subtle difference among setmelanotide and NDP-α-MSH activated MC4R indicates that MC4R agonists modulate G-protein binding at IL2.

NDP-α-MSH−MC4R, the corresponding Gs-protein, setmelanotide−MC4R and its Gs-protein are colored orange, dark green, yellow and slate, respectively. Amino acids are shown as sticks. The protein is depicted as ribbon. Cryo-EM maps are displayed as mesh and volume, contoured at 4 σ level and colored corresponding to the displayed proteins.

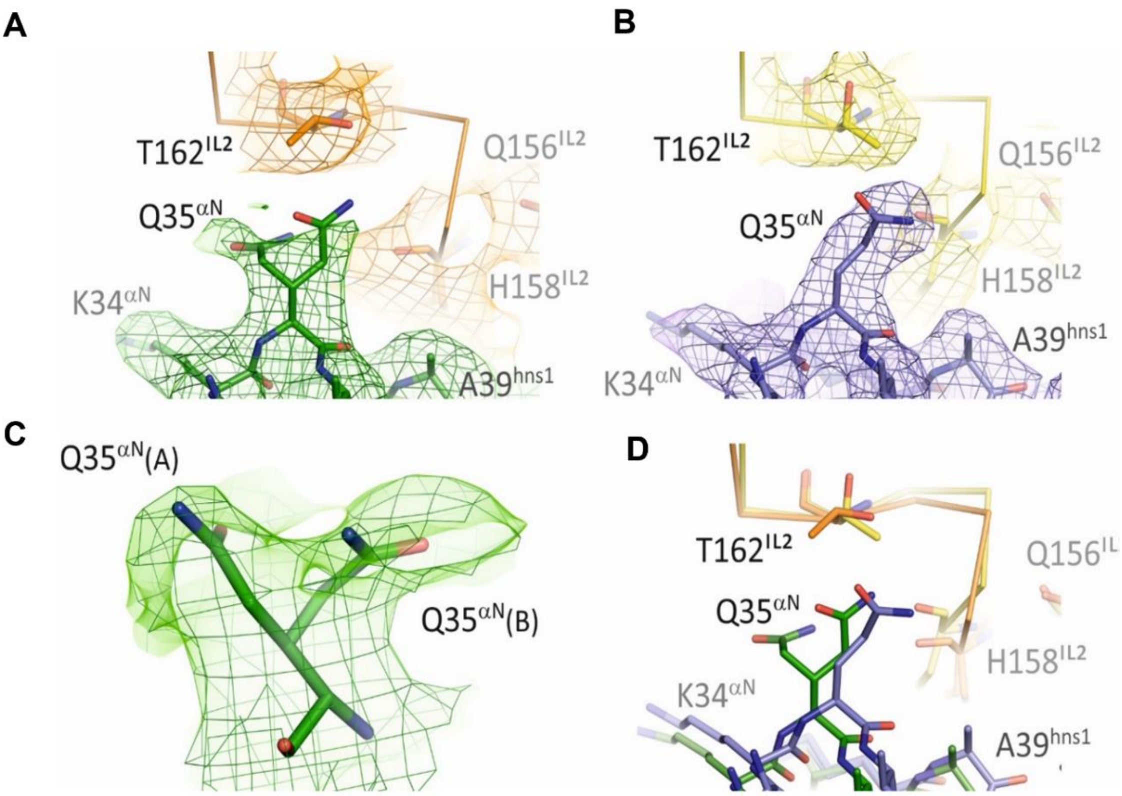


**Figure S23:** ***Cryo-EM map differences of the NDP-α-MSH−MC4R−Gs−Nb35 and setmelanotide-MC4R−Gs−Nb35 complexes at T162$^{IL2}$ in the IL2−Gs interface.***

**(A)** Close-up view on the cryo-EM maps for the amino acid T162$^{IL2}$ in the NDP-α-MSH−MC4R−Gs−Nb35 and

**(B)** setmelanotide−MC4R−Gs−Nb35 complex and

**(D)** the superposition of both complexes.

At T162$^{IL2}$ different rotamers were observed for both agonist-bound complexes, whereby only in the NDP-α-MSH−MC4R−Gs−Nb35 complex the hydroxyl group of T162$^{IL2}$ is in hydrogen bond distance to Nε2-atom of Q35$^{\alpha N}$.

NDP-α-MSH−MC4R, the corresponding Gs-protein, setmelanotide−MC4R and its Gs-protein are colored orange, dark green, yellow and slate, respectively.

Amino acids are depicted in stick representation. The protein is visualized as ribbon.

Cryo-EM maps are displayed as mesh and volume.

**(C)** In the upper row, the cryo-EM maps are contoured at 4 σ level. To verify the double conformation of Q35$^{\alpha N}$, the contour level is set to 2 σ level. All cryo-EM maps are colored corresponding to the displayed proteins.

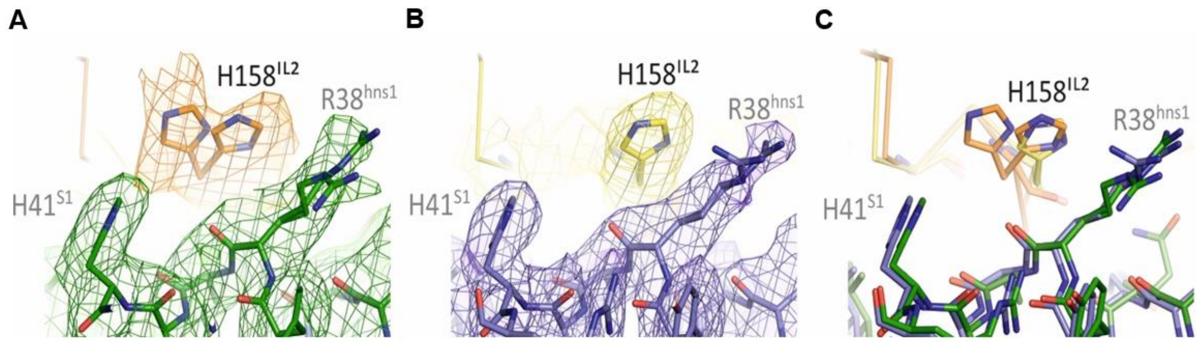


**Figure S24*: Cryo-EM map differences of the NDP-α-MSH−MC4R−Gs−Nb35 and setmelanotide-MC4R−Gs−Nb35 complexes at H158$^{IL2}$ in the IL2−Gs interface.***

**(A)** The close-up view reveals for H158$^{IL2}$ a double conformation in the NDP-α-MSH−MC4R−Gs−Nb35 complex

**(B)** but not for the setmelanotide−MC4R−Gs−Nb35 complex.

**(C)** The superposition of both complexes underlines the structural differences between the two activated MC4R and point out that MC4R agonists effect the IL2 conformation.

NDP-α-MSH–MC4R, the corresponding Gs-protein, setmelanotide–MC4R and its Gs-protein are colored orange, dark green, yellow and slate, respectively. Amino acids are visualized as sticks and the protein as ribbon. Cryo-EM maps are displayed as mesh and volume, contoured at 4 σ level and colored corresponding to the displayed proteins.

### SUPPLEMENTAL TABLES

#### Table S1. Data collection, processing and refinement statistics

| | **NDP-α-MSH-MC4R-$G_s\alpha\beta\gamma$-Nb35 complex** | **Setmelanotide-MC4R-$G_s\alpha\beta\gamma$-Nb35 complex** |
|---|---|---|
| Magnification | 96000 | |
| Voltage (kV) | 300 | |
| Electron exposure ($e^-$/ Å²) | 40 | |
| Defocus range (μm) | -0.8 / -2.0 | |
| Pixel size (Å) | 0.832 | |
| Symmetry imposed | *C1* | |
| Micrographs used (total) | 5403 (5618) | 6979 (7583) |
| Initial particle image (no.) | 2746119 | 4330500 |
| Final particle images (no.) | 221682 | 370621 |
| Map resolution (Å) | 2.86 | 2.58 |
| FSC threshold | 0.143 | 0.143 |
| **Refinement** | | |
| Initial model used (PDB code) | 6W25 / 3SN6 | NDP-MC4R-$G_s\alpha\beta\gamma$-Nb35 |
| Model resolution (Å) | 2.88 | 2.60 |
| Model resolution range (Å) | 233 – 2.88 | 233 – 2.60 |
| Map sharpening B factor (Å²) | -99 | -36 |
| **Model composition** | | |
| Total atoms | 7814 | 7788 |
| Water | 115 | 88 |
| Protein atoms | 7696 | 7696 |
| **B-factors (Å²)** | | |
| Overall | 69.37 | 48.11 |
| R.m.s. deviations bond length (Å) | 0.013 | 0.013 |
| R.m.s. deviations bond angle (°) | 1.705 | 1.707 |
| **Validation** | | |
| Molprobity score | 1.54 | 1.42 |
| Clash score | 3.53 | 3.39 |
| Poor rotamers (%) | 0.86 | 0.97 |
| **Ramachandran plot** | | |
| Favoured (%) | 94.09 | 95.77 |
| Allowed (%) | 5.70 | 4.13 |
| Disallowed (%) | 0.21 | 0.11 |

**Table S2. Contact distances between MC4R and NDP-α-MSH.** Interactions between the binding peptide and the MC4R were analyzed using CONTACT, a program of the CCP4 software suite (Winn et al., 2011). Interactions were calculated between any atoms of the peptide and the receptor, water molecules as well as the calcium ion with a maximum distance of 3.9 Å. Potential hydrogen bonds with a maximum distance of 3.5 Å are highlighted bold. D-phenylalanine and norleucine are abbreviated as three letter codes Dpn and Nle, respectively.

| **Source** | **Number/ Chain** | **Atom** | **Target** | **Number/ Chain** | **Atom** | **Distance [Å]** |
|---|---|---|---|---|---|---|
| Phe | 51R | CD2 | His | 6P | CE1 | 3.83 |
| | | | His | 6P | NE2 | 3.31 |
| Phe | 51R | CE2 | His | 6P | CE1 | 3.44 |
| | | | His | 6P | CD2 | 3.69 |
| | | | His | 6P | NE2 | 3.24 |
| Phe | 51R | CZ | His | 6P | CD2 | 3.76 |
| | | | His | 6P | NE2 | 3.81 |
| Glu | 100R | CD | Nle | 4P | CE | 3.76 |
| Glu | 100R | OE2 | Nle | 4P | CE | 3.63 |
| | | | Glu | 5P | O | 3.66 |
| | | | **Ca** | **1F** | **CA** | **3.20** |
| Thr | 101R | CA | His | 6P | NE2 | 3.89 |
| Thr | 101R | OG1 | His | 6P | CD2 | 3.64 |
| | | | **His** | **6P** | **NE2** | **3.06** |
| Ile | 104R | CG2 | Nle | 4P | CE | 3.09 |
| Asp | 122R | CG | Nle | 4P | CB | 3.60 |
| | | | Nle | 4P | CD | 3.62 |
| | | | Nle | 4P | CE | 3.89 |
| Asp | 122R | OD1 | Nle | 4P | CB | 3.77 |
| | | | Nle | 4P | CG | 3.88 |
| | | | Nle | 4P | CD | 2.91 |
| | | | Nle | 4P | CE | 3.14 |
| Asp | 122R | OD2 | Nle | 4P | CB | 2.93 |
| | | | Nle | 4P | CG | 3.87 |
| | | | Nle | 4P | CD | 3.68 |
| | | | Nle | 4P | CE | 3.85 |
| | | | **Ca** | **1F** | **CA** | **3.09** |
| Asp | 126R | CB | Arg | 8P | NH2 | 3.64 |
| Asp | 126R | CG | Arg | 8P | NE | 3.68 |
| | | | Arg | 8P | NH2 | 3.69 |
| | | | Ca | 1F | CA | 3.50 |
| **Asp** | **126R** | **OD1** | **Ca** | **1F** | **CA** | **3.33** |
| Asp | 126R | OD2 | Dpn | 7P | O | 3.78 |
| | | | Arg | 8P | CD | 3.72 |
| | | | **Arg** | **8P** | **NE** | **2.59** |
| | | | Arg | 8P | CZ | 3.19 |
| | | | **Arg** | **8P** | **NH2** | **2.95** |
| | | | **Ca** | **1F** | **CA** | **2.93** |
| Ile | 129R | CG2 | Dpn | 7P | CG | 3.64 |
| | | | Dpn | 7P | CD2 | 3.43 |
| | | | Dpn | 7P | CE2 | 3.64 |
| Ile | 129R | CD1 | Dpn | 7P | CB | 3.68 |

| | | | | | | |
|---|---|---|---|---|---|---|
| **Ile** | **185R** | **O** | **Arg** | **8P** | **NH1** | **3.50** |
| Ile | 185R | CG2 | Arg | 8P | NH2 | 3.39 |
| Ser | 188R | CA | Trp | 9P | CZ2 | 3.86 |
| | | | Trp | 9P | NE1 | 3.76 |
| Ser | 188R | C | Trp | 9P | NE1 | 3.60 |
| Ser | 188R | O | Trp | 9P | CD1 | 3.87 |
| | | | **Trp** | **9P** | **NE1** | **2.95** |
| Ser | 188R | CB | Arg | 8P | NH1 | 3.28 |
| | | | Trp | 9P | CZ2 | 3.81 |
| | | | Trp | 9P | NE1 | 3.75 |
| Ser | 188R | OG | Arg | 8P | CZ | 3.88 |
| | | | **Arg** | **8P** | **NH1** | **2.64** |
| Val | 193R | CG1 | Trp | 9P | CZ2 | 3.82 |
| Ile | 194R | CG1 | Trp | 9P | CD1 | 3.88 |
| | | | Trp | 9P | NE1 | 3.68 |
| Ile | 194R | CD1 | Trp | 9P | CD1 | 3.64 |
| | | | Trp | 9P | NE1 | 3.75 |
| Leu | 197R | CD1 | Trp | 9P | CZ3 | 3.78 |
| | | | Trp | 9P | CH2 | 3.63 |
| His | 264R | CD2 | Trp | 9P | O | 3.71 |
| His | 264R | CE1 | Gly | 10P | O | 3.64 |
| His | 264R | NE2 | Gly | 10P | O | 3.67 |
| | | | **Trp** | **9P** | **O** | **3.03** |
| Tyr | 268R | CD2 | Trp | 9P | O | 3.18 |
| Tyr | 268R | CE2 | Trp | 9P | CB | 3.86 |
| | | | Trp | 9P | CA | 3.73 |
| | | | Trp | 9P | C | 3.76 |
| | | | Lys | 11P | O | 3.27 |
| | | | Trp | 9P | O | 3.25 |
| | | | Pro | 12P | CA | 3.86 |
| | | | Pro | 12P | O | 3.59 |
| Tyr | 268R | CZ | Pro | 12P | O | 3.56 |
| Tyr | 268R | OH | Lys | 11P | O | 3.90 |
| | | | Pro | 12P | O | 3.55 |
| Phe | 284R | CG | Gly | 10P | CA | 3.49 |
| Phe | 284R | CD1 | Gly | 10P | CA | 3.60 |
| Phe | 284R | CD2 | Gly | 10P | CA | 3.43 |
| Phe | 284R | CE1 | Gly | 10P | CA | 3.68 |
| Phe | 284R | CE2 | His | 6P | O | 3.43 |
| | | | Gly | 10P | CA | 3.51 |
| Phe | 284R | CZ | Gly | 10P | CA | 3.63 |
| | | | Arg | 8P | O | 3.43 |
| | | | Gly | 10P | N | 3.78 |
| Leu | 288R | CD1 | His | 6P | CB | 3.55 |
| Leu | 288R | CD2 | Dpn | 7P | CD2 | 3.89 |
| **$Ca^{2+}$** | **1F** | **CA** | **Glu** | **5P** | **O** | **2.68** |
| | | | Glu | 5P | C | 3.88 |
| **$Ca^{2+}$** | **1F** | **CA** | **Dpn** | **7P** | **O** | **2.50** |
| | | | Dpn | 7P | C | 3.72 |
| **Nle** | **4P** | **O** | **water** | **43H** | **O** | **2.78** |

**Table S3. Contact distances between MC4R and the agonist setmelanotide.** Interactions between the binding peptide and the MC4R were analyzed using CONTACT, a program of the CCP4 software suite (Winn et al., 2011). Interactions were calculated between any atoms of the peptide and the receptor, water molecules as well as the calcium ion with a maximum distance of 3.9 Å. Potential hydrogen bonds with a maximum distance of 3.5 Å are highlighted bold. D-phenylalanine and D-alanine are abbreviated as three letter codes Dpn and Dal, respectively. *Amino acids in double conformation (amino acid rotamers A and B).

| Source | Number/ Chain | Atom | Target | Number/ Chain | Atom | Distance [Å] |
|---|---|---|---|---|---|---|
| Phe | 51R | CD2 | His | 4P | CE1 | 3.56 |
| Phe | 51R | CE2 | His | 4P | CE1 | 3.47 |
| | | | His | 4P | NE2 | 3.46 |
| Phe | 51R | CZ | His | 4P | CE1 | 3.74 |
| | | | His | 4P | NE2 | 3.83 |
| | | | His | 4P | ND1 | 3.88 |
| | | | His | 4P | CE1 | 3.56 |
| Glu(A)* | 100R | OE1 | Dpn | 5P | N | 3.57 |
| **Glu(A)*** | **100R** | **OE2** | **Ca** | **1F** | **CA** | **3.02** |
| Glu(B)* | 100R | OE1 | Dal | 3P | O | 3.88 |
| **Glu(B)*** | **100R** | **OE2** | **Ca** | **1F** | **CA** | **2.90** |
| Thr | 101R | CB | His | 4P | CE1 | 3.84 |
| **Thr** | **101R** | **OG1** | **His** | **4P** | **ND1** | **3.39** |
| | | | His | 4P | CE1 | 2.86 |
| Ile | 104R | CG1 | Dal | 3P | O | 3.69 |
| Ile | 104R | CD1 | Dal | 3P | O | 3.89 |
| Asp | 122R | CG | Arg | 1P | NH1 | 3.81 |
| **Asp** | **122R** | **OD1** | **Arg** | **6P** | **NH1** | **3.12** |
| Asp | 122R | OD2 | Arg | 1P | CZ | 3.58 |
| | | | **Arg** | **1P** | **NH1** | **2.80** |
| | | | **Ca** | **1F** | **CA** | **3.35** |
| **Asn** | **123R** | **ND2** | **Arg** | **6P** | **NH1** | **3.16** |
| Asp | 126R | CB | Arg | 6P | NH1 | 3.72 |
| Asp | 126R | CG | Arg | 6P | NE | 3.67 |
| | | | Arg | 6P | NH1 | 3.35 |
| | | | Ca | 1F | CA | 3.34 |
| **Asp** | **126R** | **OD1** | **Ca** | **1F** | **CA** | **2.87** |
| **Asp** | **126R** | **OD2** | **Dpn** | **5P** | **O** | **3.42** |
| | | | **Arg** | **6P** | **NE** | **2.65** |
| | | | **Arg** | **6P** | **NH1** | **2.54** |
| | | | **Ca** | **1F** | **CA** | **3.07** |
| | | | Arg | 6P | CD | 3.88 |
| | | | Arg | 6P | CZ | 2.95 |
| | | | Arg | 6P | CB | 3.88 |
| Ile | 129R | CG2 | Dpn | 5P | CG | 3.61 |
| | | | Dpn | 5P | CD1 | 3.86 |
| | | | Dpn | 5P | CD2 | 3.61 |
| | | | Dpn | 5P | CE2 | 3.88 |
| Ile | 129R | CD1 | Dpn | 5P | CB | 3.82 |
| Leu | 133R | CD1 | Dpn | 5P | CE2 | 3.46 |
| | | | Dpn | 5P | CZ | 3.69 |

| | | | | | | |
|---|---|---|---|---|---|---|
| Ser | 188R | CA | Trp | 7P | NE1 | 3.66 |
| Ser | 188R | C | Trp | 7P | NE1 | 3.62 |
| Ser | 188R | O | Trp | 7P | CD1 | 3.79 |
| | | | **Trp** | **7P** | **NE1** | **3.07** |
| Ser | 188R | CB | Arg | 6P | CD | 3.56 |
| | | | Trp | 7P | NE1 | 3.50 |
| **Ser** | **188R** | **OG** | **Arg** | **6P** | **NH2** | **3.42** |
| Ile | 194R | CD1 | Trp | 7P | CD2 | 3.57 |
| | | | Trp | 7P | CG | 3.68 |
| | | | Trp | 7P | CD1 | 3.63 |
| | | | Trp | 7P | NE1 | 3.52 |
| | | | Trp | 7P | CE2 | 3.48 |
| Leu | 197R | CD1 | Trp | 7P | CH2 | 3.81 |
| Phe | 261R | CZ | Dpn | 5P | CE2 | 3.49 |
| His | 264R | CD2 | Trp | 7P | O | 3.58 |
| His | 264R | CE1 | Cys | 8P | C | 3.80 |
| His | 264R | NE2 | Cys | 8P | CA | 3.67 |
| | | | Cys | 8P | C | 3.65 |
| | | | Trp | 7P | C | 3.90 |
| | | | **Trp** | **7P** | **O** | **2.91** |
| Tyr | 268R | CG | Trp | 7P | O | 3.71 |
| Tyr | 268R | CD1 | Trp | 7P | C | 3.86 |
| | | | Trp | 7P | O | 3.15 |
| Tyr | 268R | CE1 | Trp | 7P | CA | 3.84 |
| | | | Trp | 7P | C | 3.52 |
| | | | Trp | 7P | O | 3.27 |
| Tyr | 268R | CE2 | Trp | 7P | CB | 3.82 |
| Tyr | 268R | CZ | Trp | 7P | CA | 3.69 |
| | | | Trp | 7P | CB | 3.89 |
| | | | Trp | 7P | O | 3.89 |
| Tyr | 268R | OH | Trp | 7P | CA | 3.81 |
| Phe | 284R | CG | Cys | 8P | CA | 3.82 |
| | | | Cys | 8P | CB | 3.76 |
| Phe | 284R | CD2 | His | 4P | O | 3.72 |
| | | | Cys | 8P | CA | 3.88 |
| | | | Cys | 8P | CB | 3.39 |
| Phe | 284R | CE1 | Arg | 6P | O | 3.68 |
| Phe | 284R | CE2 | His | 4P | O | 3.03 |
| | | | Cys | 8P | CB | 3.71 |
| | | | Arg | 6P | O | 3.86 |
| Phe | 284R | CZ | Arg | 6P | N | 3.86 |
| | | | Arg | 6P | O | 3.08 |
| Asn | 285R | ND2 | His | 4P | CD2 | 3.51 |
| Leu | 288R | CD1 | His | 4P | CB | 3.49 |
| **$Ca^{2+}$** | **1F** | **CA** | **Dpn** | **5** | **O** | **2.64** |
| $Ca^{2+}$ | 1F | CA | Dpn | 5 | C | 3.82 |
| **water** | **30H** | **O** | **His** | **4P** | **N** | **3.41** |
| **water** | **62H** | **O** | **His** | **4P** | **NE2** | **2.50** |
| **water** | **63H** | **O** | **Arg** | **1P** | **NE** | **3.44** |

**Table S4. Contact distances between MC4R and the antagonist SHU9119.** Interactions between the binding peptide and the MC4R were analyzed using CONTACT, a program of the CCP4 software suite (Winn et al., 2011). Interactions were calculated between any atoms of the peptide and the receptor, water molecules as well as the calcium ion with a maximum distance of 3.9 Å. Potential hydrogen bonds with a maximum distance of 3.5 Å are highlighted bold. D-Nal corresponds to (2R)-2-amino-3-(naphthalene-2-YL)propanoic acid abbreviated as 4J2 in the protein database (PDB ID: 6w25). Norleucine is shown as Nle in the three letter code. At the second position of the SHU9119 peptide there is a homoserine. In the corresponding pdb file (PDB ID: 6w25), this amino acid is abbreviated as Asp B2.

| Source | Number/ Chain | Atom | Target | Number/ Chain | Atom | Distance [Å] |
|---|---|---|---|---|---|---|
| Leu | 97A | CD2 | DNal | 4B | CB | 3.80 |
| Glu | 100A | CD | Asp | 2B | O | 3.77 |
| | | | His | 3B | CA | 3.86 |
| | | | DNal | 4B | N | 3.45 |
| | | | Ca | 2101A | CA | 3.37 |
| Glu | 100A | OE1 | His | 3B | CA | 3.45 |
| | | | His | 3B | C | 3.56 |
| | | | **DNal** | **4B** | **N** | **2.78** |
| | | | DNal | 4B | CA | 3.71 |
| | | | DNal | 4B | CB | 3.53 |
| | | | **Ca** | **2101A** | **CA** | **2.21** |
| Glu | 100A | OE2 | DNal | 4B | O | 3.59 |
| | | | **Asp** | **2B** | **O** | **3.25** |
| | | | **DNal** | **4B** | **N** | **3.48** |
| | | | **Ca** | **2101A** | **CA** | **2.21** |
| Thr | 101A | CA | His | 3B | NE2 | 3.76 |
| Thr | 101A | CB | His | 3B | NE2 | 3.71 |
| Thr | 101A | OG1 | His | 3B | CD2 | 3.30 |
| | | | **His** | **3B** | **NE2** | **3.31** |
| Thr | 101A | CG2 | His | 3B | NE2 | 3.51 |
| Val | 103A | CG1 | Nle | 1B | CE | 3.78 |
| Ile | 104A | CG1 | Nle | 1B | CD | 3.82 |
| Ile | 104A | CD1 | Asp | 2B | C | 3.66 |
| | | | His | 3B | N | 3.52 |
| | | | His | 3B | CG | 3.87 |
| | | | His | 3B | ND1 | 3.80 |
| | | | His | 3B | CE1 | 3.87 |
| Thr | 118A | O | Nle | 1B | CE | 3.77 |
| Asp | 122A | CB | Nle | 1B | CB | 3.84 |
| | | | Nle | 1B | CD | 3.80 |
| | | | Nle | 1B | CE | 3.83 |
| Asp | 122A | CG | Nle | 1B | CB | 3.56 |
| | | | Nle | 1B | CD | 3.85 |
| | | | Ca | 2101A | CA | 3.27 |
| Asp | 122A | OD1 | Nle | 1B | CB | 3.74 |
| | | | Nle | 1B | CD | 3.47 |
| | | | Asp | 2B | O | 3.54 |
| | | | **Ca** | **2101A** | **CA** | **2.27** |

| | | | | | | |
|---|---|---|---|---|---|---|
| Asp | 122A | OD2 | Nle | 1B | CB | 3.78 |
| | | | **Arg** | **5B** | **NH1** | **3.18** |
| | | | Ca | 2101A | CA | 3.65 |
| Asn | 123A | CA | Arg | 5B | NH1 | 3.79 |
| Asn | 123A | CG | Arg | 5B | NH1 | 3.83 |
| Asn | 123A | OD1 | Arg | 5B | NH2 | 3.79 |
| | | | Arg | 5B | CZ | 3.72 |
| | | | **Arg** | **5B** | **NH1** | **2.83** |
| Asp | 126A | O | DNal | 4B | CE3 | 3.56 |
| Asp | 126A | CB | Arg | 5B | NE | 3.52 |
| Asp | 126A | CG | DNal | 4B | O | 3.55 |
| | | | Arg | 5B | NH1 | 3.83 |
| | | | Arg | 5B | NE | 3.38 |
| | | | Ca | 2101A | CA | 3.04 |
| **Asp** | **126A** | **OD1** | **DNal** | **4B** | **O** | **3.02** |
| | | | DNal | 4B | C | 3.61 |
| | | | DNal | 4B | CD2 | 3.60 |
| | | | DNal | 4B | CB | 3.50 |
| | | | **Ca** | **2101A** | **CA** | **2.64** |
| Asp | 126A | OD2 | DNal | 4B | O | 3.54 |
| | | | Arg | 5B | CZ | 3.26 |
| | | | **Arg** | **5B** | **NH1** | **2.85** |
| | | | **Arg** | **5B** | **NE** | **2.80** |
| | | | **Ca** | **2101A** | **CA** | **2.73** |
| Ile | 129A | CG2 | DNal | 4B | CD2 | 3.81 |
| | | | DNal | 4B | CZ1 | 3.58 |
| | | | DNal | 4B | CE2 | 3.65 |
| | | | DNal | 4B | CD1 | 3.82 |
| | | | DNal | 4B | CE1 | 3.67 |
| | | | DNal | 4B | CB | 3.61 |
| | | | DNal | 4B | CG | 3.82 |
| Cys | 130A | CA | DNal | 4B | CE4 | 3.69 |
| Cys | 130A | CB | DNal | 4B | CE4 | 3.50 |
| Cys | 130A | SG | DNal | 4B | CE4 | 3.78 |
| Leu | 133A | CD2 | DNal | 4B | CZ2 | 3.84 |
| | | | DNal | 4B | CZ3 | 3.63 |
| **Ile** | **185A** | **O** | **Arg** | **5B** | **NH2** | **3.36** |
| Ile | 185A | CG2 | Arg | 5B | NH2 | 3.73 |
| | | | Arg | 5B | CZ | 3.61 |
| | | | Arg | 5B | NH1 | 3.72 |
| Ile | 185A | CD1 | Arg | 5B | CD | 3.66 |
| | | | DNal | 4B | CE3 | 3.82 |
| Ser | 188A | C | Trp | 6B | NE1 | 3.54 |
| Ser | 188A | O | Trp | 6B | CD1 | 3.69 |
| | | | **Trp** | **6B** | **NE1** | **2.78** |
| | | | Trp | 6B | CE2 | 3.72 |
| Ser | 188A | CB | Arg | 5B | NH2 | 3.47 |
| | | | Arg | 5B | CD | 3.69 |
| | | | Trp | 6B | NE1 | 3.77 |
| | | | Trp | 6B | CZ2 | 3.85 |
| **Ser** | **188A** | **OG** | **Arg** | **5B** | **NH2** | **2.83** |
| Ile | 194A | CG1 | Trp | 6B | NE1 | 3.65 |
| Ile | 194A | CD1 | Trp | 6B | CD1 | 3.78 |

| | | | | | | |
|---|---|---|---|---|---|---|
| | | | Trp | 6B | NE1 | 3.60 |
| Leu | 197A | CD1 | Trp | 6B | CH2 | 3.68 |
| | | | DNal | 4B | CE1 | 3.62 |
| Phe | 261A | CE2 | DNal | 4B | CZ2 | 3.71 |
| Phe | 261A | CZ | DNal | 4B | CZ1 | 3.65 |
| | | | DNal | 4B | CZ2 | 3.44 |
| | | | DNal | 4B | CE1 | 3.51 |
| His | 264A | CD2 | Trp | 6B | O | 3.28 |
| | | | Nh2 | 8B | N | 3.32 |
| His | 264A | NE2 | Trp | 6B | C | 3.73 |
| | | | Lys | 7B | CA | 3.55 |
| | | | Lys | 7B | C | 3.77 |
| | | | **Trp** | **6B** | **O** | **2.70** |
| | | | **Nh2** | **8B** | **N** | **3.45** |
| Leu | 265A | CD1 | Trp | 6B | CB | 3.81 |
| | | | Trp | 6B | CE3 | 3.57 |
| Tyr | 268A | CD1 | Trp | 6B | C | 3.75 |
| | | | Trp | 6B | O | 3.30 |
| | | | Trp | 6B | CB | 3.88 |
| Tyr | 268A | CE1 | Trp | 6B | CA | 3.71 |
| | | | Wat | 103B | O | 3.63 |
| | | | Trp | 6B | C | 3.71 |
| | | | Lys | 7B | O | 3.80 |
| | | | Trp | 6B | O | 3.71 |
| Tyr | 268A | CE2 | Trp | 6B | CD1 | 3.56 |
| Tyr | 268A | CZ | Trp | 6B | CD1 | 3.90 |
| Tyr | 268A | OH | Wat | 102B | O | 3.89 |
| | | | **Wat** | **103B** | **O** | **3.47** |
| Met | 281A | O | Nh2 | 8B | N | 3.86 |
| Met | 281A | CE | Nh2 | 8B | N | 3.65 |
| Phe | 284A | CG | Lys | 7B | CB | 3.90 |
| Phe | 284A | CD2 | Lys | 7B | CB | 3.42 |
| Phe | 284A | CE1 | Arg | 5B | O | 3.81 |
| Phe | 284A | CE2 | Lys | 7B | CB | 3.62 |
| | | | His | 3B | O | 3.17 |
| Phe | 284A | CZ | Arg | 5B | O | 3.30 |
| Leu | 288A | CD1 | His | 3B | CB | 3.48 |
| Leu | 288A | CD2 | DNal | 4B | CD1 | 3.71 |
| **water** | **101B** | **O** | **Asp** | **2B** | **OD1** | **2.41** |
| **water** | **102B** | **O** | **Trp** | **6B** | **N** | **3.17** |
| **water** | **103B** | **O** | **Nh2** | **8B** | **O** | **3.23** |
| **water** | **103B** | **O** | **Water** | **102B** | **O** | **2.85** |
| **$Ca^{2+}$** | **2101A** | **CA** | **Asp** | **2B** | **O** | **2.57** |
| $Ca^{2+}$ | 2101A | CA | Asp | 2B | C | 3.78 |
| **$Ca^{2+}$** | **2101A** | **CA** | **DNal** | **4B** | **O** | **2.26** |
| $Ca^{2+}$ | 2101A | CA | DNal | 4B | C | 3.40 |
| $Ca^{2+}$ | 2101A | CA | DNal | 4B | N | 3.80 |

**Table S5. Distances between calcium ion and ligand and MC4R binding partners.** Interactions between the coordinating calcium ion and its binding partners in the three available models of the MC4R binding the peptides NDP-α-MSH, setmelanotide and SHU9119 were analyzed using CONTACT, a program of the CCP4 software suite (Winn et al., 2011). Interactions were calculated between any atoms of the peptide and the receptor, water molecules as well as the calcium ion with a maximum distance of 3.9 Å. Potential hydrogen bonds with a maximum distance of 3.5 Å are highlighted bold. Covalent interactions with $Ca^{2+}$ with a maximum distance of 2.4 Å are highlighted bold and red. D-Nal corresponds to (2R)-2-amino-3-(naphthalene-2-YL)propanoic acid abbreviated as 4J2 in the protein database (PDB entry 6w25). At the second position of the SHU9119 peptide there is a homoserine. In the corresponding pdb file (PDB ID: 6w25), this amino acid is abbreviated as Asp B2. D-phenylalanine is abridged as three letter code Dpn.

| Target atoms | | | NDP-α-MSH | Setmelanotide | SHU9119 |
|---|---|---|---|---|---|
| **Amino acid** | **Number/ Chain** | **Atom** | **Distance [Å]** | **Distance [Å]** | **Distance [Å]** |
| Glu | 100A/R | CD | - | - | 3.37 |
| Glu | 100A/R | OE1 | - | - | **2.21** |
| Glu | 100A/R | OE2 | **3.20** | **2.90/3.02** | **2.21** |
| Asp | 122A/R | CG | - | - | **3.27** |
| Asp | 122A/R | OD1 | - | - | **2.27** |
| Asp | 122A/R | OD2 | **3.09** | **3.35** | 3.65 |
| Asp | 126A/R | CG | 3.50 | 3.34 | 3.04 |
| Asp | 126A/R | OD1 | **3.33** | **2.87** | **2.64** |
| Asp | 126A/R | OD2 | **2.93** | **3.07** | **2.73** |
| Glu | $5P^{-1}$ | O | **2.68** | - | - |
| Glu | $5P^{-1}$ | C | 3.88 | - | - |
| Dpn | $7P^{1}$ | O | **2.50** | - | - |
| Dpn | $7P^{1}$ | C | 3.72 | - | - |
| Dpn | $5P^{1}$ | O | - | **2.64** | - |
| Dpn | $5P^{1}$ | C | - | 3.82 | - |
| His | $4P^{0}$ | N | - | **3.41** | - |
| His | $4P^{0}$ | NE2 | - | **2.50** | - |
| Arg | $1P^{-3}$ | NE | - | **3.44** | - |
| Asp | $2B^{-1}$ | O | - | - | **2.57** |
| Asp | $2B^{-1}$ | C | - | - | 3.78 |
| DNal | $4B^{1}$ | O | - | - | **2.26** |
| DNal | $4B^{1}$ | C | - | - | 3.40 |
| DNal | $4B^{1}$ | N | - | - | 3.80 |

**Table S6. Comparison of the receptor–ligand contacts in complexes of MC4R with the agonists NDP-α-MSH and setmelanotide as well as the antagonist SHU9119.** Interactions of all relevant amino acids of the MC4R the calcium ion ($Ca^{2+}$) and water molecules within the ligand binding pocket towards the binding peptides NDP-α-MSH, setmelanotide and SHU9119 were summarized based on the tables S2-4. Amino acids were abbreviated as one letter code, capital numbers display the position within the receptor or peptide (in the pdb file), and superscripted numbering refers to the conserved *HxRW* motif of the binding peptides. D-Nal corresponds to (2R)-2-amino-3-(naphthalene-2-YL)propanoic acid abbreviated as 4J2 in the protein database (PDB ID: 6w25). D-phenylalanine, D-alanine and norleucine are abridged as three letter code Dpn, Dal and Nle, respectively.

| **MC4R** | **NDP-α-MSH** (agonist) | **Setmelanotide** (agonist) | **SHU9119** (antagonist) |
|---|---|---|---|
| F51 | H6$^{0}$ | H4$^{0}$ | - |
| N97 (L97 - antagonized structure) | - | - | Dnal4$^{1}$ |
| E100 | Nle4$^{-2}$, E5$^{-1}$ | A: Dpn5$^{1}$<br>B: Dal3$^{-1}$ | H3$^{0}$, Dnal4$^{1}$, D2$^{-1}$ |
| T101 | H6$^{0}$ | H4$^{0}$ | H3$^{0}$ |
| V103 | - | - | Nle1$^{-2}$ |
| I104 | Nle4$^{-2}$ | Dal3$^{-1}$ | Nle1$^{-2}$, D2$^{-1}$, H3$^{0}$ |
| T118 | - | - | Nle1$^{-2}$ |
| D122 | Nle4$^{-2}$ | R1$^{-3}$, R6$^{2}$ | Nle1$^{-2}$, D2$^{-1}$, R5$^{2}$ |
| N123 | - | R6$^{2}$ | R5$^{2}$ |
| D126 | Dpn7$^{1}$, R8$^{2}$ | Dpn5$^{1}$, Arg6$^{2}$ | Dnal4$^{1}$, Arg5$^{2}$ |
| I129 | Dpn7$^{1}$ | Dpn5$^{1}$ | Dnal4$^{1}$ |
| C130 | - | - | Dnal4$^{1}$ |
| L133* | - | Dpn5$^{1}$ | Dnal4$^{1}$ |
| I185* | R8$^{2}$ | - | Dnal4$^{1}$, R5$^{2}$ |
| S188 | R8$^{2}$, W9$^{3}$ | R6$^{2}$, W7$^{3}$ | R5$^{2}$, W6$^{3}$ |
| V193 | W9$^{3}$ | - | - |
| I194 | W9$^{3}$ | W7$^{3}$ | W6$^{3}$ |
| L197 | W9$^{3}$ | W7$^{3}$ | Dnal4$^{1}$, W6$^{3}$ |
| F261 | - | Dpn5$^{1}$ | Dnal4$^{1}$ |
| H264 | W9$^{3}$, G10$^{4}$ | W7$^{3}$, C8$^{4}$ | W6$^{3}$, K7$^{4}$, $NH_2$8$^{5}$ |
| L265 | - | - | W6$^{3}$ |
| Y268 | W9$^{3}$, K11$^{5}$, P12$^{6}$ | W7$^{3}$ | W6$^{3}$, K7$^{4}$ |
| M281 | - | - | $NH_2$8$^{5}$ |
| F284 | H6$^{0}$, R8$^{2}$, G10$^{4}$ | H4$^{0}$, R6$^{2}$, C8$^{4}$ | H3$^{0}$, R5$^{2}$, K7$^{4}$ |
| N285 | - | H4$^{0}$ | - |
| L288 | H6$^{0}$, Dpn7$^{1}$ | H4$^{0}$ | H3$^{0}$, Dnal4$^{1}$ |
| **$Ca^{2+}$** | MC4R: E100, D122, D126; Ligand: E5$^{-1}$, Dpn7$^{1}$ | MC4R: E100, D122, D126; Ligand: Dpn5$^{1}$ | MC4R: E100, D122, D126; Ligand: D2$^{-1}$, Dnal4$^{1}$ |
| water 30 | - | Dal3$^{-1}$, H4$^{0}$, C8$^{4}$, Q43, F284, N285 | - |
| water 43 | Nle4$^{-2}$ | - | - |
| water 63 | - | R1$^{-3}$, R6$^{2}$ | - |

| | | | |
|---|---|---|---|
| water 101 | - | - | $D2^{-1}$, $H3^{0}$, $K7^{4}$, |
| water 102 | - | - | $R5^{2}$, $W6^{3}$, Y268, water 103, water 104 |
| water 103 | - | - | $W6^{3}$, $K7^{4}$, Y268, water102, water 105 |
| water 104 | - | - | $Dnal4^{-1}$, $R5^{2}$, $W6^{3}$, $K7^{4}$, water 102, water 105 |
| water 105 | - | - | $D2^{-1}$, $K7^{4}$, water 103, water 104 |

**Table S7. Determination of total and cell surface expression with Nano-Glo®HiBiT Lytic/Extracellular detection system, and maximal cAMP response $E_{max}$ via AlphaScreen®.** Data are given as the result of four to eight independent experiments performed in triplicates ± SEM. Wild-type MC4R (WT) stimulation as fold over MC4R basal for all assays is 20 ± 1.06 fold for α-MSH and 22 ± 1.15 fold for NDP-α-MSH, both set as 100. Expression data were cleaned by performing a ROUT test with Q = 1%. Statistics were done by one-way ANOVA with Kruskal-Wallis test. WT was tested against all mutants stimulated with the indicated ligand: a: $p < 0.05$; b: $p < 0.01$; c: $p < 0.001$; d: $p < 0.0001$.

| Substitution | Total expression [fold over WT basal] | Cell surface expression [fold over WT basal] | Basal [fold over WT basal] | α-MSH $E_{max}$ [% of WT at 1 µM] | NDP-α-MSH $E_{max}$ [% of WT at 1 µM] |
|---|---|---|---|---|---|
| MC4R WT | 1 | 1 | 1 | 100 | 100 |
| E100N | 1.30 ± 0.10 | 2.42 ± 0.15[d] | 0.97 ± 0.12 | 11.35 ± 1.45[a] | 56.10 ± 4.13[a] |
| T101A | 1.41 ± 0.07 | 0.89 ± 0.11 | 1.16 ± 0.16[a] | 189 ± 10[d] | 155 ± 12[b] |
| D122S | 1.63 ± 0.06[c] | 1.41 ± 0.09 | 1.07 ± 0.08 | 40.10 ± 3.46 | 129 ± 8.56 |
| N123A | 0.53 ± 0.03[d] | 0.60 ± 0.05[c] | 1.03 ± 0.07 | 163 ± 13[b] | 119 ± 7.89 |
| D126S | 1.28 ± 0.07 | 1.87 ± 0.14[c] | 0.88 ± 0.07[a] | 13.92 ± 1.67[a] | 91.23 ± 7.74 |
| C130A | 0.99 ± 0.04 | 1.13 ± 0.09 | 1.03 ± 0.09 | 147 ± 9.24[a] | 114 ± 9.25 |
| L133A | 1.11 ± 0.10 | 0.78 ± 0.15 | 1.03 ± 0.05 | 130 ± 6.33 | 152± 8.29[a] |
| L133F | 0.97 ± 0.09 | 1.34 ± 0.11 | 0.85 ± 0.04[a] | 112 ± 6.17 | 137± 6.74 |
| I137A | 0.83 ± 0.04 | 0.79 ± 0.03 | 1.30 ± 0.08 | 92.78 ± 4.96 | 99.40 ± 4.89 |
| I137F | 0.81 ± 0.07 | 0.57 ± 0.06[b] | 1.02 ± 0.06 | 76.65 ± 3.93 | 94.21 ± 5.53 |
| T150A | 0.60 ± 0.06[c] | 0.58 ± 0.04[b] | 0.88 ± 0.08[c] | 26.11. ± 1.64[d] | 18.07 ± 2.17[d] |
| T150I | 0.69 ± 0.06[a] | 0.55 ± 0.02[b] | 0.81 ± 0.06[c] | 36.46 ± 2.24[d] | 40.74 ± 2.08[d] |
| T150D | 0.41 ± 0.03[d] | 0.47 ± 0.03[c] | 0.85 ± 0.05 | 9.88 ± 0.91[d] | 10.85 ± 0.86[c] |
| T150F | 1.09 ± 0.05 | 0.88 ± 0.03 | 0.77 ± 0.05[a] | 42.60 ± 2.41[d] | 55.68 ± 4.64[a] |
| T150S | 1.44 ± 0.09 | 1.23 ± 0.05 | 0.62 ± 0.04[c] | 77.81 ± 5.11[a] | 86.18 ± 5.14 |
| H158A | 1.35 ± 0.17 | 1.37 ± 0.16 | 2.87 ± 0.23[d] | 129 ± 15 | 129 ± 15 |
| F184V | 1.14 ± 0.07 | 1.82 ± 0.14[c] | 1.24 ± 0.11 | 149 ± 14 | 91.23 ± 8.29 |
| S188A | 1.10 ± 0.13 | 1.15 ± 0.06 | 0.94 ± 0.07 | 148 ± 11[a] | 142 ± 15 |
| D189S | 0.98 ± 0.06 | 1.51 ± 0.13 | 0.80 ± 0.06 | 151 ± 16 | 166 ± 28 |
| S191A | 1.00 ± 0.06 | 0.89 ± 0.04 | 1.00 ± 0.04 | 199 ± 18[c] | 197 ± 22[c] |
| L197A | 0.78 ± 0.06 | 1.12 ± 0.16 | 0.84 ± 0.05 | 152 ± 11 | 171 ± 24[a] |
| M204A | 0.60 ± 0.04[c] | 0.69 ± 0.04 | 1.81 ± 0.13[d] | 113.04 ± 4.51 | 103 ± 5.33 |
| L205A | 0.92 ± 0.06 | 0.82 ± 0.08 | 1.21 ± 0.06 | 101.39 ± 6.65 | 99.85 ± 5.61 |
| L205F | 1.14 ± 0.05 | 1.02 ± 0.04 | 2.32 ± 0.20[d] | 111.57 ± 6.40 | 112 ± 6.27 |
| F254A | 1.73 ± 0.15[a] | 0.94 ± 0.04 | 1.02 ± 0.03 | 88.67 ± 5.41 | 85.09 ± 5.23 |
| F254M | 1.99 ± 0.17[b] | 1.57 ± 0.10[a] | 0.99 ± 0.04 | 77.33 ± 8.14 | 97.12 ± 10.32 |
| W258A | 0.73 ± 0.07 | 0.48 ± 0.04[c] | 0.94 ± 0.03 | 61.40 ± 4.13[d] | 65.04 ± 5.30[b] |
| W258F | 1.30 ± 0.09 | 0.96 ± 0.06 | 2.48 ± 0.22[d] | 78.13 ± 4.39[a] | 72.66 ± 4.83 |
| F261V | 0.81 ± 0.04 | 0.77 ± 0.10 | 0.92 ± 0.10 | 63.54 ± 4.19 | 61.61 ± 5.96 |
| H264A | 0.47 ± 0.04[d] | 1.35 ± 0.05 | 1.18 ± 0.09 | 10.21 ± 0.94[a] | 133 ± 9.10 |
| L265A | 0.62 ± 0.04[d] | 0.78 ± 0.05 | 0.89 ± 0.08 | 83.68 ± 6.16 | 140 ± 12 |
| Y268F | 0.98 ± 0.07 | 1.06 ± 0.06 | 0.87 ± 0.03 | 140 ± 16 | 133 ± 14 |
| H283A | 0.95 ± 0.07 | 1.17 ± 0.08 | 1.40 ± 0.18 | 164 ± 15[a] | 152 ± 21 |
| F284V | 0.67 ± 0.09[b] | 0.90 ± 0.05 | 1.08 ± 0.03 | 145 ± 14 | 165 ± 27 |
| N285S | 0.86 ± 0.07 | 0.99 ± 0.07 | 1.14 ± 0.05 | 173 ± 16[a] | 167 ± 22 |

| | | | | | |
|---|---|---|---|---|---|
| Y287V | 0.74 ± 0.02 | 1.75 ± 0.10[b] | 0.99 ± 0.04 | 148 ± 12 | 142 ± 18 |
| L288A | 0.51 ± 0.04[d] | 0.47 ± 0.05[d] | 0.94 ± 0.04 | 76.41 ± 5.51 | 129 ± 12 |
| I291A | 0.49 ± 0.03[d] | 0.54 ± 0.05[b] | 0.94 ± 0.03 | 11.09 ± 0.94[d] | 8.98 ± 0.69[d] |
| I291F | 0.81 ± 0.08 | 0.54 ± 0.04[c] | 0.92 ± 0.05 | 79.83 ± 6.47 | 71.34 ± 6.02 |

**Table S8. Determination of half maximal effective concentration $EC_{50}$ via cAMP accumulation using the AlphaScreen® assay after addition of agonists α-MSH or NDP-α-MSH.** Data are given as the result of four to eight independent experiments performed in triplicates ± SEM. Statistics were done by one-way ANOVA with Kruskal-Wallis test. Wild-type MC4R (WT) was tested against all mutants stimulated with the indicated ligand. a: $p < 0.05$; b: $p < 0.01$; c: $p < 0.001$; d: $p < 0.0001$; n.d. = not determined due to either too low $E_{max}$ or due to severely shifted concentration-response curve which do not allow proper $EC_{50}$ calculation.

| Substitution | α-MSH $EC_{50}$ [nM] | NDP-α-MSH $EC_{50}$ [nM] |
|---|---|---|
| MC4R WT | 17.6 ± 4.21 | 1.17 ± 0.26 |
| E100N | n.d. | 383 ± 139[c] |
| T101A | 49.4 ± 6.12 | 0.9 ± 0.05 |
| D122S | n.d. | 9.74 ± 1.61[b] |
| N123A | 33.3 ± 5.97 | 0.93 ± 0.23 |
| D126S | n.d. | 166 ± 34.2[c] |
| C130A | 136 ± 21.9[c] | 2.20 ± 0.71 |
| L133A | 94.7 ± 1.58 | 2.07 ± 0.26 |
| L133F | 76.1 ± 17.0 | 3.79 ± 0.77 |
| I137A | 19.7 ± 5.97 | 1.15 ± 0.08 |
| T150A | n.d. | n.d. |
| T150I | 86.2 ± 34.9 | 1.41 ± 0.35 |
| T150F | 91.9 ± 16.9 | 1.52 ± 0.33 |
| T150S | 29.3 ± 7.65 | 0.81 ± 0.19 |
| H158A | 7.59 ± 3.83 | 0.58 ± 0.14 |
| S188A | 101 ± 30.2 | 0.97 ± 0.19 |
| M204A | 6.34 ± 1.17 | 1.06 ± 0.44 |
| L205F | 7.18 ± 1.84 | 1.38 ± 0.45 |
| W258A | 70.7 ± 17.7 | 2.53 ± 0.34 |
| W258F | 2.60 ± 0.53 | 0.59 ± 0.2 |
| F261V | 118 ± 14.5[c] | 2.96 ± 1.44 |
| H264A | n.d. | 179 ± 32.5[d] |

#### REFERENCES SUPPLEMENTAL INFORMATION


Adams, P.D., Afonine, P.V., Bunkoczi, G., Chen, V.B., Davis, I.W., Echols, N., Headd, J.J., Hung, L.W., Kapral, G.J., Grosse-Kunstleve, R.W., *et al.* (2010). PHENIX: a comprehensive Python-based system for macromolecular structure solution. Acta Crystallogr D Biol Crystallogr *66*, 213-221.

Afonine, P.V., Poon, B.K., Read, R.J., Sobolev, O.V., Terwilliger, T.C., Urzhumtsev, A., and Adams, P.D. (2018). Real-space refinement in PHENIX for cryo-EM and crystallography. Acta Crystallogr D Struct Biol *74*, 531-544.

Ballesteros, J.A., and Weinstein, H. (1995). Integrated methods for the construction of three-dimensional models and computational probing of structure-function relationships in G-protein coupled receptors. Methods Neurosci *25*, 366-428.

Berman, H.M., Westbrook, J., Feng, Z., Gilliland, G., Bhat, T.N., Weissig, H., Shindyalov, I.N., and Bourne, P.E. (2000). The Protein Data Bank. Nucleic Acids Res *28*, 235-242.

Biebermann, H., Castaneda, T.R., van Landeghem, F., von Deimling, A., Escher, F., Brabant, G., Hebebrand, J., Hinney, A., Tschop, M.H., Gruters, A., *et al.* (2006). A role for beta-melanocyte-stimulating hormone in human body-weight regulation. Cell Metab *3*, 141-146.

Chen, V.B., Arendall, W.B., 3rd, Headd, J.J., Keedy, D.A., Immormino, R.M., Kapral, G.J., Murray, L.W., Richardson, J.S., and Richardson, D.C. (2010). MolProbity: all-atom structure validation for macromolecular crystallography. Acta Crystallogr D Biol Crystallogr *66*, 12-21.

Chrencik, J.E., Roth, C.B., Terakado, M., Kurata, H., Omi, R., Kihara, Y., Warshaviak, D., Nakade, S., Asmar-Rovira, G., Mileni, M., *et al.* (2015). Crystal Structure of Antagonist Bound Human Lysophosphatidic Acid Receptor 1. Cell *161*, 1633-1643.

Collaborative Computational Project, N. (1994). The CCP4 suite: programs for protein crystallography. Acta Crystallogr D Biol Crystallogr *50*, 760-763.

Emsley, P., Lohkamp, B., Scott, W.G., and Cowtan, K. (2010). Features and development of Coot. Acta Crystallogr D Biol Crystallogr *66*, 486-501.

Hall, T.A. (1999). BioEdit: a user-friendly biological sequence alignment editor and analysis program for Windows 95/98/NT. Nucleic Acids Symposium Series *Series 41*, 95-98.

Hanson, M.A., Roth, C.B., Jo, E., Griffith, M.T., Scott, F.L., Reinhart, G., Desale, H., Clemons, B., Cahalan, S.M., Schuerer, S.C., *et al.* (2012). Crystal structure of a lipid G protein-coupled receptor. Science *335*, 851-855.

Kang, Y., Kuybeda, O., de Waal, P.W., Mukherjee, S., Van Eps, N., Dutka, P., Zhou, X.E., Bartesaghi, A., Erramilli, S., Morizumi, T., *et al.* (2018). Cryo-EM structure of human rhodopsin bound to an inhibitory G protein. Nature *558*, 553-558.

Kim, K., Che, T., Panova, O., DiBerto, J.F., Lyu, J., Krumm, B.E., Wacker, D., Robertson, M.J., Seven, A.B., Nichols, D.E., *et al.* (2020). Structure of a Hallucinogen-Activated Gq-Coupled 5-HT2A Serotonin Receptor. Cell *182*, 1574-1588 e1519.

Laskowski, R.A., and Swindells, M.B. (2011). LigPlot+: multiple ligand-protein interaction diagrams for drug discovery. J Chem Inf Model *51*, 2778-2786.

Liu, K., Wu, L., Yuan, S., Wu, M., Xu, Y., Sun, Q., Li, S., Zhao, S., Hua, T., and Liu, Z.J. (2020). Structural basis of CXC chemokine receptor 2 activation and signalling. Nature *585*, 135-140.

McDonald, I.K., and Thornton, J.M. (1994). Satisfying hydrogen bonding potential in proteins. J Mol Biol *238*, 777-793.

Okada, T., Sugihara, M., Bondar, A.N., Elstner, M., Entel, P., and Buss, V. (2004). The retinal conformation and its environment in rhodopsin in light of a new 2.2 A crystal structure. J Mol Biol *342*, 571-583.

Paisdzior, S., Dimitriou, I.M., Schope, P.C., Annibale, P., Scheerer, P., Krude, H., Lohse, M.J., Biebermann, H., and Kuhnen, P. (2020). Differential Signaling Profiles of MC4R Mutations with Three Different Ligands. Int J Mol Sci *21*.

Pettersen, E.F., Goddard, T.D., Huang, C.C., Couch, G.S., Greenblatt, D.M., Meng, E.C., and Ferrin, T.E. (2004). UCSF Chimera--a visualization system for exploratory research and analysis. J Comput Chem *25*, 1605-1612.

Punjani, A., Zhang, H., and Fleet, D.J. (2020). Non-uniform refinement: adaptive regularization improves single-particle cryo-EM reconstruction. Nat Methods *17*, 1214-1221.

Rasmussen, S.G., DeVree, B.T., Zou, Y., Kruse, A.C., Chung, K.Y., Kobilka, T.S., Thian, F.S., Chae, P.S., Pardon, E., Calinski, D., *et al.* (2011). Crystal structure of the beta2 adrenergic receptor-Gs protein complex. Nature *477*, 549-555.

Schwefel, D., Groom, H.C., Boucherit, V.C., Christodoulou, E., Walker, P.A., Stoye, J.P., Bishop, K.N., and Taylor, I.A. (2014). Structural basis of lentiviral subversion of a cellular protein degradation pathway. Nature *505*, 234-238.

Stoddart, L.A., Johnstone, E.K.M., Wheal, A.J., Goulding, J., Robers, M.B., Machleidt, T., Wood, K.V., Hill, S.J., and Pfleger, K.D.G. (2015). Application of BRET to monitor ligand binding to GPCRs. Nat Methods *12*, 661-663.

Vagin, A.A., Steiner, R.A., Lebedev, A.A., Potterton, L., McNicholas, S., Long, F., and Murshudov, G.N. (2004). REFMAC5 dictionary: organization of prior chemical knowledge and guidelines for its use. Acta Crystallogr D Biol Crystallogr *60*, 2184-2195.

Vaguine, A.A., Richelle, J., and Wodak, S.J. (1999). SFCHECK: a unified set of procedures for evaluating the quality of macromolecular structure-factor data and their agreement with the atomic model. Acta Crystallogr D Biol Crystallogr *55*, 191-205.

Winn, M.D., Ballard, C.C., Cowtan, K.D., Dodson, E.J., Emsley, P., Evans, P.R., Keegan, R.M., Krissinel, E.B., Leslie, A.G., McCoy, A., *et al.* (2011). Overview of the CCP4 suite and current developments. Acta Crystallogr D Biol Crystallogr *67*, 235-242.

Yin, J., Chen, K.M., Clark, M.J., Hijazi, M., Kumari, P., Bai, X.C., Sunahara, R.K., Barth, P., and Rosenbaum, D.M. (2020). Structure of a D2 dopamine receptor-G-protein complex in a lipid membrane. Nature *584*, 125-129.

Yu, J., Gimenez, L.E., Hernandez, C.C., Wu, Y., Wein, A.H., Han, G.W., McClary, K., Mittal, S.R., Burdsall, K., Stauch, B., *et al.* (2020). Determination of the melanocortin-4 receptor structure identifies Ca(2+) as a cofactor for ligand binding. Science *368*, 428-433.

Zhao, Y., Chapman, D.A., and Jones, I.M. (2003). Improving baculovirus recombination. Nucleic Acids Res *31*, E6-6.
